# *N*-arylpyrazole NOD2 agonists promote immune checkpoint inhibitor therapy

**DOI:** 10.1101/2023.01.26.525573

**Authors:** Matthew E. Griffin, Taku Tsukidate, Howard C. Hang

## Abstract

The characterization of microbiota mechanisms in health and disease has reinvigorated pattern recognition receptors as prominent targets for immunotherapy. Notably, our recent studies on *Enterococcus* species revealed peptidoglycan remodeling and activation of NOD2 as key mechanisms for microbiota enhancement of immune checkpoint inhibitor therapy. Inspired by this work and other studies of NOD2 activation, we performed *in silico* ligand screening and developed *N*-arylpyrazole dipeptides as novel NOD2 agonists. Importantly, our *N*-arylpyrazole NOD2 agonist is enantiomer-specific, effective at promoting immune checkpoint inhibitor therapy and requires NOD2 for activity *in vivo*. Given the significant functions of NOD2 in innate and adaptive immunity, these next-generation agonists afford new therapeutic leads and adjuvants for a variety of NOD2-responsive diseases.

## MAIN

Pattern recognition receptors (PRRs) provide important sensors for host recognition of microbes as well as environmental and cellular stresses to initiate innate and adaptive immunity responses.^1^ These immune sensors have therefore emerged as excellent therapeutic targets for modulation of infection and inflammatory diseases in animals and humans. For example, the discovery of toll-like receptors (TLRs) has been transformative for understanding elucidating mechanisms of innate immunity and enabled the development several TLR-targeted therapeutics in clinical trials.^2^ Beyond these PRRs, nucleotide-binding domain, leucine-rich repeat receptors (NLRs) also encode intracellular sensors of microbial ligands and stress.^3^ Amongst the NLRs, the founding members of this protein superfamily, nucleotide-binding and oligomerization domain (NOD) proteins 1 and 2 have been recognized key protein targets for immune activation.^4, 5^ Notably, one of the key active components of the bacterial cell wall in Freund’s adjuvant,^6^ muramyl dipeptide (MDP) is the minimal ligand for required the activation of NOD2 and downstream pro-inflammatory NF-κB signaling^7–9^. In contrast, NOD1 recognizes γ-d-glutamyl-*meso*-diaminopimelic acid (iE-DAP) an alternative peptidoglycan fragment of bacterial cell walls.^10, 11^ These two intracellular NLRs serve as important intracellular sensors of peptidoglycan fragments that is conserved in most bacterial species and crucial for host defense against diverse pathogens, as *Nod1* and *Nod2*-deficient mice are more susceptible to bacterial and viral infections.^12^ In addition to pathogens, these immune sensors of peptidoglycan also play key roles for detection of microbiota and maintenance of mucosal barriers. Notably, loss-of-function alleles of NOD2 are associated with compromised intestinal barrier function, microbiota dysbiosis and are major risk factors for inflammatory bowel disease (IBD) such as Crohn’s disease.^12^ Alternatively, gain-of-function NOD2 mutants are implicated in auto-inflammatory disease such as Blau syndrome.^12^ Interestingly, NOD1 and NOD2 have also been implicated in sensing endoplasmic reticulum (ER) stress,^13^ potentially through binding of sphingosine-1-phosphate,^14^ and are also associated with inflammatory disorders such as obesity and neurodegeneration^12^. Furthermore, NOD2 expression in hypothalamic inhibitory neurons has been shown to regulate appetite and body temperature in mice, which suggests additional roles for NOD2 in metabolism and behavior.^15^ These intracellular NLRs are therefore at the nexus of health and disease and prominent targets for therapeutic development.^16, 17^

Based on the significance of NOD1 and NOD2 in health and disease, their peptidoglycan ligands have been extensively derivatized to generate more effective synthetic compounds for therapeutic applications. Early studies on iE-DAP and MDP suggests these NOD1 and NOD2 agonists could stimulate neutrophils and monocytes, respectively, and facilitate the clearance of pathogens as well as tumor cells *ex vivo* and *in vivo*, but their modest potency and short half-lives limited their efficacy in animal models.^17^ Several studies were then initiated to improve the activity of these peptidoglycan fragments. In particular, the addition of lipophilic groups onto iE-DAP and MDP have improved their potency in cells and half-lives *in vivo*. For example, the addition of N-lauroyl fatty acid onto iE-DAP resulted in C12-iE-DAP, which exhibits ~100-fold greater activation of NOD1-dependent NF-κB reporter activation.^17^ Similar approaches towards MDP have also yielded more potent NOD2 agonists. Notably, the addition of fatty acyl chains onto the 6-OH of MurNAc and/or esterification of amino acids afforded more potent MDP-based NOD2 agonists.^17^ While these and other improved NOD1/2 agonists exhibited greater immunostimulatory activity, their efficacy as anti-infectives or adjuvants *in vivo* were limited. However, the development of muramyl tripeptide-phosphatidylethanolamine (MTP-PE) or mifamurtide into a liposomal formulation resulted in the drug Mepact,^18^ which has been approved to treat pediatric osteosarcoma in Europe. While Mepact was not approved by the FDA in USA, the collective clinical studies indicate NOD2 activation was beneficial for the preventing tumor metastases.^18^ Of note, microbiome profiling of cancer patients suggested immune checkpoint inhibitor therapy-responsive individuals have unique microbiota composition, which included several species of *Enterococcus*.^19^ Our subsequent studies demonstrated the *Enterococcus* species that express the unique secreted peptidoglycan hydrolase (SagA) can remodel peptidoglycan and activate NOD2 to promote immune checkpoint inhibitor cancer therapy in mice.^20^ Moreover, the co-administration of MDP as the free molecule^20^ or encapsulated in nanoparticles^21^ with immune checkpoint inhibitors resulted in enhanced tumor clearance in mice, suggesting improved NOD2 agonist formulation and/or composition may result in more effective immune checkpoint inhibitor therapy and treatment of other disease indications. Indeed, efforts to generate more effective NOD2 agonists are ongoing through the development of desmuramyl dipeptide (dMDP) analogs, which replace the potentially labile and dynamic structure of MurNAc with more stable and functional chemotypes.^22^ For example, lipophilic cinnamic acid-based dipeptide analogs have been developed as dMDP NOD2 agonists *ex vivo*^23, 24^ and are effective as adjuvants for promoting antigen-specific immune responses in mice^25^. Inspired by our SagA-expressing *Enterococcus* peptidoglycan remodeling and MDP co-administration studies with immune checkpoint inhibitor therapy,^20^ we explored new desmuramyl dipeptide NOD2 agonists by structure-based *in silico* screening and developed *N*-arylpyrazole dipeptides as novel NOD2 agonists, which are specific and effective at promoting immune checkpoint inhibitor therapy *in vivo*. These next-generation agonists should function as new therapeutic leads and adjuvants for a variety of NOD2-responsive diseases.

## RESULTS

### Identification of *N*-arylpyrazole dipeptide NOD2 agonists by structure-based *in silico* screening

Structure-based *in silico* screening is a powerful approach to generate new ligand scaffolds.^26^ To enable *in silico* screening for NOD2 agonists, whose ligand-bound structure remains unknown, we docked MDP onto the putative ligand-binding leucine-rich repeat (LRR) domain of NOD2 with the induced-fit approach^27^ (**Fig. 1A**). This model shows MDP makes contacts with amino acid residues that are critical for receptor activation.^28–30^ To further validate the model, we performed template-based docking of active and inactive MDP derivatives and dMDPs reported in literature by aligning the maximum common substructures with the bound MDP (**Fig. 1B**). This resulted in the enrichment of the active derivatives in generated models. While the template-based approach was less sensitive for dMDPs, perhaps indicative of the weak or moderate affinity of nearly all reported dMDPs compared to MDP, it remained highly selective for our screening purpose. With the template-based docking method in hand, we virtually screened a library of ~31,000 dMDPs (**Fig. 1C**). We constructed this *in silico* library by conjugating enamine carboxylic acid building blocks to the dipeptide L-Val-D-Glu, which improves the activity of dMDPs relative to the native L-Ala-D-Glu and filtered the candidate ligands based on their physicochemical properties. Template docking resulted in 1,683 poses, and the top-scoring 10% were clustered and manually inspected. As further validation of our *in silico* method, our screening returned multiple, structurally distinct hits. In addition, some hits resembled previously reported dMDPs, including cinnamoyl-Gly-L-Val-D-Glu(OEt)2 or CinGVE^23^. A subset of the top 10% virtual screening hits encompassing the chemical diversity of the hits were synthesized and tested for activity with the NOD2-HEK-Blue NF-κB reporter cells (**Fig. 1D**). Six of the initial nine compounds effectively activated NOD2 with similar efficacy as CinGVE (**Fig. 1E**). Interestingly, one of the *N*-arylpyrazole compounds **TT007** was active whereas the other two *N*-arylpyrazole compounds **TT008** and **TT009** were inactive, which suggested a structure–activity relationship around the pyrazole ring. Importantly, **TT007** was inactive in the NULL and NOD1 HEK-Blue NF-κB reporter cells (**Extended Data 1**), which confirmed its selectivity for NOD2, and elicited a clear dose-dependent NOD2 activation with pEC_50_ ~6.0 (**Figs. 1F**). Encouraged by these results, we subsequently focused our effort on the *N*-arylpyrazole scaffold.

**Fig. 1.**
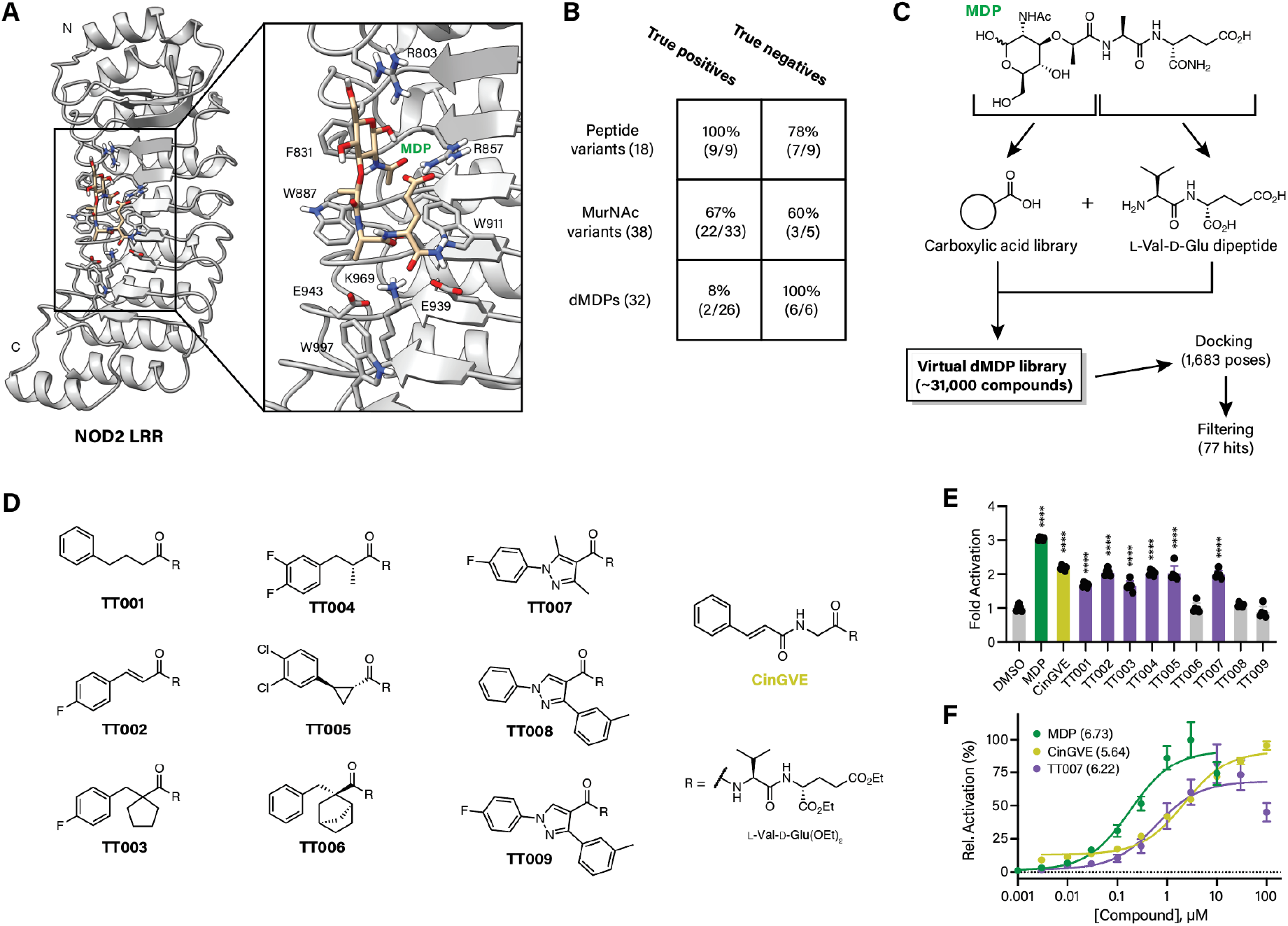
Virtual screening for small molecule NOD2 agonists. **a**, Docked structure of muramyl dipeptide (MDP) in the leucine-rich repeat (LRR) domain of rabbit NOD2 (pdb 5IRM). Inset shows amino acids within 5 angstroms of the docked MDP molecule. **b**, Results of virtual docking approach with reported chemical structures related to MDP. **c**, Workflow of virtual screening in which a library was generated using carboxylic acids appended to the L-Val-D-Glu dipeptide and then docked to the NOD2 LRR domain. **d**, Desmuramyl MDP (dMDP) structures resulting from the virtual screen that were synthesized for further screening. Chemicals shown as a pure enantiomer were synthesized as a racemic mixture. **e**, Activation of NOD2-expressing HEK-293T cells treated for 16 h with 10 μM of the indicated compound using a colorimetric assay. **f**, Dose-dependent activation of NOD2-expressing HEK-293T cells using the indicated compounds. Numbers in parentheses represent calculated negative log of the half maximal effective concentration (pEC50) values. NOD activation assays were analyzed using a one-way ANOVA with Dunnett’s multiple comparisons test compared to the DMSO control. pEC50 values were derived by a three-parameter dose-response nonlinear regression. **** *P* < 0.0001.

### Optimization of *N*-arylpyrazole dipeptide NOD2 agonists

We explored the chemical space of the *N*-arylpyrazole scaffold in two parts: (1) pyrazole group and (2) *N*-aryl group (**Fig. 2A** and **Extended Data 2A**). The dipeptide was synthesized as previously described and conjugated to *N*-arylpyrazole scaffolds that were either purchased or synthesized via Knorr pyrazole synthesis or Chan-Evans-Lam coupling routes (**Extended Data 2B-D** and **Supplementary Materials**). As the initial screening results suggested that substitutions on the pyrazole ring have a large effect on potency, we designed analogs of the three *N*-arylpyrazole compounds with different substitutions (**Fig. 2B**). Removing either one of the methyl groups from **TT007** as in **TT010** and **TT011** reduced potency by up to two orders of magnitude. Increasing the size of a substituent as in **TT009** and **TT012** did not compensate the lack of the other substituent. This trend was also observed with compounds with various *N*-aryl group as in **TT013**-**TT017** (**Extended Data 2E**). Methyl groups can have profound effects on the binding affinity of small molecule ligands.^31^ One possible explanation for our observation may be that the two methyl groups stabilize the binding conformation by creating an energy barrier for the rotation around the bond between the aryl and pyrazole groups or the bond between the pyrazole and the amide groups. In addition, *N*-methylation on Val (**TT018** vs. **TT019**, **TT010** vs. **TT020**, **TT011** vs. **TT021**, **Fig. 2C**), which was introduced to perturb the dihedral angle between the pyrazole and amide groups, abolished the activity regardless of substituents on the pyrazole ring. We then designed analogues of **TT007** with various *N*-aryl groups. Topliss tree approach^32^ as in **TT018**, **TT022**, **TT023**, and **TT024** did not yield compounds with appreciable differences in potency except for a slight increase in **TT022** possibly due to the increased steric bulk (**Fig. 2D**). We therefore introduced heteroaromatic rings that could increase steric bulk and potentially participate in polar interactions. Substitutions with a hydrogen bond donor group on the *para* or *meta* position relative to the pyrazole ring as in **TT025**-**TT030** increased potency up to ~10-fold (**Fig. 2E**). On the other hand, replacing the fluorophenyl group with a hydrogen bond acceptor group such as pyridine, pyrimidine, or anisole as in **TT031**-**TT036** did not improve potency relative to **TT007 (Fig. 2F**). Because MDP activity can be improved by lipid acylation^17^, we wanted to ensure that the increased activity of our compounds was not due to a concomitant increase in lipophilicity. Comparison of the *N*-arylpyrazole compounds showed a moderately higher potency from the initial hit while also decreasing compound lipophilicity (**Fig. 2G**), with **TT030** exhibiting the highest overall lipophilic ligand efficiency (LLE) of 6.81. Together, these structure–activity relationship studies led to improved potency for the *N*-arylpyrazole hit guided by the modification of tolerant portions of the pharmacophore while also maintaining low lipophilicity.

**Fig. 2.**
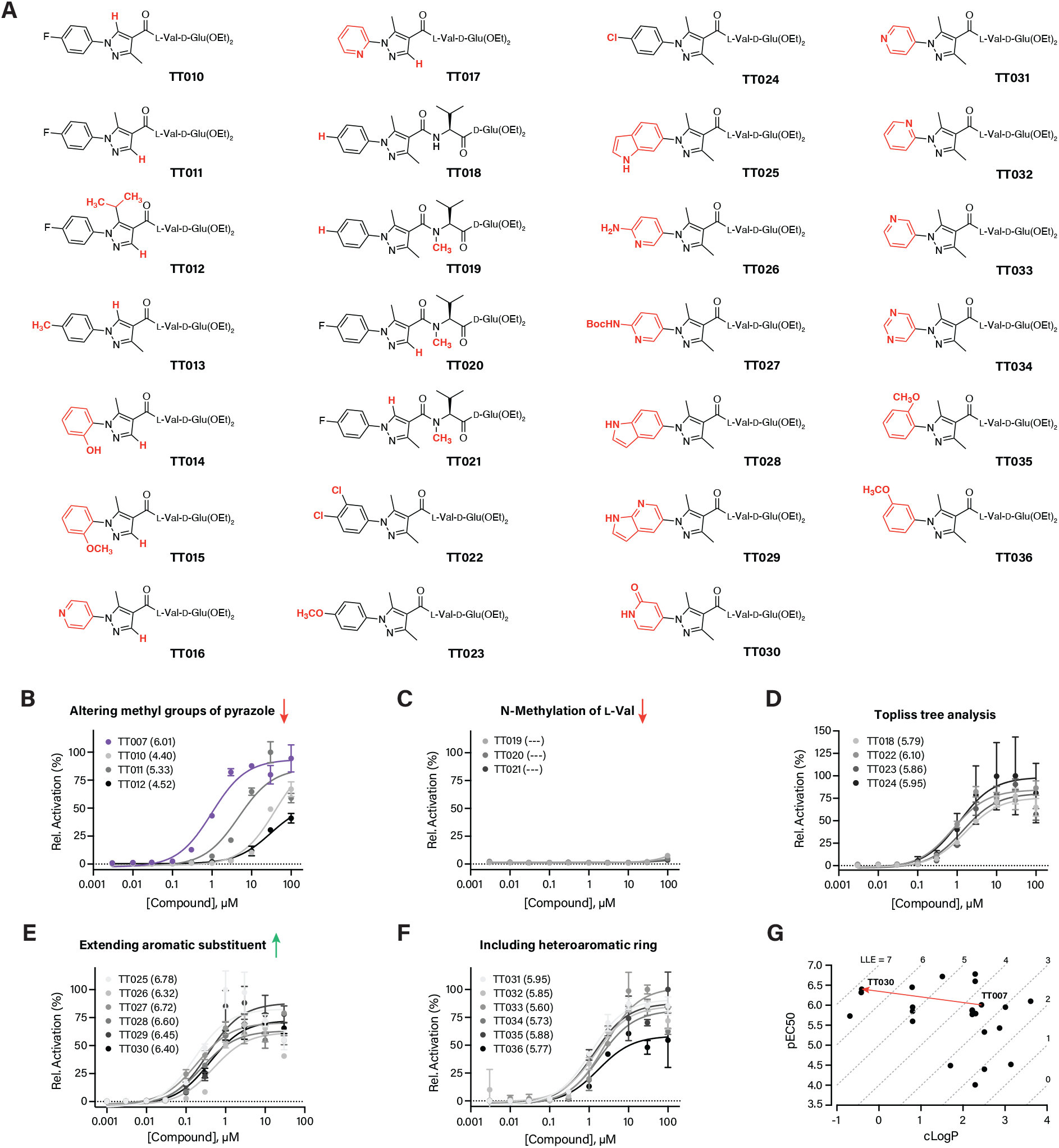
Structure-activity relationship studies of 3,5-dimethyl-*N*-arylpyrazole NOD2 agonists. **a**, Structures of 3,5-dimethyl-*N*-arylpyrazole structures synthesized for SAR studies. **b-f**, Dose-dependent activation of NOD2-expressing HEK-293T cells using the indicated compounds grouped by similar structural perturbations. Numbers in parentheses represent calculated pEC50 values. **g**, Graph of calculated pEC50 values vs. calculated partition coefficient (clogP). Dotted lines show equivalent lipophilic ligand efficiency (LLE) values. Red arrow indicates optimization of the **TT007** scaffold to **TT030**. pEC50 values were derived by a three-parameter dose-response nonlinear regression.

### *N*-arylpyrazole NOD2 agonist promotes cytokine production *ex vivo*

To screen NOD2 activation in a more physiologically relevant system, we examined the cytokine release profile of the optimized lead **TT030** using primary human immune cells. Healthy human peripheral mononuclear blood cells (PBMCs) were first depleted of red blood cells and then stimulated overnight with vehicle, MDP, or **TT030**. Because bacterial lipopolysaccharide has been reported to amplify cytokine secretion caused by NOD2 stimulation, we also conducted the experiment in parallel with LPS co-stimulation. As expected, MDP stimulation led to a significant increase of canonical, NOD2-dependent cytokines in the medium after 24 h, including interleukin-1 beta (IL-1β) (**Fig. 3A**) and tumor necrosis factor alpha (TNFα) (**Fig. 3B**). Similarly, **TT030** was able to induce equivalent concentrations of these cytokines. In addition, **TT030** caused an increase in IL-12p70 secretion with and without LPS (**Fig. 3C**), as well as the production of the IL-1 family member IL-18 (**Extended Fig. 3A**) and the IL-12 family member IL-23 (**Extended Fig. 3B**), highlighting the broad impact of NOD2 activation on the overall cytokine environment. Importantly, IL-12 family cytokines lead to the production of interferon gamma (IFNɣ) by activated T and NK cells to enhance adaptive immune responses, and increased IFNɣ levels were also observed for both MDP and TT0124964 stimulation (**Fig. 3D**).

**Fig. 3.**
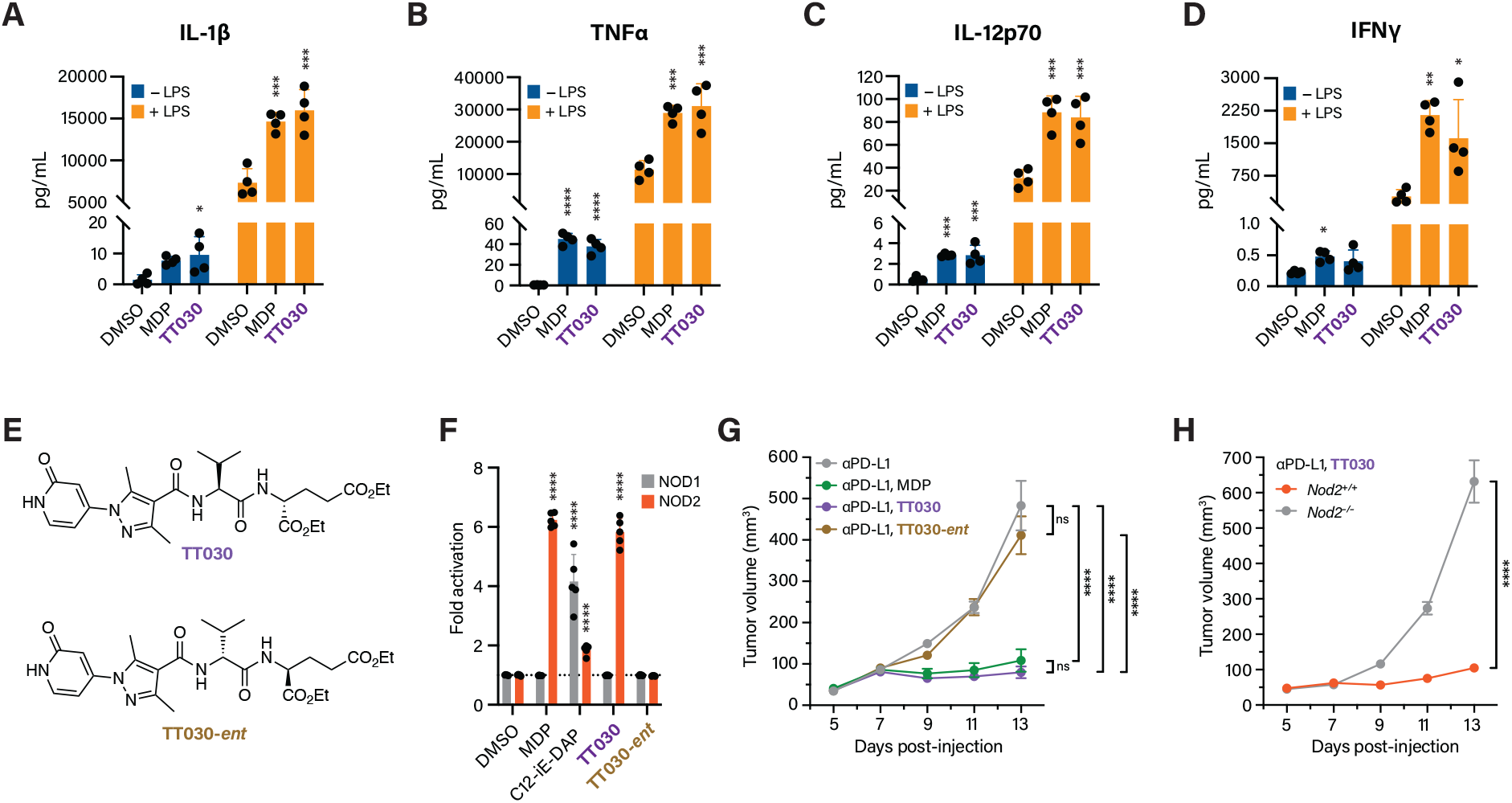
NOD2 agonist TT030 activates cytokine production in vitro and improves immune checkpoint inhibitor treatment of tumor growth in vivo. **a-d**, Measurements of secreted cytokine levels from human peripheral mononuclear blood cells (PMBCs) treated with the indicated compounds for 24 h without or with lipopolysaccharide (LPS). **e**, Structures of **TT030** and its enantiomer (**TT030-*ent***). **f**, Activation of NOD1- and NOD2-expressing HEK-293T cells with the indicated compounds. C12-iE-DAP is a lipophilic version of the NOD1 agonist, iE-DAP. **g**, Growth profiles of B16-F10 subcutaneous tumors in wild-type C57BL/6 mice treated with 100 μg α-PD-L1 and 100 μg of the indicated compounds every other day starting at Day 7. **h**, Growth profiles of B16-F10 subcutaneous tumors in wild-type or *Nod2*^-/-^ C57BL/6 mice treated with 100 μg α-PD-L1 and 100 μg **TT030** every other day starting at Day 7. Cytokine secretion assays and NOD activation assays were analyzed using a one-way ANOVA with Dunnett’s multiple comparisons test compared to the DMSO control. Longitudinal tumor growth assays were analyzed using linear mixed-effects model analysis on the log transformed data. ns = not significant, * *P* < 0.05, ** *P* < 0.01, *** *P* < 0.001, **** *P* < 0.0001.

### *N*-arylpyrazole NOD2 agonist promotes immunotherapy *in vivo*

We next compared **TT030** activity against MDP *in vivo*. Our previous work had demonstrated that MDP can potentiate the activity of anti–PD-L1 checkpoint inhibitor treatment of a subcutaneous B16-F10 melanoma model in a NOD2-dependent manner.^20^ Using a similar treatment model, mice inoculated with B16 tumors were co-administered with anti–PD-L1 and either MDP or **TT030**. In addition, the enantiomer of **TT030** (**TT030-*ent***) was chemically synthesized (**Fig. 3E**) and shown to be NOD2-inactive using the HEK-Blue *in vitro* assay (**Fig. 3F**). As a control compound, **TT030-*ent*** was administered to a separate cohort of tumor-bearing animals along with the checkpoint inhibitor. As anticipated, administration of anti–PD-L1 alone was not effective at controlling B16 tumor growth over time, whereas co-treatment with MDP significantly inhibited tumor growth (**Fig. 3G**). This anti-tumor activity was also observed for the **TT030** but not for **TT030-*ent***, indicating the *N*-arylpyrazole NOD2 agonist was active *in vivo* and enantiomer-specific. We then repeated this experiment in wild-type and *Nod2*^-/-^ mice (**Fig. 3H**). NOD2 knockout caused a loss in the anti-tumor activity observed in the wild-type cohort, demonstrating that NOD2 was required for the ability of **TT030** to augment anti–PD-L1 immunotherapy *in vivo*.

### *N*-arylpyrazole NOD2 agonist enhances intratumoral immune responses *in vivo*

We then further characterized the anti-tumor mechanism of action of the *N*-arylpyrazole NOD2 agonist *in vivo*. Infiltrating leukocytes from tumors treated with anti–PD-L1 and either **TT030** or **TT030-*ent*** were harvested via cell sorting and subjected to single cell transcriptomic profiling (**Fig. 4A** and **Extended Data 4**). After assignment of cell clusters using well-defined markers, we examined the overall composition of intratumoral immune cells by comparing the relative proportions of each cell cluster. Here, we found that tumors treated with TT0124964 showed a significant increase in the amount of a T cell cluster characterized by the expression of *Cd8a, Ifng, Gzmb*, and *Mki67* (**Figs. 4B, 4C** and **Extended Data 4 and 5A**), suggesting that the **TT030** sample contained a higher amount of dividing cytotoxic T cells. In fact, overall expression of both *Ifng* and *Gzmb* were elevated in the **TT030** sample compared to the enantiomer control (**Extended Data 5B**), indicating heighted CD8^+^ T cell activation and effector function. We also observed shifts in multiple clusters of mononuclear phagocytes (monocytes and macrophages) (**Figs. 4B, 4C**), which indicated that **TT030** stimulation could also remodel the intratumoral myeloid compartment. To better characterize changes to downstream signaling caused by **TT030**, we performed differential gene expression and gene set enrichment analyses (**Supplementary Tables S1-S15**). Numerous inflammatory, immune activation, and metabolic pathways were upregulated in multiple clusters from tumors treated with **TT030**, including IFNα and IFNɣ response as well as NF-κB signaling (**Fig. 4D**). As mentioned above, NOD2 initiates transcriptional changes through the transcription factor NF-κB, which we observed with broad enrichment in REACTOME pathways annotated for NF-κB activation (**Fig. 4E**). We also found enrichment in REACTOME pathways annotated for signaling via IL-1β (**Fig. 4E**), indicating that **TT030** activated the canonical NOD2–NF-κB–IL-1β signaling axis within the tumor microenvironment akin to our results *ex vivo* and our previous studies with MDP *in vivo*^20^.

**Fig. 4.**
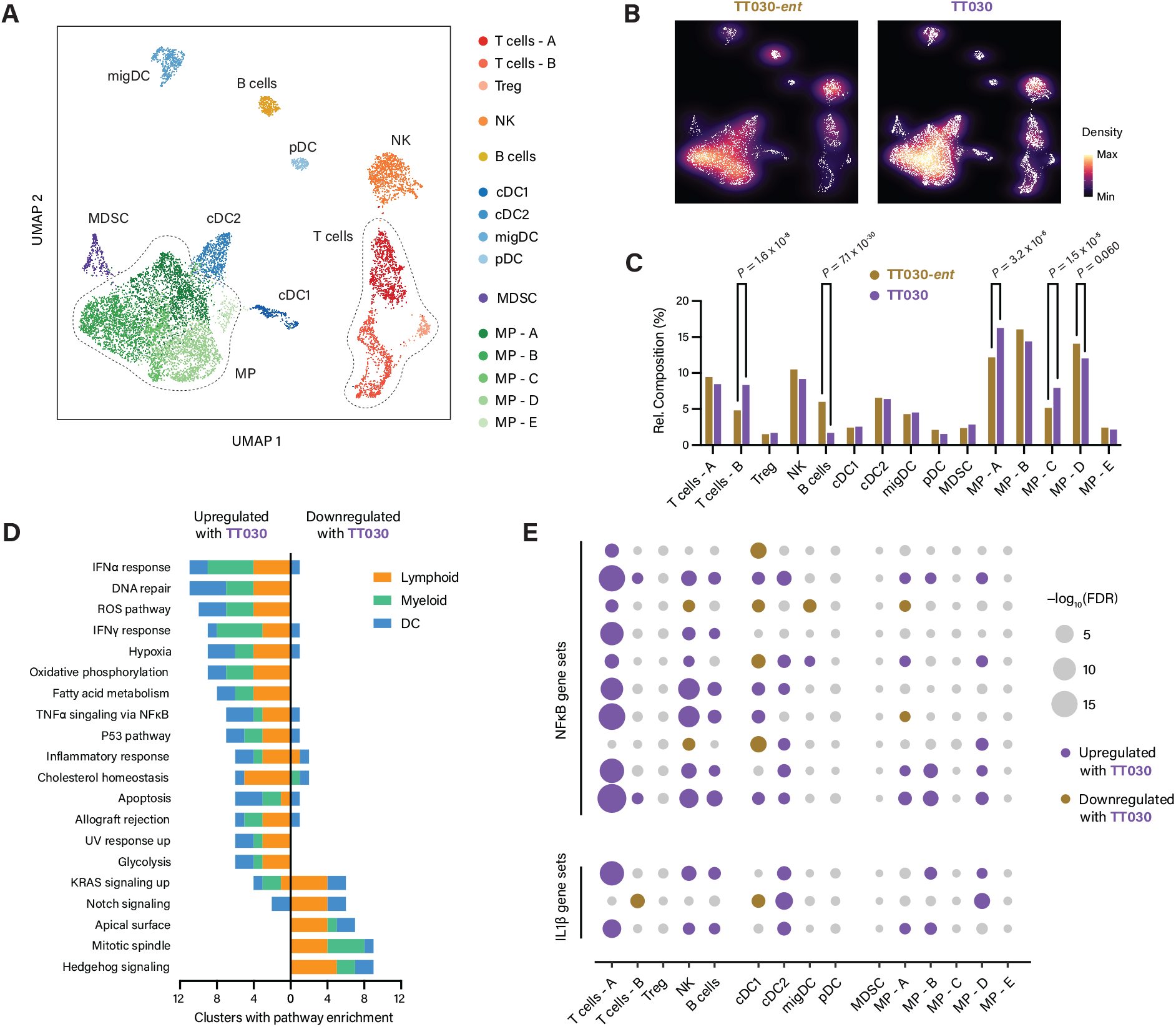
NOD2 agonist TT030 remodels intratumoral immune cell composition and elicits broad inflammatory signaling. **a**, Uniform manifold approximation and projection (UMAP) reduction of single intratumoral immune cells harvested from α-PD-L1 and **TT030**- and **TT030-*ent***-treated tumors. **b-c**, Heatmap and quantification of the relative composition of intratumoral immune cell types divided by clusters. **d**, Quantification of the number of clusters that were enriched for individual Molecular Signature Database (MSigDB) hallmark gene sets. **e**, Bubble plot for enrichment of curated REACTOME gene sets involving NF-κB or interleukin 1 across cell clusters. For relative composition, absolute cell counts were analyzed by Pearson’s chisquare test for count data with Holm’s correction for multiple comparisons; exact *P* values are given. False discovery rates (FDRs) for gene set enrichment were obtained using modelbased analysis of single transcriptomics (MAST). NOD activation assays were analyzed using a one-way ANOVA with Dunnett’s multiple comparisons test compared to the DMSO control.

## DISCUSSION

PRRs represent an attractive class of drug targets due to their broad involvement in inflammation and immune regulation. Nevertheless, many synthetic approaches to design and improve PRR agonists rely on the addition of lipophilic moieties such as lipid anchors or their incorporation into novel formulations such as liposomes and other nanoparticles. Here, we describe the development of a novel class of NOD2 agonists built upon the 3,5-dimethyl-*N*-arylpyrazole head group. This moiety has several advantages, including facile synthetic routes to access the scaffold and commercial availability of diverse precursors for structure–activity relationship studies. These factors enabled the generation and screening of multiple analogs, ultimately providing lead compounds with nanomolar EC_50_ values *in vitro*. Importantly, we found that the *N*-arylpyrazole group played a critical role in NOD2 binding, with multiple desmethyl analogs showing significantly decreased potency. The two methyl groups may potentially play an important role in orienting the planarity of the *N*-arylpyrazole group with respect to the neighboring amide bond. Further conformational studies of the molecules or structural analyses of NOD2 in the ligand-bound state may provide additional insights into the binding mode of this pharmacophore.

As expected, our results indicate that the *N*-arylpyrazole NOD2 agonist **TT030** functions similarly to MDP, the naturally occurring molecular pattern for NOD2. Both MDP and **TT030** elicit a variety of cytokines that mediate innate and adaptive immune responses, including IL-1β, TNFα, and IL-12p70. **TT030** also exhibited equivalent anti-tumor activity as MDP in combination with anti–PD-L1. We have previously demonstrated that MDP can synergize with antibodies against all clinically approved checkpoint targets; therefore, the *N*-arylpyrazole NOD2 agonist may also function as an adjuvant for multiple checkpoint inhibitors and will be examined in future studies. A recent report has indicated that phosphorylation of MDP at the 6-O-hydroxyl position by *N*-acetylglucosamine kinase (NAGK) is required for NOD2 activation using *Nagk*^-/-^ mammalian cells *ex vivo*.^33^ Our results, along with other reports of desmuramyl NOD2 agonists that lack a potential phosphorylation site^22–25^, suggest that synthetic agonists can engage NOD2 without further modification, and additional biophysical or structural studies may help to illuminate potentially multiple binding poses of NOD2 agonists. Importantly, the activity observed in our system required both the proper stereochemistry of the *N*-arylpyrazole dipeptide as well as the expression of host NOD2, providing further evidence for its predicted, on-target activity. Finally, alterations in intratumoral immune cell composition and activation mirror those observed with MDP stimulation, including increased levels of CD8^+^ effector T cells and activation of NF-κB and IL-1β signaling. Altogether, our results support TT0124964 and the 3,5-dimethyl-*N*-arylpyrazole scaffold as a potent lead compound and chemotype to stimulate NOD2 signaling for eliciting both innate and adaptive immune responses *in vivo* for immunotherapy.

## METHODS

### General synthetic procedures

All chemicals and solvents were purchased from Acros Organics, Alfa Aesar, Ambeed, Enamine, Sigma-Aldrich, TCI America and others and used without further purification.

#### HATU-mediated coupling

The corresponding carboxylic acid derivative (1.0–1.1 equiv.) and HATU (1.0–1.1 equiv.) were weighed in a scintillation vial and dissolved in DCM or DMF (0.2 M). The vial was flashed with argon gas and cooled to 0 °C. To this was added DIPEA (5.0 equiv.), and the reaction mixture was stirred for 15 min. Then, a solution of the corresponding dipeptide (deprotected by acidolysis with 25 % TFA/DCM or 4N HCl/1,4-dioxane) (1.0 equiv.) in DCM or DMF (0.2 M) was added, and the reaction mixture was allowed to warm to RT and stirred for 24–72 h. The reaction mixture was directly purified with flash chromatography when DCM was used as solvent or first concentrated by half before loading onto column when DMF was used as solvent. Alternatively, the reaction mixture was diluted with 70 % EtOAc/hexanes, washed with aqueous NH_4_Cl and brine, dried over MgSO_4_, filtered, and concentrated before loading column. Flash chromatography was typically performed with a gradient of EtOAc/hexanes or MeOH/DCM.

#### Alkaline hydrolysis

The corresponding ester was weighed in a scintillation vial and dissolved in ethanol. The vial was flashed with argon gas. To a solution of ethyl ester in ethanol, was added 2 M LiOH (final concentration of LiOH: 1 M), and the reaction mixture was stirred at 50 °C for 24 h. Upon completion of the reaction, as monitored by TLC, the aqueous phase was acidified with 1 N HCl to pH ≈ 3 and extracted with EtOAc. The organic layer was dried over MgSO_4_, filtered, and concentrated. Alternatively, the reaction mixture was acidified with Amberlite resin, filtered, and concentrated. The residue was dried under vacuum at 50 °C.

### Cell culture

The following cell lines were used and sourced as follows: HEK-Blue NOD2 (InvivoGen, hkb-hnod2), HEK-Blue NOD1 (InvivoGen, hkb-nod1), HEK-Blue Null2 (hkb-null2), and B16-F10 (ATCC CRL-6475). All cell lines were cultured at 37 °C in 5% CO_2_ in DMEM (ThermoFisher, 11995065) supplemented with 10% fetal bovine serum (FBS), 100 U mL^−1^ penicillin, and 100 μg mL^−1^ streptomycin. Cells were maintained at no greater than 70% confluency and subcultured using TrypLE Express. Cell lines were routinely cultured without antibiotics to ensure no bacterial infection and tested for mycoplasma using a Universal Mycoplasma Detection Kit (ATCC 30-1012K). Primary human peripheral mononuclear blood cells (PBMCs) were cultured at 37 °C in 5% CO_2_ in RPMI 1640 (ThermoFisher, 61870143) supplemented with 10% FBS. All experimental work using human samples was reviewed and approved by the Scripps Research Institutional Review Board (Protocol IRB-21-7816NOD2).

### Animals

Specific pathogen-free, eight-week old male C57BL/6J (B6, 000664) and B6.129S1-*Nod2^tm1Flv^*/J (*Nod2*^-/-^ 005763 from Flavell Lab, Yale School of Medicine) were obtained from The Jackson Laboratory. Animals were housed in autoclaved caging with SaniChip bedding and enrichment for nest building on a 12-hour light/dark cycle. Animals were provided gamma-irradiated chow (LabDiet, 5053) and sterile drinking water ad libitum. Animal care and experiments were conducted in accordance with NIH guidelines and approved by the Institutional Animal Care and Use Committee at Scripps Research (Protocol AUP-21-095).

### Preparation of NOD2/LRR–MDP docking model

All computational studies were carried out using the Schrödinger Drug Discovery Platform and RDKit. Structures and NOD2 agonist activity data of MDP and its analogues and derivatives were curated from literature. These ligands were prepared for docking using LigPrep to test the performance of docking models. MDP conformers were generated using an expanded sampling protocol in MacroModel.^34^ The leucine-rich repeat (LRR) domain of NOD2 was excised from the crystal structure of *Oc*NOD2 (PDB ID: 5IRN) and prepared for docking using Protein Preparation Wizard. A receptor-based pharmacophore model was generated by E-Pharmacophore. In Phase, this pharmacophore model was used to screen the MDP conformers to generate initial docking models. These docking models were then refined using Glide SP with vdW scaling and clustered, and 100 representative models were selected. For each of the 100 models, the side chains within 5 Å of MDP were minimised using Prime MM-GB/SA to account for local receptor flexibility. Resulting receptors were clustered, and 13 representative models were selected for redocking. In each receptor of the ensemble was docked MDP using Glide SP-PEP. These docking models were then refined and rescored using Prime MM-GB/SA, and 10 top-scoring models were selected for further evaluation. To test if any of these models can correctly tell active ligands from inactive ligands, the set of known MDP analogues and derivatives was docked using Glide SP while their maximum common structures with MDP were fixed in position. For desmuramyl dipeptides with aromatic surrogates, an additional positional constraint for an aromatic atom was inferred from receptor-based pharmacophore models and applied in Glide XP. The best-performing model was selected based on the balance of sensitivity and selectivity and visually inspected.

### DMDP library preparation and screening

Chemical structures of Enamine Carboxylic Acid Building Blocks were downloaded as SDF (07/18/2020). Using an RDKit script, these carboxylic acid derivatives (31344 compounds) were conjugated with *L*-Val-*D*-Glu, and resulting DMDPs were filtered against PAINS and required to have at least one aromatic ring and to reside within the beyond-rule-of-five space boundaries. The filtered ligands were prepared for docking using LigPrep and docked using a cascade of Glide SP (passed: 3938 compounds) and XP (passed: 1683 compounds) while their maximum common structures with MDP were fixed in position. Resulting models were refined and rescored using Prime MM-GB/SA, and the top 10% (168 ligands) were selected for further analysis. Filtering out ligands that considerably deviated from their original docking poses during the refinement (RMSD for MCS with MDP > 1.5 Å) and tautomers left 77 unique compounds, which were clustered and visually inspected.

### Reporter cell line activation assay

Approximately 24 h before the assay, culture medium for each of the HEK-Blue reporter cell lines was replaced with a 1:1 mixture of DMEM supplemented with 10% FBS and OptiMEM. On the day of the assay, single cell suspensions (150,000 cells/mL) were produced in HEK-Blue detection medium (InvivoGen, hb-det2). Cells were aliquoted at 200 uL per well of a 96-well plate containing 0.2 μL of a 1000x stock of each test compound in DMSO. Cells were then incubated at 37 °C in 5% CO_2_ for 8–10 h. To measure activity, wells were then gently pipetted up and down to mix the conditioned medium, and absorbance from the colorimetric product of the secreted alkaline phosphatase was measured at 630 nm. In some assays, cells were stimulated for 16 h with compounds as described above in complete DMEM. Then, 20 μL of the conditioned medium was mixed with 180 μL QUANTI-Blue solution (InvivoGen, rep-qbs) and incubated for 2-4 h at 37 °C. Activity was then measured by absorbance as described above.

### Cytokine release assay

Human PMBCs were obtained from whole blood donated by healthy individuals via the Scripps Normal Blood Donor Service. Whole blood was diluted 1:1 with phosphate-buffered saline (PBS) containing 2% FBS and then layered onto Lymphoprep (STEMCELL Technologies, 07851) in SepMate-50 isolation tubes (STEMCELL Technologies, 85480). The discontinuous gradient was centrifuged for 10 min at 1,200 x *g*, and mononuclear cells were removed by pouring off. Mononuclear cells were then resuspended in PBS + 2% FBS and centrifuged for 3 min at 200 x *g* twice before further use. Washed mononuclear cells were then aliquoted at 5 × 10^5^ cells per well of a 48-well plate with 300 μL RPMI containing 10% FBS (complete RPMI). Cells were then diluted with 100 μL complete RPMI containing 50 μM of the test compound (10 μM final concentration). Then, 100 μL complete RPMI (–LPS) or 100 μL complete RPMI containing 5 ng mL^−1^ LPS (+LPS, 10 ng mL^−1^ final concentration) was added. Cells were incubated overnight at 37 °C in 5% CO_2_, and conditioned medium was harvested and centrifuged for 5 min at 500 × *g* to remove residual cells. Samples were used directly for cytokine analysis with the LEGENDPlex Human Inflammatory Panel 1 (BioLegend, 740809) according to the manufacturer’s protocol. For analysis, 25 μL of the conditioned medium was used for –LPS samples, and 12.5 μL was used for the +LPS samples. On-bead ELISA readouts were measured using the standard PE and APC detection filter parameters on a ZE5 flow cytometer (Bio-Rad; 405, 488, 561, and 640 lasers) in the Scripps Research Flow Cytometry Core Facility. Raw data files were then imported into the LEGENDPlex Cloud Data Analysis software suite (BioLegend) and analyzed using the manufacturer’s pre-set parameters.

### Tumor growth assay

Tumor growth assays were carried out as previously described.^20^ B16-F10 cells were subcultured on the day prior to harvesting for tumor inoculation. On the day of injection, cells were harvested using TrypLE Express, washed and resuspended in cold PBS, and counted with a hemocytometer. Cells were then resuspended at 2 × 10^6^ cells mL^−1^ in PBS and mixed 1:1 with Matrigel matrix (Corning; growth factor reduced, 356231). Cell suspensions were kept on ice prior to injection. Animals were anesthetized using 3% isoflurane, and their right flank was shaved with hair clippers. Animals were then subcutaneously injected with 100 μL of the cell suspension on their mid-right flank (1 × 10^5^ cells per injection). Once tumors were established at roughly 25-100 mm^3^, tumor volume was measured every other day. Digital calipers were used to measure the length and width of each tumor, and tumor volume was calculated as length x width x 0.5, where width was the smaller of the two measurements. Checkpoint inhibitor treatment was started two days following the first measurement and continued every other day for three total injections. For each injection, 100 μg anti-PD-L1 (BioXCell, BP0101) and 100 μg of either MDP (InvivoGen, tlrl-mdp), **TT030**, or **TT030-*ent*** were intraperitoneally injected using 200 μL antibody diluent (BioXCell, IP0065). Animals were humanely euthanized with CO_2_ asphyxiation at the end of each experiment.

### Single cell RNA sequencing

Sample preparation and analysis were performed as previously described.^20^ Tumors treated with either **TT030** or **TT030-*ent*** (*n* = 8 tumors per condition) were dissected the day following the third anti-PD-L1 treatment, minced with scissors, and resuspended in 5 mL of RPMI. Each suspension was then supplemented with 0.1 mL of RPMI containing 1.25 mg mL^−1^ Liberase TM (Roche, 5401119001) and 20 mg mL^−1^ DNase I (Roche, 11284932001) and then incubated at 37 °C for 30 min with gentle shaking (80 rpm). Digested tumor tissue was then passed through a 70 μm filter using the blunt end of a syringe plunger, and cells were then pelleted by centrifugation for 2 min at 300 x *g*. Cell pellets were then resuspended in 5 mL of red blood cell lysis buffer (ThermoFisher, 00-4333-57) and incubated for 5 min at RT. Lysis was halted by addition of 20 mL of RPMI, and cells were centrifuged as described above, washed with PBS + 2% FBS, and centrifuged again. Cells were then transferred to flow cluster tubes, centrifuged, and resuspended in 50 μL of residual liquid after aspiration of the PBS wash. To each tube, 1 μL FcBlockX was added, and samples were incubated for 10 min on ice. Cells were then diluted with 50 μL PBS + 2% FBS and FITC anti-CD45 (BioLegend, 304005; 1:1,000 final dilution) and incubated for 20 min on ice in the dark. Cells were washed twice with PBS + 2% FBS and then resuspended in 400 μL PBS + 2% FBS containing DAPI (Millipore Sigma, D9542; 1 mg mL^−1^ stock solution, 1:50,000 final dilution). From each sample, 20,000 CD45^+^DAPI^−^ live tumor-infiltrating leukocytes were isolated by using a MoFlo Astrios EQ jet-in-air sorting flow cytometer (Beckman Coulter; 355, 405, 488, 561, and 640 lasers) in the Scripps Research Flow Cytometry Core Facility. Sorted cells from each condition were pooled, and 10,000 cells per condition were targeted for droplet formation using a Chromium Single Cell System (10x Genomics) in the Scripps Research Genomics Core Facility. Samples were processes according to the manufacturer’s protocol (10x Genomics, Chromium Next Single Cell 3’ GEM Library and Gel Bead Kit, PN-1000121). Libraries were then sequenced on an Illumina NovaSeq 6000 sequencer with a NovaSeq SP flow cell. Pre-processing of the sequencing results were conducted using the 10x Genomics Cell Ranger pipeline (v6.0.0) with default settings, and analysis was performed in R (v4.2.1) using Seurat (v4.2.0).^35, 36^ Datasets were converted into Seurat objects, and ambient RNA was removed by DecontX.^37^ Single particles were filtered using the following cut-offs: mitochondrial RNA < 10%, number of genes > 500, number of UMIs > 1,000, and log(genes UMI^−1^) > 0.75. Seurat objects were then normalized, integrated, and combined. Preliminary clustering was used to identify B16-F10 cells by expression of *Pmel* and *Mlana*, which were removed from downstream analyses. Additionally, clusters containing empty droplets with few to no differentially expressed genes were excluded from further analysis. Data processing of the **TT030** and **TT030-*ent*** samples yielded an expression matrix of 10,082 cells x 19,991 genes. Principle component analysis (PCA) was performed, and the top 18 PCs, which cumulatively contained >90% of the relative standard deviation and <5% incremental increase, were used to cluster the data by FindNeighbors and FindClusters with a resolution of 0.5. Uniform manifold approximation and projection (UMAP) was used for two-dimensional representation of the data. Clusters were assigned identities based on markers described in **Extended Fig. 4**. Differential gene expression analysis and single-cell gene set enrichment analysis (scGSEA) between samples were performed with MAST (Bioconductor v3.15).^38^ Genes with false discovery rates (FDRs) less than 0.01 were identified as differentially expressed between groups. Positive *z* scores represent enrichment in the **TT030** sample and vice versa. For scGSEA, hallmark, curated (C2:CP), and biological process gene ontology (C5:GOBP) gene sets were fetched from the Molecular Signatures Database (MSigDB, v7.5).^39^ Gene sets with a combined adjusted *P* value less than 0.25 were identified as significantly enriched, and the sign of the *z* score was used to denote sample enrichment as described above. Visualizations and quantile-quantile plots of selected genes from enriched pathways were generated by R.

### Statistical analyses

Unless otherwise noted, non-sequence data were analyzed with Prism 8.4.0 (GraphPad). Sequence-based data were analyzed with R as described in the appropriate methods section. Single concentration activation assays or cytokine release assays were analyzed using a one-way ANOVA with Dunnett’s multiple comparisons test compared to the relevant DMSO control. Dose-dependent response assays were analyzed by fitting to a three-parameter nonlinear regression with the Hill coefficient set to 1. Tumor growth assays were analyzed using a mixed-effects analysis on the log pre-processed tumor volumes with Tukey’s multiple comparisons post hoc test (https://github.com/kroemerlab/TumGrowth). *P* < 0.05 was considered statistically significant for all assays, and individual *P* values are denoted as follows: \**P* < 0.05, \*\**P* < 0.01, \*\*\**P* < 0.001, \*\*\*\**P* < 0.0001.

## DATA AVAILABILITY

All scRNA-seq sequencing data are available via the NCBI (BioProject number pending assignment). All other data supporting the findings of this study are available within the article and its supplementary information files and from the corresponding author on reasonable request.

## ACKNOWLEDGEMENTS

This work was supported by the National Institutes of Health (1R01CA245292-01, H.C.H.). M.E.G. is a Hope Funds for Cancer Research Fellow supported by the Hope Funds for Cancer Research (HCFR-19-03-02) and is also funded by a career development award from the Melanoma Research Foundation. T.T. thanks the Tri-Institutional Program in Chemical Biology at The Rockefeller University and Takenaka Scholarship.

## AUTHOR INFORMATION

**Department of Immunology and Microbiology, Scripps Research, La Jolla, CA 92037.**

Matthew E. Griffin, Howard C. Hang

**Department of Chemistry, University of California, Irvine, Irvine, CA 92697.**

Matthew E. Griffin

**Laboratory of Chemical Biology and Microbial Pathogenesis, The Rockefeller University, New York, NY 10065.**

Taku Tsukidate

**Department of Chemistry, Scripps Research, La Jolla, CA 92037.**

Howard C. Hang

### Author contributions

Conceptualization: T.T., M.E.G. and H.C.H.; Methodology: T.T., M.E.G. and H.C.H.; Investigation: M.E.G. and T.T.; Formal analysis: T.T., M.E.G. and H.C.H.; Supervision: H.C.H.; Writing – original draft: M.E.G., T.T and H.C.H.; Writing – review and editing: M.E.G., T.T and H.C.H.

### Corresponding author

Correspondence to Howard C. Hang.

## ETHICAL DECLARATIONS

M.E.G., T.T. and H.C.H. have filed a patent application for the commercial use of *N*-arylpyrazole NOD2 agonists for immunotherapy.

---

**Extended Fig. 1.**
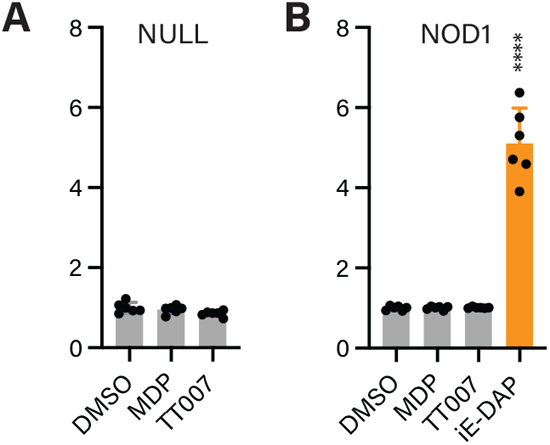
Lead 3,5-dimethyl-*N*-arylpyrazole TT007 specifically activates NOD2. **a-b**, Activation of Null (control) and NOD1-expressing HEK-293T cells treated for 16 h with 10 μM of the indicated compound using a colorimetric assay. Activation assays were analyzed using a one-way ANOVA with Dunnett’s multiple comparisons test compared to the DMSO control. **** *P* < 0.0001.

**Extended Fig. 2.**
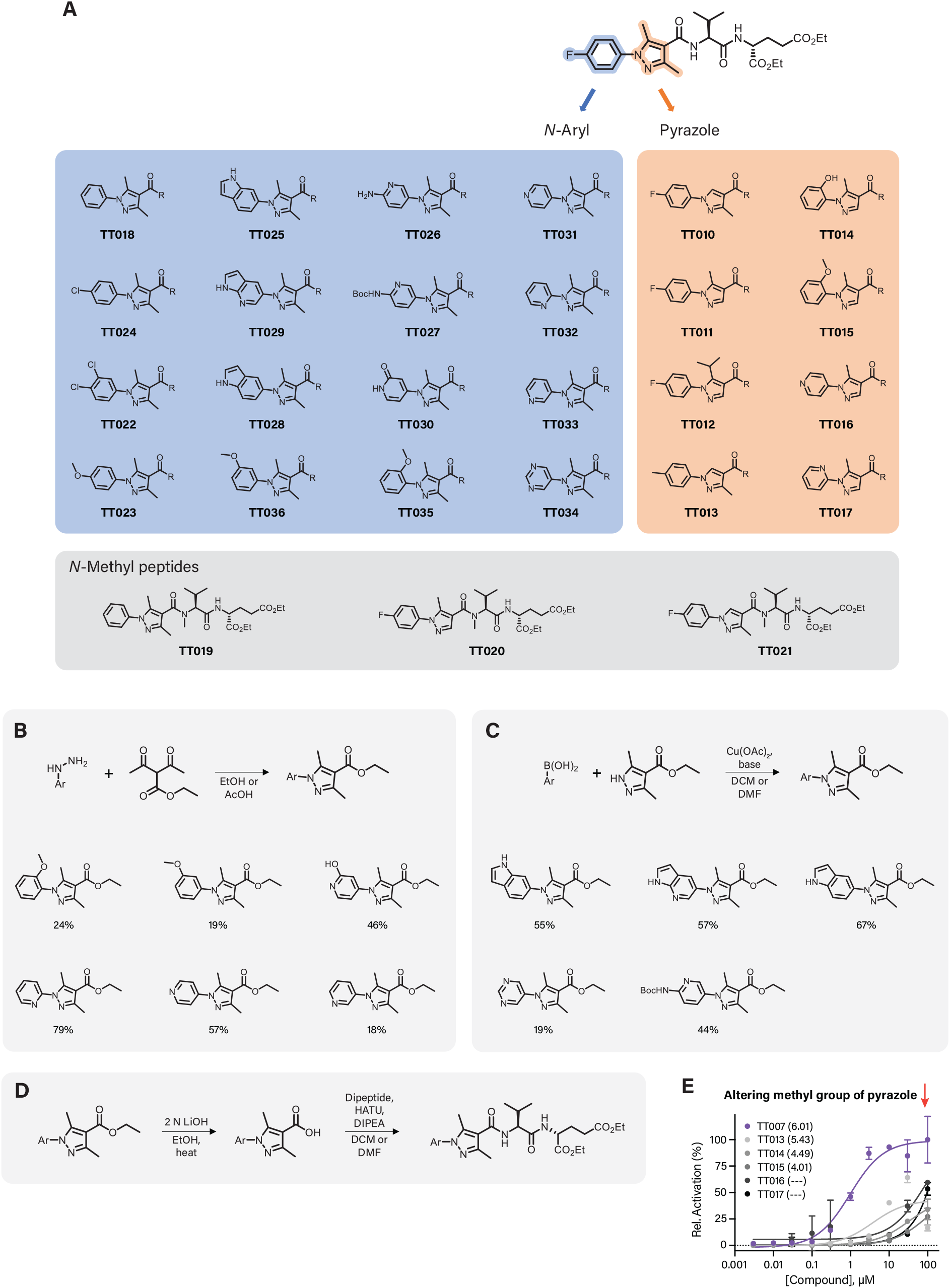
Approach and synthesis of 3,5-dimethyl-*N*-arylpyrazole compound library. **a**, Diversification approach for compound library focusing on the *N*-aryl or pyrazole ring. **b**, Sample of library precursors generated with the Knorr pyrazole method. Numbers represent chemical yield. **c**, Sample of library precursors generated with the Chan-Evans-Lam method. Numbers represent chemical yield. **d**, Deprotection and coupling steps to produce library compounds. **e**, Dose-dependent activation of NOD2-expressing HEK-293T cells using the indicated compounds. Numbers in parentheses represent calculated pEC50 values. pEC50 values were derived by a three-parameter dose-response nonlinear regression.

**Extended Fig. 3.**
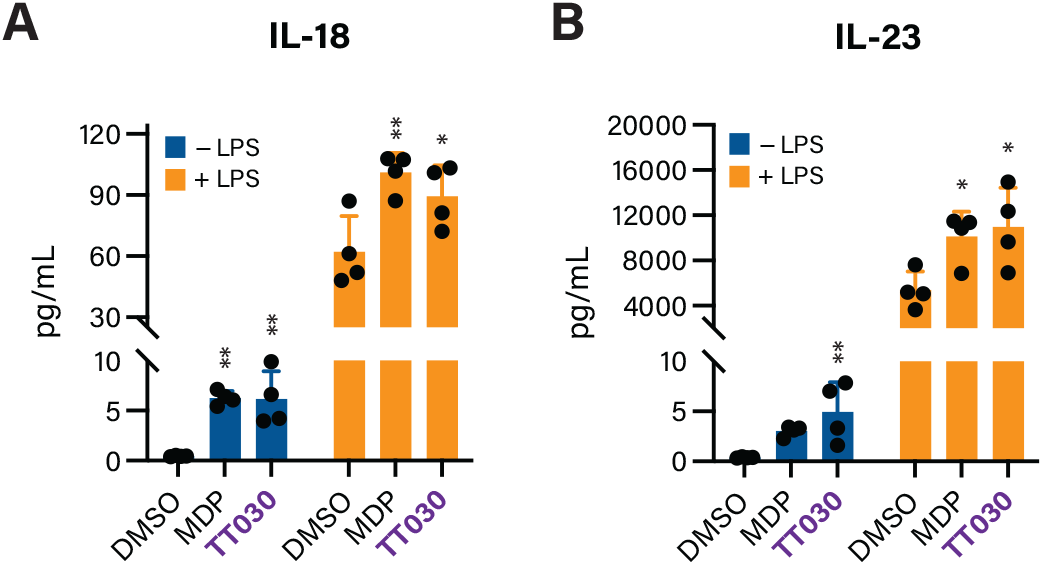
NOD2 agonist TT030 activates cytokine production in vitro. **a-b**, Measurements of secreted cytokine levels from human peripheral mononuclear blood cells (PMBCs) treated with the indicated compounds for 24 h without or with lipopolysaccharide (LPS). Cytokine secretion assays and NOD activation assays were analyzed using a one-way ANOVA with Dunnett’s multiple comparisons test compared to the DMSO control. * *P* < 0.05, ** *P* < 0.01.

**Extended Fig. 4.**
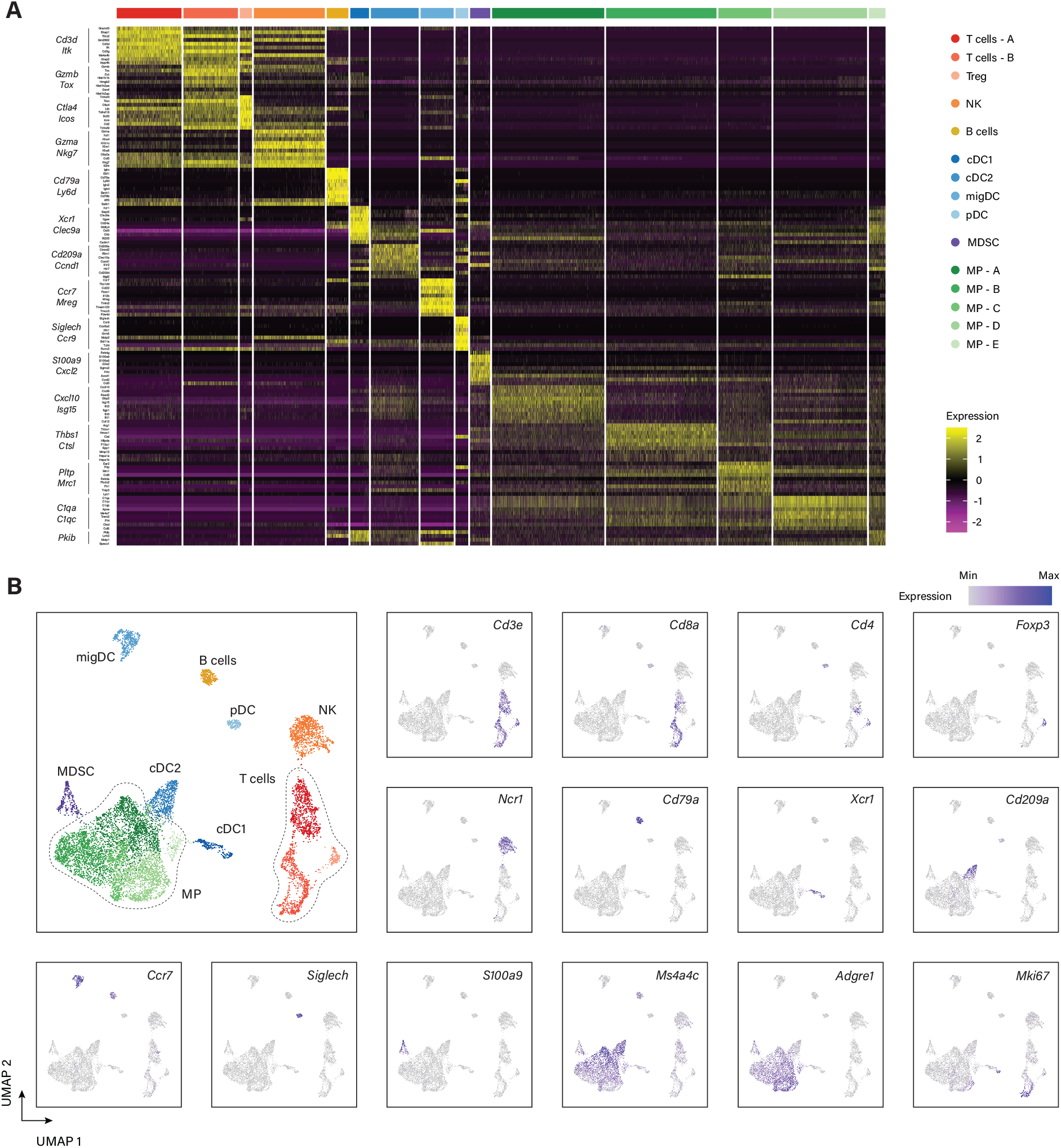
Single cell RNA sequencing clustering and classification. **a**, Heat map of top 10 globally differentially expressed genes in each cluster (within cluster vs. other cells). Clusters are indicated by color, and representative genes are listed on the left. **b**, Heat map of individual genes used to classify gene clusters mapped onto a UMAP reduction of individual cells.

**Extended Fig. 5.**
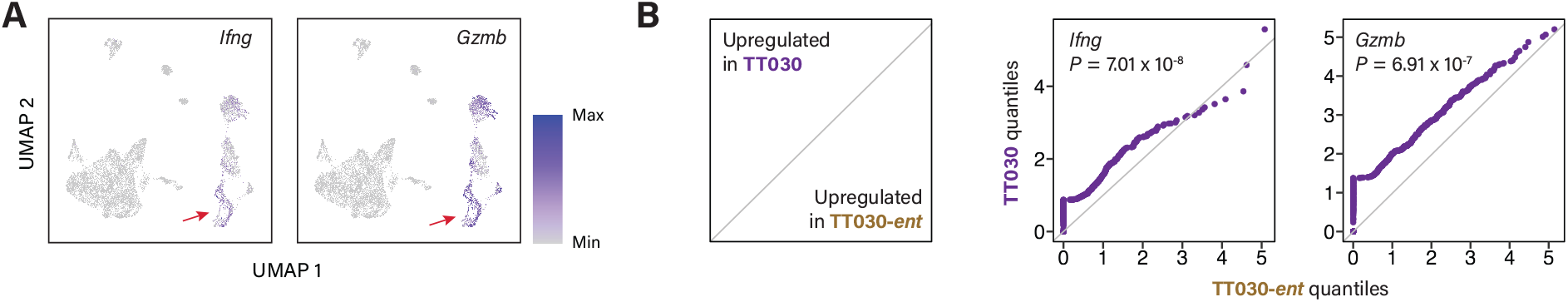
TT030 upregulates cytotoxic T cell gene expression. **a**, Heat map of interferon gamma (*Ifng*) and granzyme B (*Gzmb*) expression mapped onto a UMAP reduction of individual cells. Red arrow indicates the increased cluster of CD8^+^ T cells. **b**, Quantile-quantile (QQ) plots of cytotoxic T cell genes *Ifng* and *Gzmb*. QQ plots were analyzed using a two-sided Wilcoxon rank-sum test; exact *P* values are given.

