## Supplementary Information for "*N*-arylpyrazole NOD2 agonists promote immune checkpoint inhibitor therapy"

**Affiliations**
[1] Department of Immunology and Microbiology, Scripps Research, La Jolla, CA 92037.

[2] Department of Chemistry, University of California, Irvine, Irvine, CA 92697.

[3] Laboratory of Chemical Biology and Microbial Pathogenesis, The Rockefeller University, New York, NY 10065.

[4] Department of Chemistry, Scripps Research, La Jolla, CA 92037.

[#]These authors contributed equally.

**Contents**

Description of **Tables S1-S15** .............................. S2

Synthesis and chemical characterization ............... S3

**Supplementary Tables S1-S15**

Differential gene expression and gene set enrichment analyses for each individual cell cluster within the scRNA-seq dataset comparing CD45$^+$ cells from B16 tumors treated with anti-PD-L1 and either **TT030** or **TT030-*ent***. Tables are labelled with the appropriate cell cluster and numbered in order as described in **Figure 4**. Each table contains separate sheets for differential gene expression and analyses from hallmark (H), curated (C2:CP), and biological process gene ontology gene sets (C5:GOBP). Positive values correlate with upregulation in the **TT030** sample.

**Synthesis and chemical characterization.**

Diethyl *D*-glutamate hydrochloride

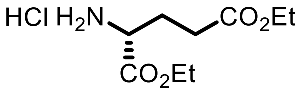


To EtOH (20 mL) was added acetyl chloride (2.1 mL, 0.03 mmol) on ice. After 30 min, *D*-glutamic acid (2.00 g, 0.014 mmol) was added in one portion, and the reaction mixture was refluxed for 4 h. The solvent was evaporated to provide crude material as white solid.

$^1$H NMR (DMSO-$d_6$, 600 MHz) δ 8.68 (br, 3H), 4.19 (q, 2 H, *J* = 7.0 Hz), 4.07 (q, 2H *J* = 7.0 Hz), 4.00 (m, 1H), 2.54 (m, 1H), 2.47 (m, 1H), 2.06 (m, 2H), 1.23 (t, 3H, *J* = 7.1 Hz), 1.18 (t, 3H, *J* = 7.1 Hz).

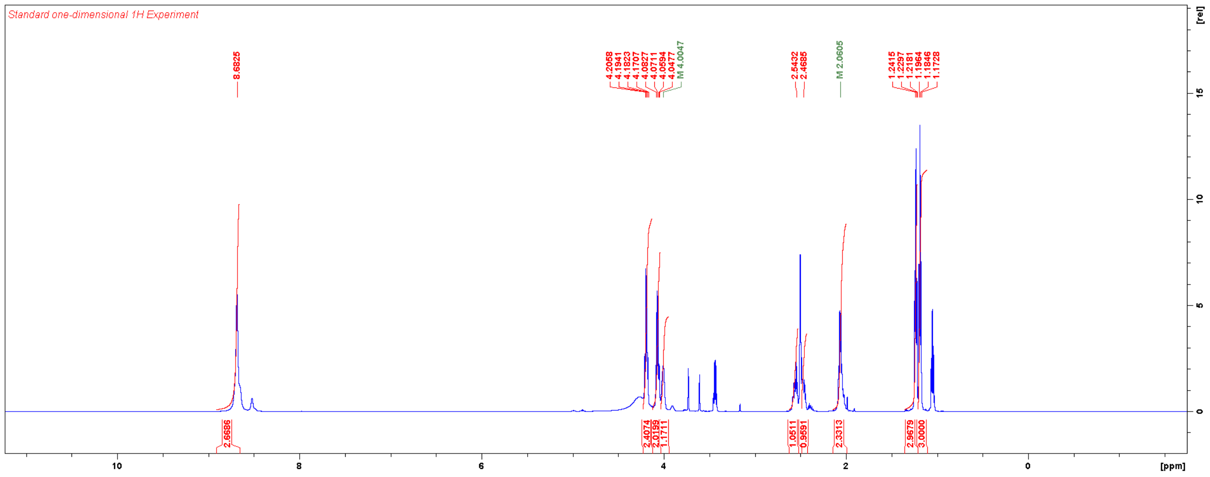

Diethyl (tert-butoxycarbonyl)-*L*-valyl-*D*-glutamate

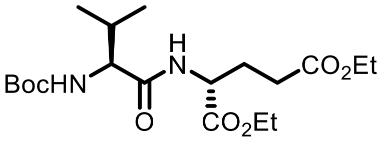

To a solution of Boc-*L*-valine-OH (906 mg, 4.17 mmol), diethyl *D*-glutamate hydrochloride (1000 mg, 4.17 mmol), EDC-HCl (500 mg, 4.17 mmol) and HOBt-x$H_2O$ (620 mg, 4.59 mmol) in DMF (20 mL) was added TEA (1.2 mL, 8.34 mmol) at 0 °C. The reaction mixture was allowed to warm to RT overnight. The reaction mixture was diluted with EtOAc, washed with aqueous $NaHCO_3$, aqueous $NH_4Cl$, and brine, dried over $MgSO_4$, filtered, and concentrated. The residue was purified by column chromatography (10% EtOAc/DCM). Yield: 914 mg (2.27 mmol, 54%), white powder.

$^1$H NMR ($CDCl_3$, 600 MHz) δ 6.67 (br, 1H), 4.97 (m, 1H), 4.59 (m, 1H), 4.20 (q, 2H, *J* = 7.0 Hz), 4.14 (q, 2H, *J* = 7.0 Hz), 4.00 (m 1H), 2.38 (m, 2H), 2.21 (m, 2H), 2.01 (m, 1H), 1.45 (s, 9H), 1.27 (m, 6H), 0.98 (d, 3H, *J* = 6.3 Hz), 0.91 (d, 3H, *J* = 6.5 Hz).

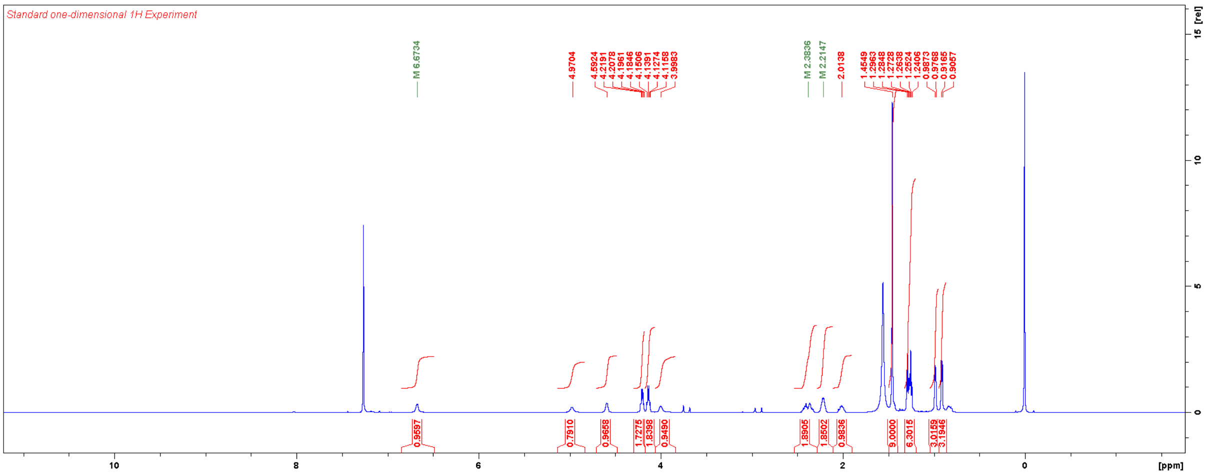

Diethyl (tert-butoxycarbonyl)glycyl-*L*-valyl-*D*-glutamate

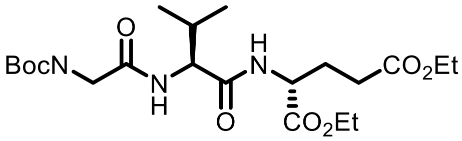

Diethyl (tert-butoxycarbonyl)-*L*-valyl-*D*-glutamate (150 mg, 0.37 mmol) was deprotected with 25% TFA/DCM (4 mL). To a solution of Boc-Gly-OH (72 mg, 0.41 mmol), HBTU (79 mg, 0.41 mmol), and TEA (260 µL, 1.86 mmol) in DMF (4 mL) was added the deprotected amine at 0 °C. The reaction mixture was allowed to warm to RT over 1 h, diluted with EtOAc, washed with aqueous $NaHCO_3$, aqueous $NH_4Cl$, and brine, dried over $MgSO_4$, filtered, and concentrated. The residue was purified with column chromatography (25% EtOAc/DCM). Yield: 59 mg (0.13 mmol, 34%), yellow oil.

$^1$H NMR ($CDCl_3$, 600 MHz) δ 7.00 (br, 1H), 6.63 (br, 1H), 5.23 (br, 1H), 4.53 (m, 1H), 4.38 (m, 1H), 4.21 (q, 2H, *J* = 7.0 Hz), 4.16 (q, 2H, *J* = 7.0 Hz), 3.91 (m, 1H), 3.82 (m, 1H), 2.5–2.0 (m, 5H), 1.49 (s, 9H), 1.29 (m, 6H), 0.99 (d, 3H, *J* = 6.7 Hz), 0.95 (d, 3H, *J* = 6.8 Hz).

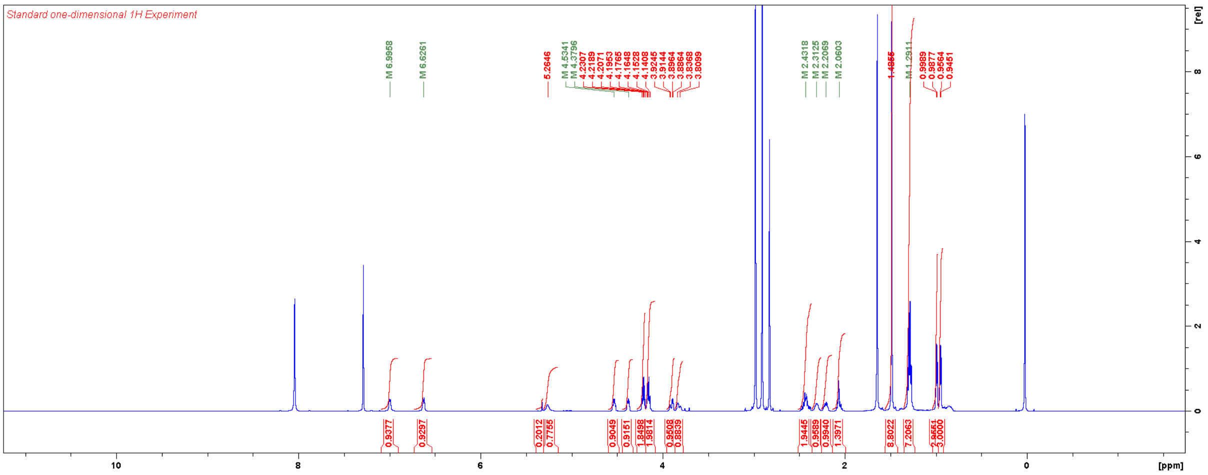

Diethyl cinnamoylglycyl-*L*-valyl-*D*-glutamate (**CinGVE**)

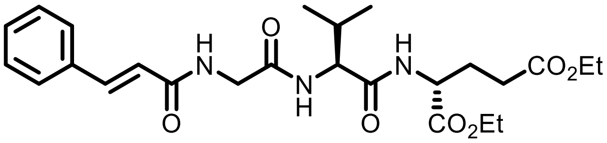

Diethyl (tert-butoxycarbonyl)glycyl-*L*-valyl-*D*-glutamate (30 mg, 0.065 mmol) was deprotected with 25% TFA/DCM (1.5 mL). To the solution of the deprotected amine and TEA (45 µL, 0.33 mmol) in DMF (0.6 mL) was added cinnamoyl chloride (22 mg, 0.13 mmol) at 0 °C. The reaction mixture was allowed to warm to RT over 30 min, diluted with EtOAc, washed with aqueous $NaHCO_3$, aqueous $NH_4Cl$, and brine, dried over $MgSO_4$, filtered, and concentrated. The residue was purified with flash chromatography using a gradient of EtOAC in DCM. Yield: 16 mg (33 µmol, 50%), white solid.

$^1$H NMR ($CDCl_3$, 600 MHz) δ 7.69 (d, 1H, $J$ = 15.6 Hz), 7.54 (m, 2H), 7.40 (m, 3H), 6.98 (br, 1H), 6.66 (br, 1H), 6.52 (br, 1H), 6.51 (d, 1H, $J$ = 15.7 Hz), 4.57 (m, 1H), 4.41 (m, 1H), 4.17 (m, 1H), 2.5–2.0 (m, 5H), 1.27 (m, 6H), 1.00 (d, 3H, $J$ = 6.7 Hz), 9.65 (d, 3H, $J$ = 6.8 Hz).

HRMS m/z: $[M+H]^+$ Calcd for $C_{25}H_{36}N_3O_7$ 490.2548; Found 490.2559.

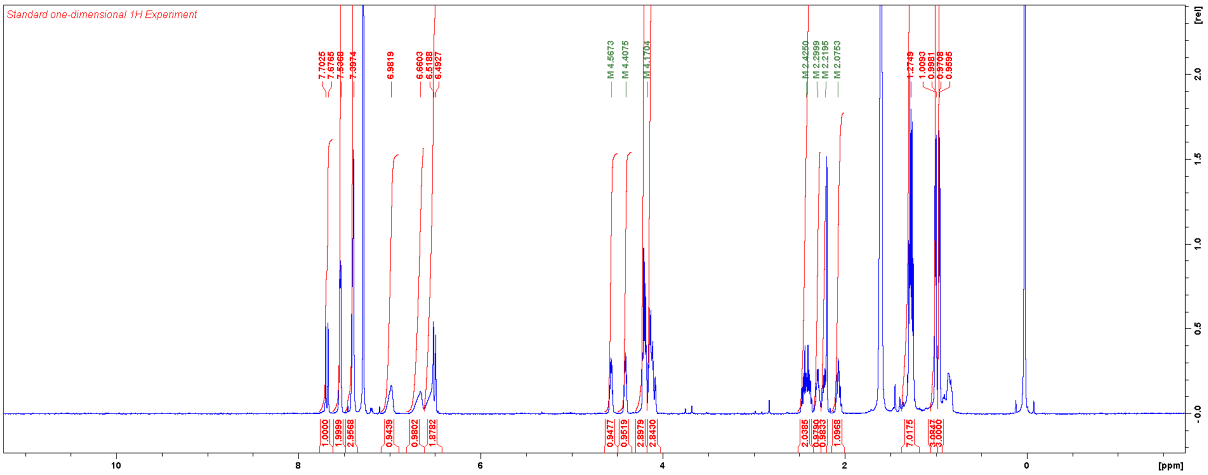

Diethyl (4-phenylbutanoyl)-*L*-valyl-*D*-glutamate (**TT001**)

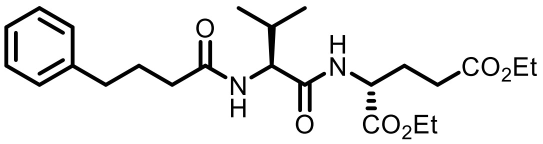


**TT001** was synthesised according to the general procedure for HATU-mediated coupling. Yield: 25 mg (0.056 mmol, 56%).

$^{1}$H NMR ($CDCl_3$, 600 MHz) δ 7.28 (m, 2H), 7.18 (m, 3H), 6.70 (br, 1H), 5.96 (br, 1H), 4.54 (m, 1H), 4.35 (m, 1H), 4.18 (q, 2H, *J* = 7.0 Hz), 4.12 (q, 2H, *J* = 7.1 Hz), 2.65 (t, 2H, *J* = 7.5 Hz), 2.50–1.90 (m, 9H), 1.26 (m, 6H), 0.96 (d, 3H, *J* = 6.8 Hz), 0.93 (d, 3H, *J* = 6.8 Hz).

HRMS m/z: $[M+H]^+$ Calcd for $C_{24}H_{37}N_2O_6$ 449.2646; Found 449.2656.

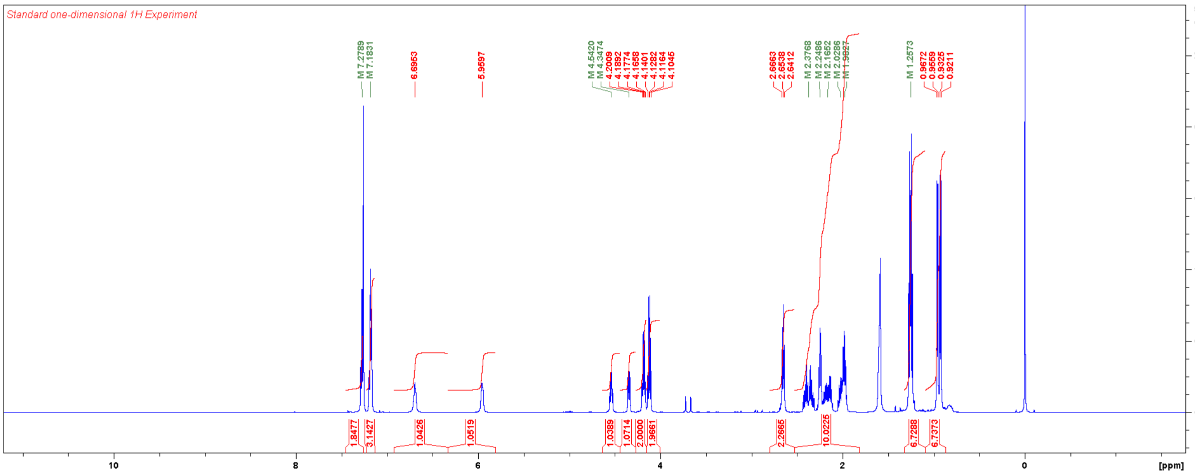

Diethyl ((*E*)-3-(4-fluorophenyl)acryloyl)-*L*-valyl-*D*-glutamate (**TT002**)

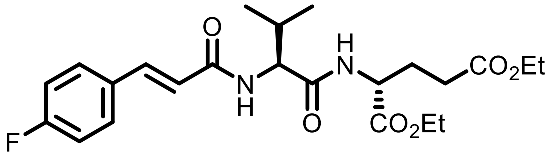


To the solution of the dipeptide (0.10 mmol) and DIPEA (87 µL, 0.50 mmol) in DMF (1 mL) was added (4-fluorophenyl)acryloyl chloride (37 mg, 0.20 mmol) at 0 °C. The reaction mixture was allowed to warm to RT over 4 h, quenched with $NH_4Cl$, extracted three times with DCM, dried over $MgSO_4$, filtered, and concentrated. The residue was purified with flash chromatography using a gradient of EtOAC in DCM. Yield: 26 mg (0.058 mmol, 58%).

$^1$H NMR ($CDCl_3$, 600 MHz) δ 7.60 (d, 1H, $J$ = 15.5 Hz), 7.49 (dd, 2H, $J_{H,H}$ = 8.3 Hz, $^4J_{H,F}$ = 5.5 Hz), 7.06 (dd, 2H, $J_{H,H}$ = 8.5 Hz, $^3J_{H,F}$ = 8.5 Hz), 6.81 (d, 1H, $J$ = 7.1 Hz), 6.38 (d, 1H, $J$ = 15.4 Hz), 6.19 (d, 1H, $J$ = 8.5 Hz), 4.56 (m, 1H), 4.49 (m, 1H), 4.20 (m, 2H), 4.13 (q, 2H, $J$ = 7.1 Hz), 2.40 (m, 2H), 2.21 (m, 2H), 2.04 (m, 1H), 1.26 (m, 6H), 1.01 (d, 3H, $J$ = 6.7 Hz), 0.98 (d, 3H, $J$ = 6.8 Hz).

HRMS m/z: $[M+H]^+$ Calcd for $C_{23}FH_{32}N_2O_6$ 451.2239; Found 451.2251.

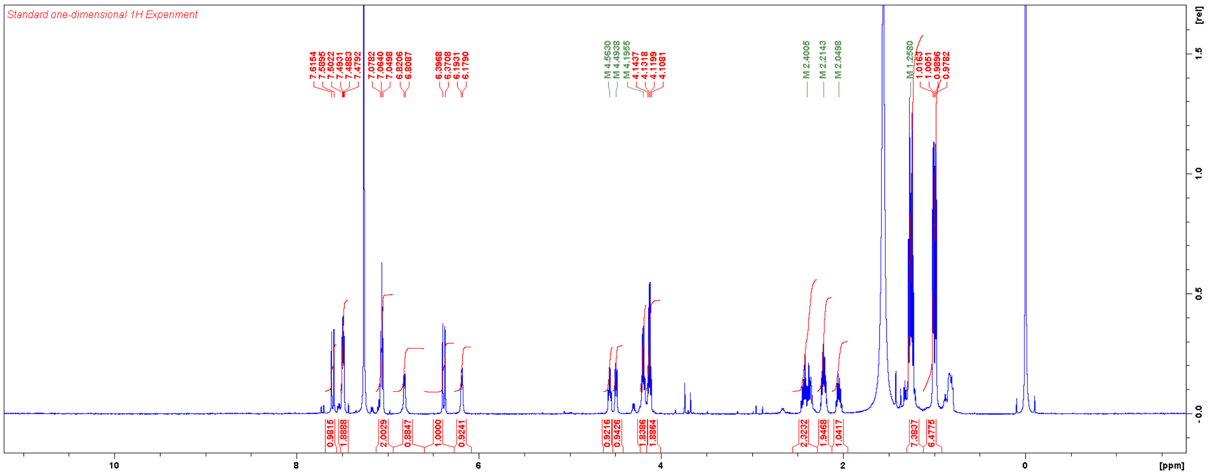

Diethyl (1-(4-fluorobenzyl)cyclopentane-1-carbonyl)-*L*-valyl-*D*-glutamate (**TT003**)

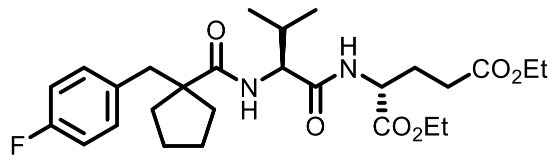

**TT003** was synthesised according to the general procedure for HATU-mediated coupling. Yield: 36 mg (0.071 mmol, 71%).

HRMS m/z: $[M+H]^+$ Calcd for $C_{27}FH_{40}N_2O_6$ 507.2865; Found 507.2877.

Diethyl (*rac*-(1*R*,2*R*)-2-(3,4-dichlorophenyl)cyclopropane-1-carbonyl)-*L*-valyl-*D*-glutamate (**TT004**)

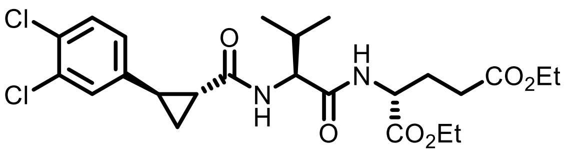

**TT004** (racemic mixture) was synthesised according to the general procedure for HATU-mediated coupling. Yield: 11 mg (0.021 mmol, 21%).

$^1$H NMR (CDCl$_3$, 600 MHz) δ 7.32 (m, 1H), 7.18 (m, 1H), 6.93 (m, 1H), 6.69 (m, 1H), 6.24 (m, 1H), 4.56 (m, 1H), 4.37 (m, 1H), 4.20 (m, 2H), 4.12 (m, 2H), 2.57–1.94 (m, 6H), 1.67 (m, 2H), 1.26 (m, 7H), 0.96 (m, 2H).

$^{13}$C NMR (CDCl$_3$, 150 MHz) δ 172.8, 171.5, 171.4, 171.03, 170.98, 141.1, 141.0, 132.5, 132.4, 130.4, 130.3, 130.2, 128.4, 128.1, 125.7, 125.6, 61.8, 60.8, 58.5, 58.4, 51.9, 31.2, 30.9, 30.3, 27.1, 27.0, 26.54, 26.48, 24.3, 24.2, 19.3, 17.9, 16.4, 16.1, 14.1.

HRMS m/z: [M+H]$^+$ Calcd for $C_{24}Cl_2H_{33}N_2O_6$ 515.1710; Found 515.1723.

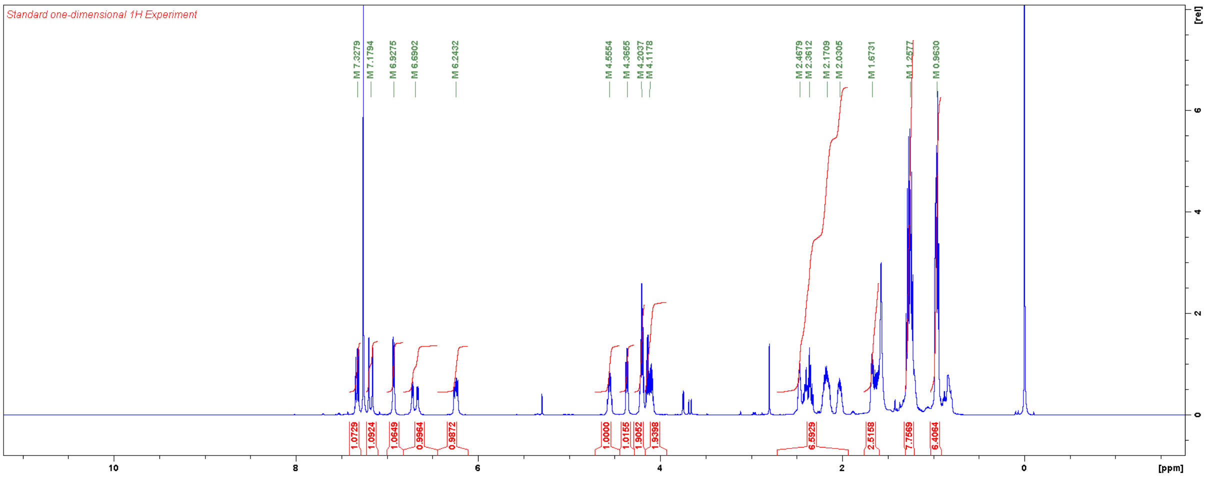


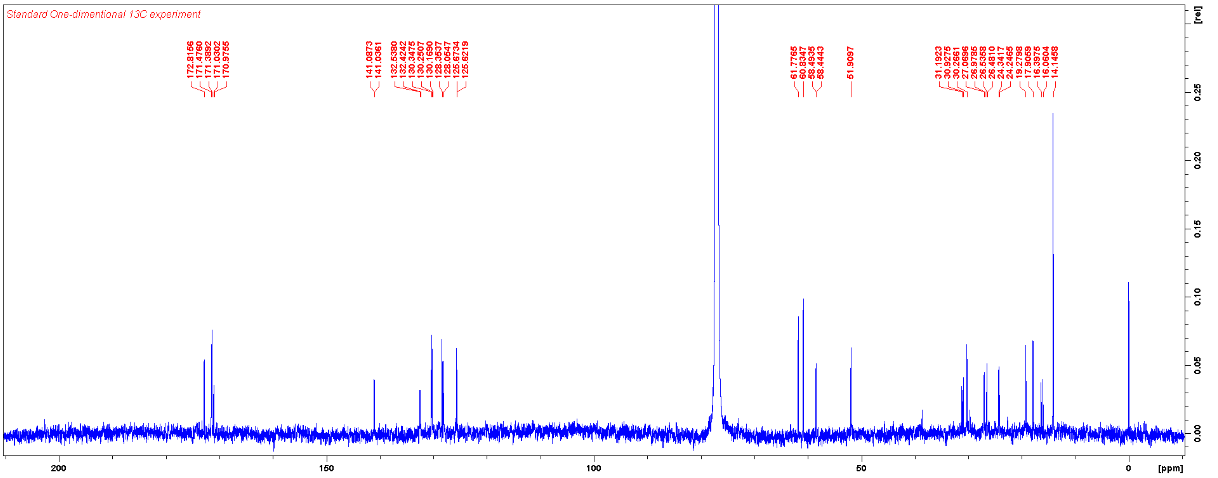

Diethyl (3-(3,4-difluorophenyl)-2-methylpropanoyl)-*L*-valyl-*D*-glutamate (**TT005**)

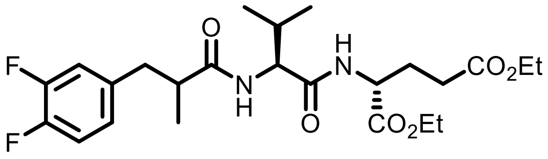


**TT005** (racemic mixture) was synthesised according to the general procedure for HATU-mediated coupling. Yield: 41 mg (0.085 mmol, 85%).

$^1$H NMR ($CDCl_3$, 600 MHz) δ 7.12–6.75 (m, 3H), 6.58 (br, 1H), 5.95 (br, 1H), 4.52 (m, 1H), 4.23 (m, 2H), 4.13 (m, 2H), 3.11–1.88 (m, 8H), 1.39–1.11 (m, 9H), 0.98–0.66 (m, 6H).

$^{13}$C NMR ($CDCl_3$, 150 MHz) δ 175.4 & 175.1 (s), 172.6 (s), 171.3 (s), 171.2 (s), 150.1 (dd, $^1J_{C,F}$ = 246 Hz, $^2J_{C,F}$ = ~ 12 Hz), 149.0 (dd, $^1J_{C,F}$ = 245 Hz, $^2J_{C,F}$ = ~ 11 Hz), 136.7 & 136.6 (s), 124.9 (s), 117.7 (dd, $^2J_{C,F}$ = 15 Hz), 117.0 (d, $^2J_{C,F}$ = 17 Hz), 61.73 (s), 61.67 (s), 60.8 (s), 58.0 (s), 43.7 & 43.2 (s), 39.5 & 39.0 (s), 31.2 & 30.9 (s), 30.2 (s), 27.2 & 27.0 (s), [three of 19.2 (s), 18.9 (s), 17.9 (s), 17.82 (s), 17.79 (s), 17.7 (s)], 14.13 (s), 14.09 (s).

HRMS m/z: $[M+H]^+$ Calcd for $C_{24}F_2H_{35}N_2O_6$ 485.2458; Found 485.2469.

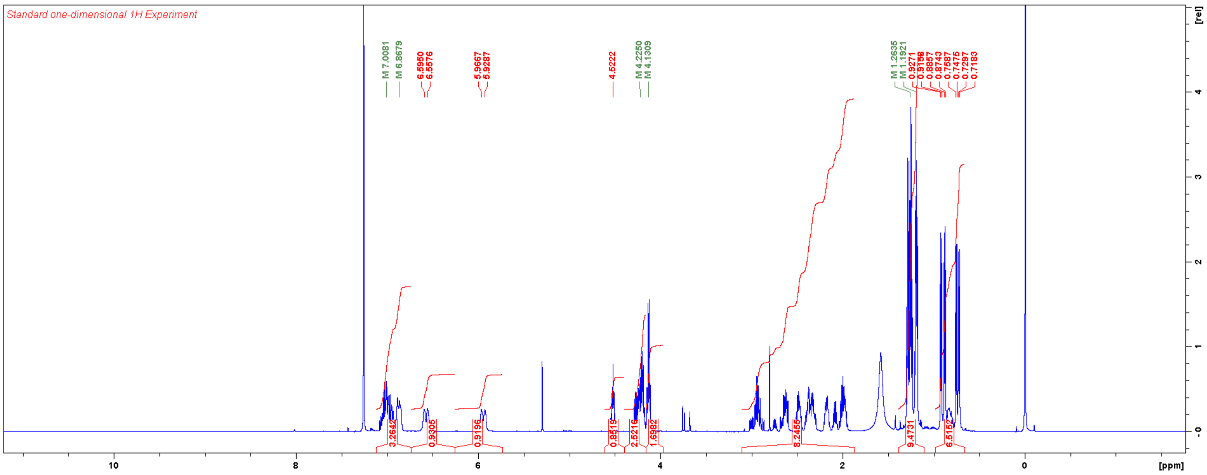


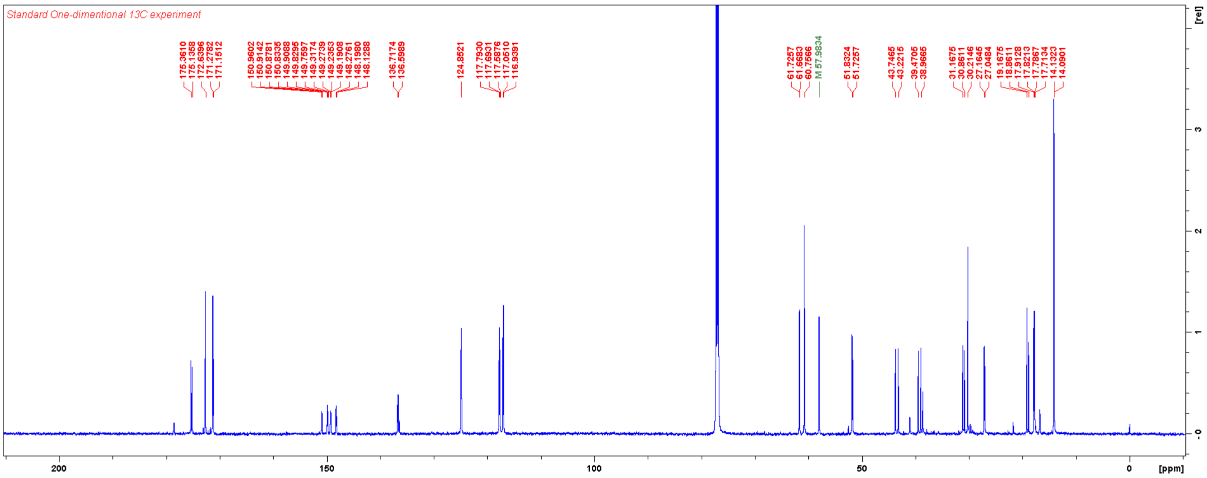

Diethyl (2-benzylbicyclo[2.2.1]heptane-2-carbonyl)-*L*-valyl-*D*-glutamate (**TT006**)

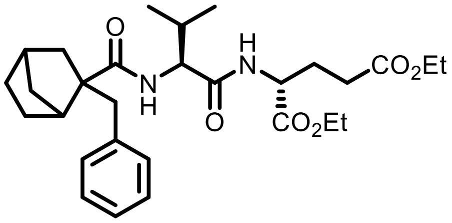


**TT006** (racemic mixture) was synthesised according to the general procedure for HATU-mediated coupling. Yield: 37 mg (0.072 mmol, 72%).

$^1$H NMR ($CDCl_3$, 600 MHz) δ 7.23–7.00 (m, 5H), 6.52 (br, 1H), 5.90 (br, 1H), 4.54 (m, 1H), 4.21 (m, 2H), 4.12 (q, 2H, $J$ = 7.1 Hz), 3.14 (m, 1H), 2.73 (m, 1H), 2.50–0.65 (m, 28H).

$^{13}$C NMR ($CDCl_3$, 150 MHz) δ 176.1, 175.8, 172.6, 171.3, 170.84, 170.80, 137.7, 129.9, 129.6, 128.15, 128.11, 126.4, 126.3, 61.73, 61.66, 60.7, 58.7, 58.4, 56.4, 56.2, 51.70, 51.67, 46.5, 46.4, 45.9, 45.6, 38.6, 38.23, 38.18, 37.8, 37.64, 37.60, 37.5, 36.5, 31.4, 31.1, 30.5, 30.22, 30.18, 28.3, 28.2, 27.5, 27.4, 25.8, 25.7, 19.2, 18.3, 18.2, 14.2.

HRMS m/z: $[M+H]^+$ Calcd for $C_{29}H_{43}N_2O_6$ 515.3116; Found 515.3130.

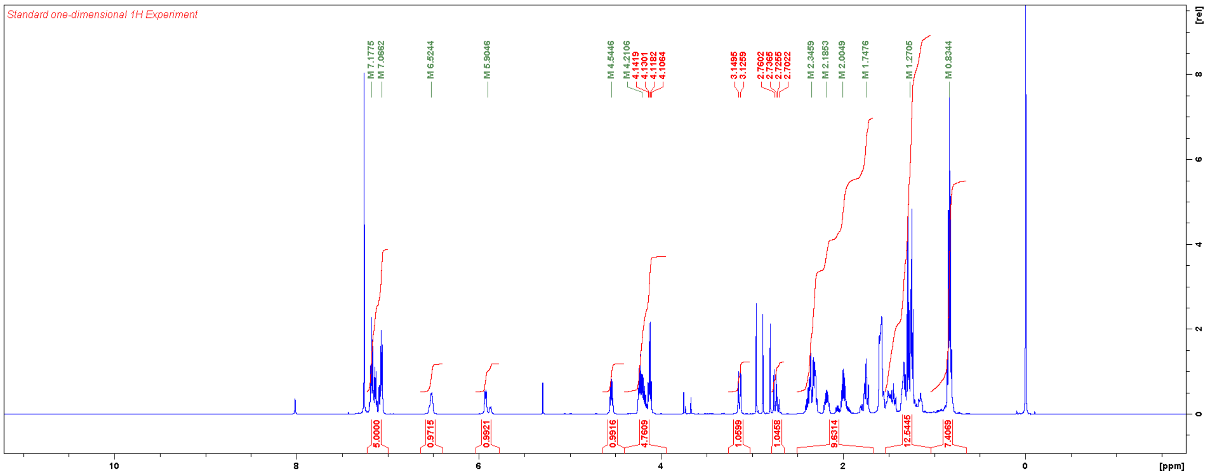


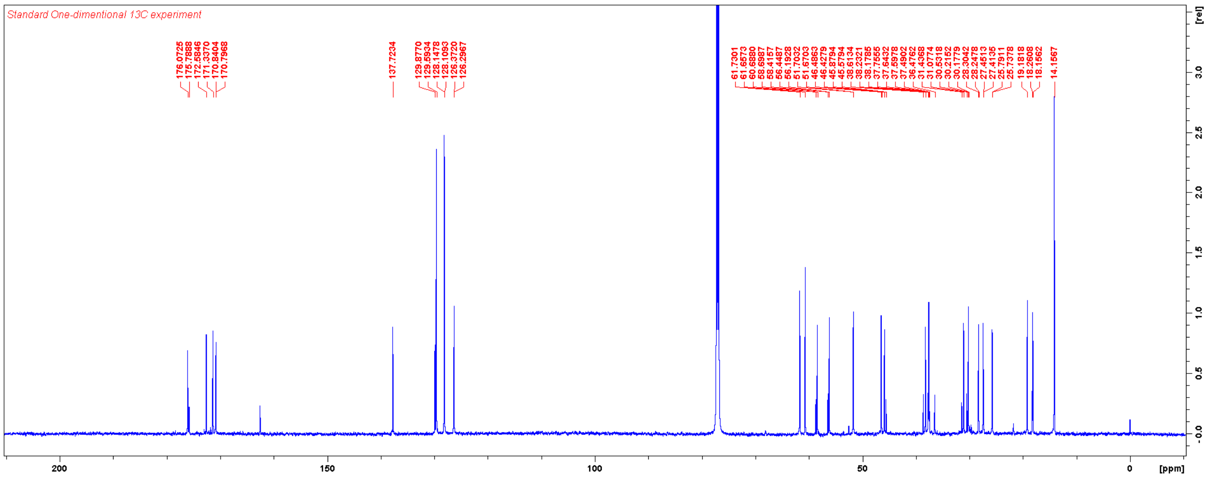

Diethyl (1-(4-fluorophenyl)-3,5-dimethyl-1*H*-pyrazole-4-carbonyl)-*L*-valyl-*D*-glutamate (**TT007**)

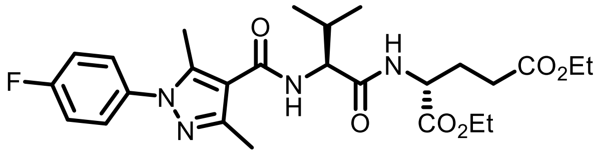


**TT007** was synthesised according to the general procedure for HATU-mediated coupling. Yield: 35 mg (0.067 mmol, 75%).

$^1$H NMR ($CDCl_3$, 600 MHz) δ 7.36 (dd, 2H, $J_{H,H}$ = 8.9 Hz, $^4J_{H,F}$ = 4.8 Hz), 7.17 (dd, 2H, $J_{H,H}$ = 8.5 Hz, $^3J_{H,F}$ = 8.5 Hz), 6.72 (d, 1H, $J$ = 7.3 Hz), 6.29 (d, 1H, $J$ = 8.2 Hz), 4.57 (m, 2H), 4.20 (m, 2H), 4.13 (m, 2H), 2.53 (s, 3H), 2.48 (s, 3H), 2.46–1.94 (m, 5H), 1.26 (m, 6H), 1.05 (d, 3H, $J$ = 6.8 Hz), 1.02 (d, 3H, $J$ = 6.8 Hz).

$^{13}$C NMR ($CDCl_3$, 150 MHz) δ 172.8 (s), 171.4 (s), 171. 2 (s), 164.5 (s), 162.2 (d, $^1J_{C,F}$ = 247 Hz), 147.5 (s), 142.5 (s), 135.0 (s), 127.5 (d, $^3J_{C,F}$ = 9 Hz), 116.2 (d, $^2J_{C,F}$ = 22 Hz), 114.4 (s), 61.7 (s), 60.8 (s), 57.9 (s), 52.0 (s), 31.3 (s), 30.3 (s), 27.1 (s), 19.5 (s), 17.9 (s), 14.21 (s), 14.15 (s), 14.12 (s), 12.2 (s).

HRMS m/z: $[M+H]^+$ Calcd for $C_{26}FH_{36}N_4O_6$ 519.2613; Found 519.2624.

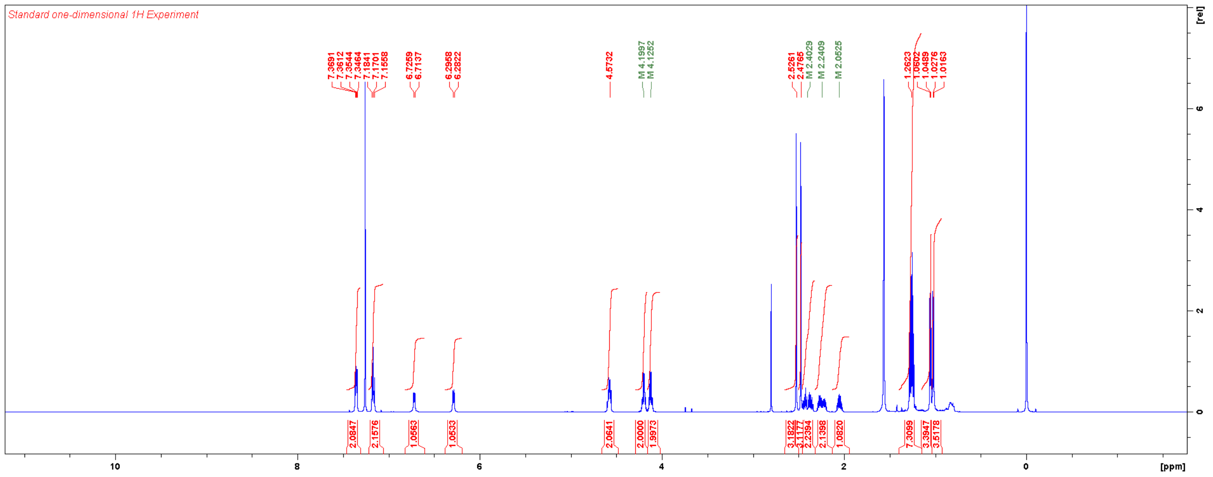


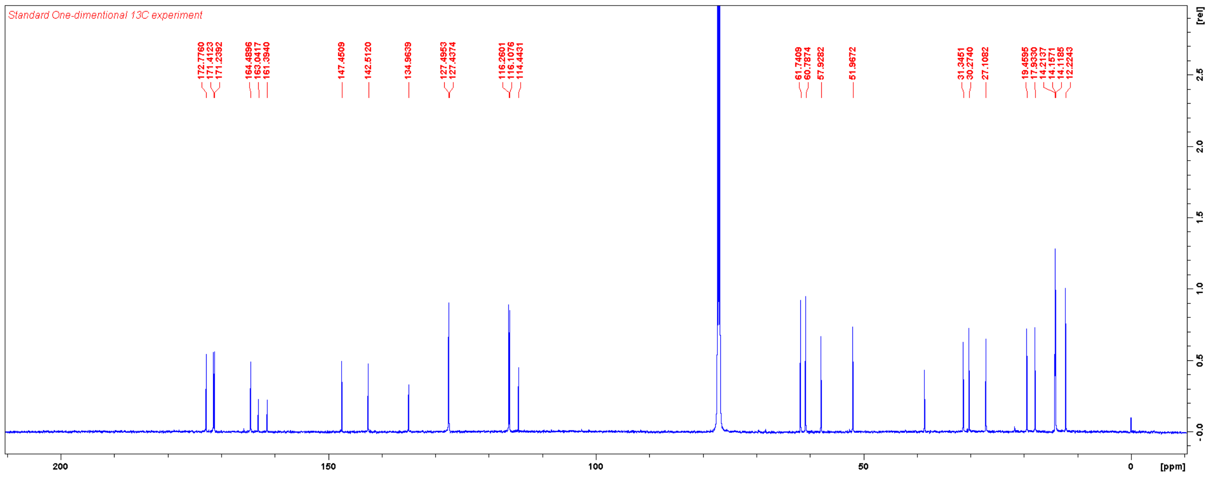

Diethyl (1-phenyl-3-(m-tolyl)-1*H*-pyrazole-4-carbonyl)-*L*-valyl-*D*-glutamate (**TT008**)

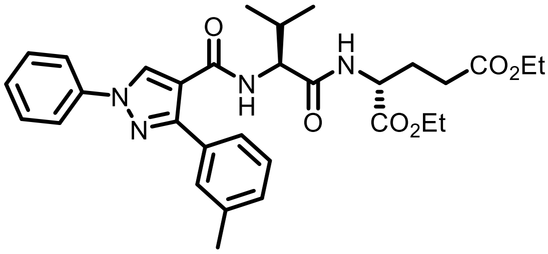


**TT008** was synthesised according to the general procedure for HATU-mediated coupling. Yield: 34 mg (0.060 mmol, 67%).

$^{1}$H NMR ($CDCl_3$, 600 MHz) δ 8.54 (s, 1H), 7.75 (d, 2H, *J* = 7.9 Hz), 7.49–7.29 (m, 7H), 6.76 (d, 1H, *J* = 7.4 Hz), 6.13 (d, 1H, *J* = 8.1 Hz), 4.53 (m, 1H), 4.39 (m, 1H), 4.18 (q, 2H, *J* = 7.1 Hz), 4.08 (m, 2H), 2.42 (s, 3H), 2.40–2.00 (m, 5H), 1.26 (t, 3H, *J* = 7.1 Hz), 1.21 (t, 3H, *J* = 7.1 Hz), 0.84 (d, 3H, *J* = 6.8 Hz), 0.61 (d, 3H, *J* = 6.8 Hz).

$^{13}$C NMR ($CDCl_3$, 150 MHz) δ 172.9, 171.5, 170.9, 162.9, 151.4, 139.3, 139.0, 132.1, 131.3, 130.3, 130.0, 129.6, 129.1, 127.3, 126.5, 119.4, 117.8, 61.6, 60.7, 58.4, 51.8, 30.3, 30.0, 27.1, 21.4, 19.4, 17.1, 14.1.

HRMS m/z: $[M+H]^+$ Calcd for $C_{31}H_{39}N_4O_6$ 563.2864; Found 563.2878.

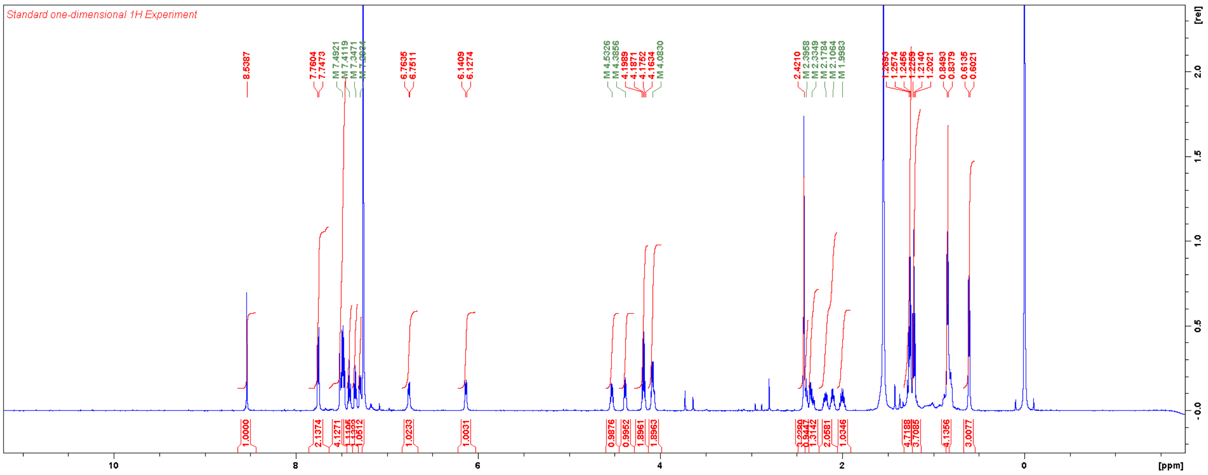


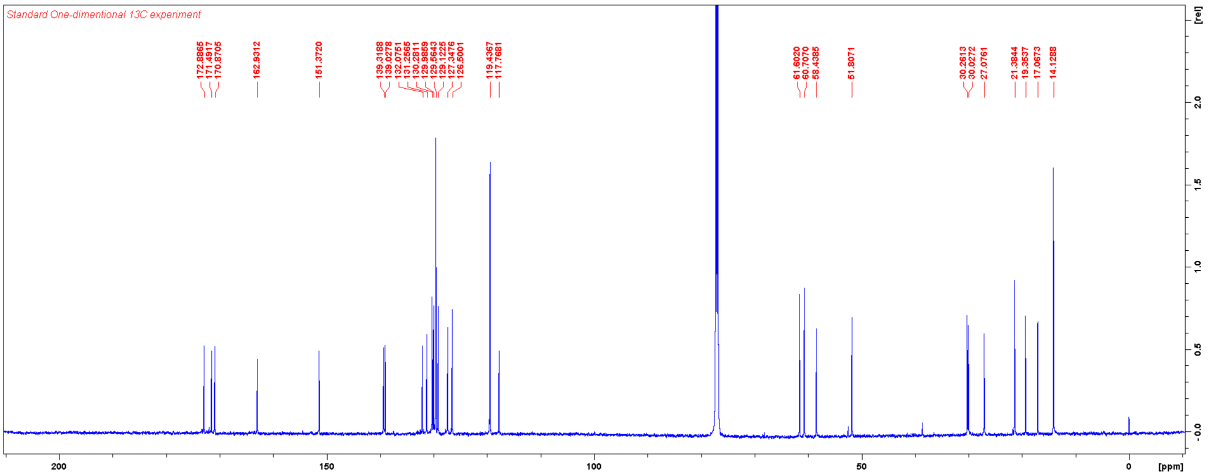

Diethyl (1-(4-fluorophenyl)-3-(m-tolyl)-1*H*-pyrazole-4-carbonyl)-*L*-valyl-*D*-glutamate (**TT009**)

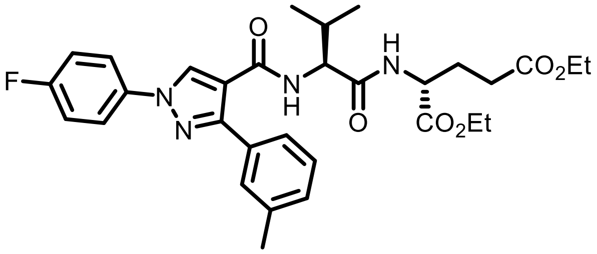


**TT009** was synthesised according to the general procedure for HATU-mediated coupling. Yield: 44 mg (0.076 mmol, 84%).

$^1$H NMR ($CDCl_3$, 600 MHz) δ 8.47 (s, 1H), 7.72 (m, 2H), 7.49 (m, 2H), 7.41 (t, 1H, $J$ = 7.6 Hz), 7.30 (d, 1H, $J$ = 7.6 Hz), 7.17 (m, 2H), 6.74 (d, 1H, $J$ = 7.4 Hz), 6.13 (d, 1H, $J$ = 8.0 Hz), 4.53 (m, 1H), 4.38 (m, 1H), 4.18 (q, 2H, $J$ = 7.1 Hz), 4.09 (m, 2H), 2.42 (s, 3H), 2.39–2.00 (m, 5H), 1.26 (t, 3H, $J$ = 7.1 Hz), 1.22 (t, 3H, $J$ = 7.1 Hz), 0.84 (d, 3H, $J$ = 6.8 Hz), 0.61 (d, 3H, $J$ = 6.8 Hz).

$^{13}$C NMR ($CDCl_3$, 150 MHz) δ 172.9 (s), 171.5 (s), 170.9 (s), 162.8 (s), 161.2 (d, $^1J_{C,F}$ = 246 Hz), 151.4 (s), 139.0 (s), 135.7 (s), 131.9 (s), 131.3 (s), 130.3 (s), 129.9 (s), 129.1 (s), 126.4 (s), 121.3 (d, $^3J_{C,F}$ = 8 Hz), 117.9 (s), 116.4 (d, $^2J_{C,F}$ = 23 Hz), 61.6 (s), 60.7 (s), 58.4 (s), 51.8 (s), 30.3 (s), 30.1 (s), 27.1 (s), 21.4 (s), 19.3 (s), 17.1 (s), 14.1 (s), 14.1 (s).

HRMS m/z: $[M+H]^+$ Calcd for $C_{31}FH_{38}N_4O_6$ 581.2770; Found 581.2783.

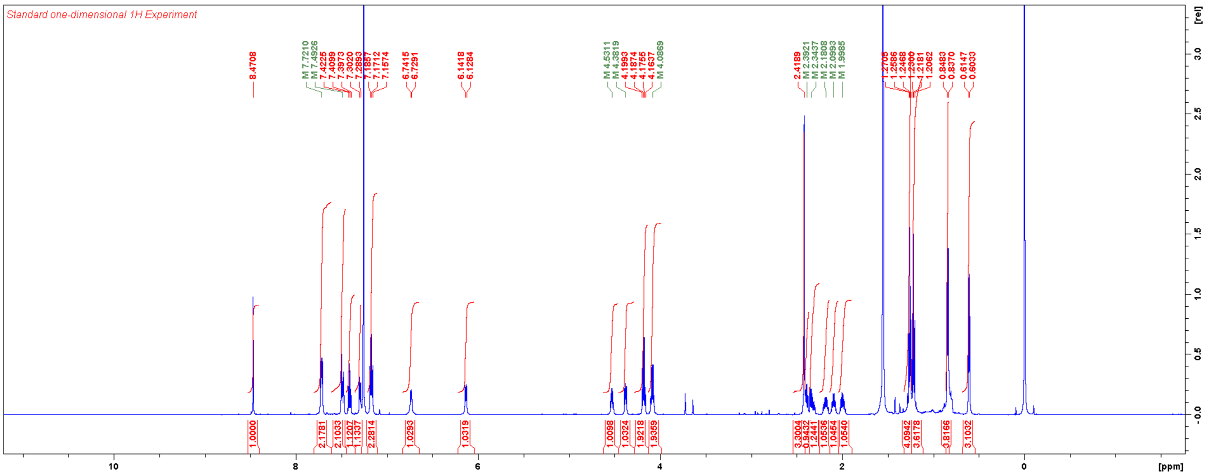


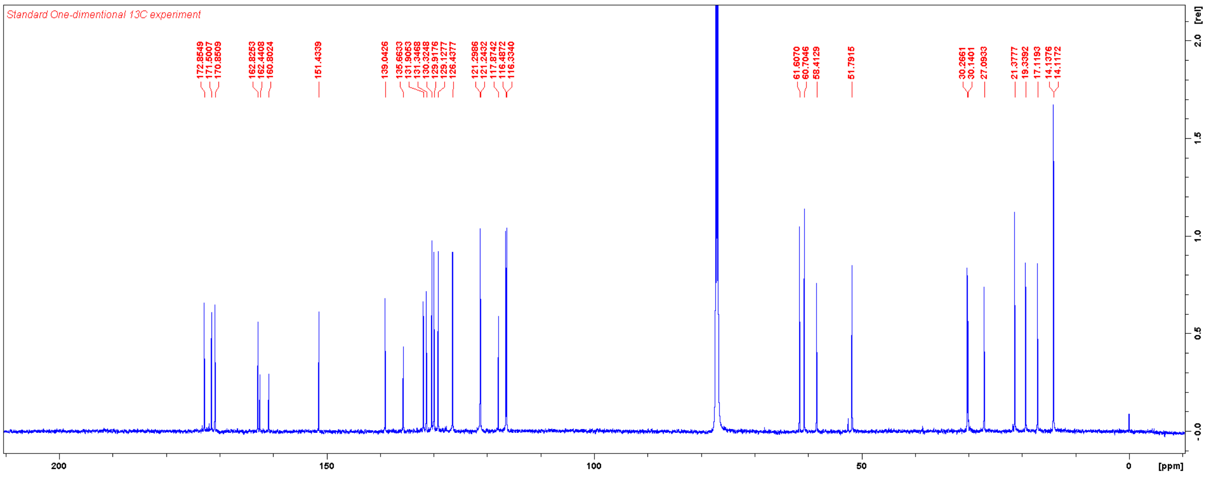

Diethyl (1-(4-fluorophenyl)-5-methyl-1*H*-pyrazole-4-carbonyl)-*L*-valyl-*D*-glutamate (**TT010**)

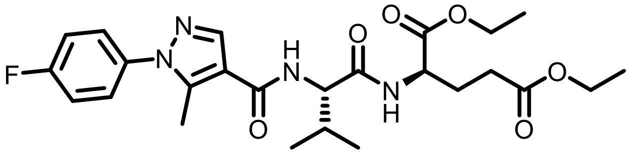


**TT010** was synthesised according to the general procedure for HATU-mediated coupling. Yield: 30 mg (0.059 mmol, 99%).

$^1$H NMR ($CDCl_3$, 600 MHz) δ 7.87 (s, 1H), 7.39 (dd, 2H, $J_{H,H}$ = 8.8 Hz, $^4J_{H,F}$ = 4.8 Hz), 7.19 (dd, 2H, $J_{H,H}$ = 8.5 Hz, $^3J_{H,F}$ = 8.5 Hz), 6.77 (d, 1H, $J$ = 7.0 Hz), 6.35 (d, 1H, $J$ = 8.3 Hz), 4.58 (m, 1H), 4.52 (m, 1H), 4.20 (m, 2H), 4.13 (q, 2H, $J$ = 7.1 Hz), 2.55 (s, 3H), 2.41 (m, 2H), 2.24 (m, 2H), 2.05 (m, 1H), 1,26 (m, 6H), 1.04 (d, 3H, $J$ = 6.8 Hz), 1.02 (d, 3H, $J$ = 6.8 Hz).

$^{13}$C NMR ($CDCl_3$, 150 MHz) δ 172.8 (s), 171.4 (s), 171.3 (s), 163.5 (s), 162.4 (d, $^1J_{C,F}$ = 248 Hz), 142.3 (s), 138.6 (s), 135.0 (s) 127.4 (d, $^3J_{C,F}$ = 9 Hz), 116.2 (d, $^2J_{C,F}$ = 23 Hz), 115.4 (s), 61.7 (s), 60.8 (s), 58.0 (s), 52.0 (s), 31.3 (s), 30.3 (s), 27.1 (s), 19.4 (s), 18.0 (s), 14.2 (s), 14.1 (s), 11.8 (s).

HRMS m/z: $[M+H]^+$ Calcd for $C_{25}H_{33}FN_4O_6$ 505.2457; Found 505.2455.

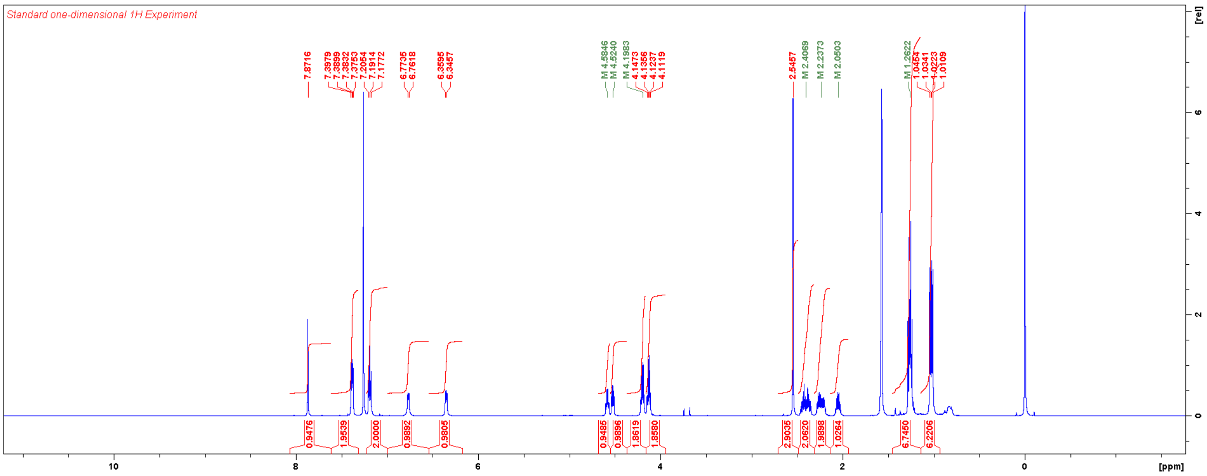


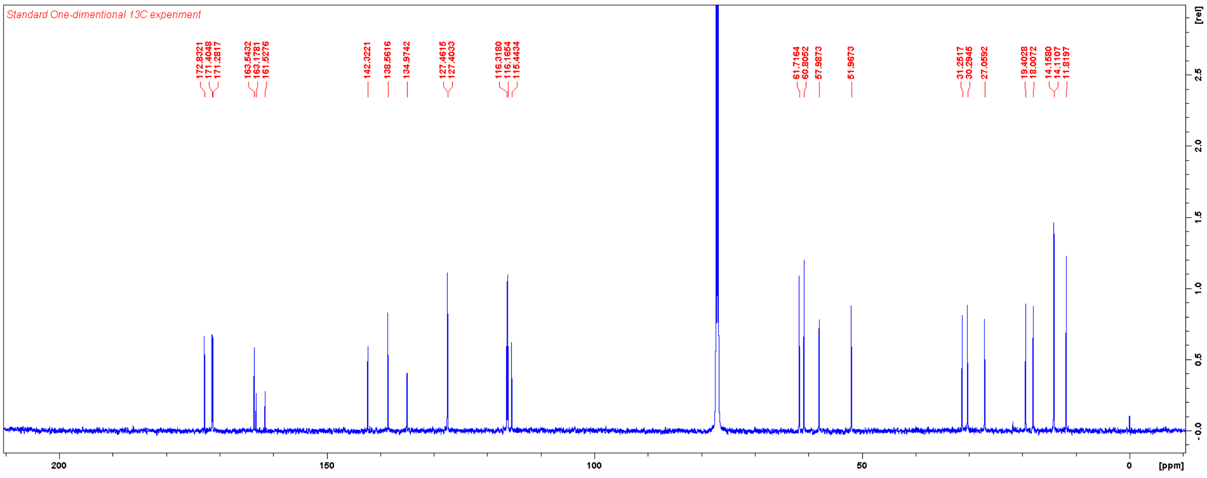

Diethyl (1-(4-fluorophenyl)-3-methyl-1*H*-pyrazole-4-carbonyl)-*L*-valyl-*D*-glutamate (**TT011**)

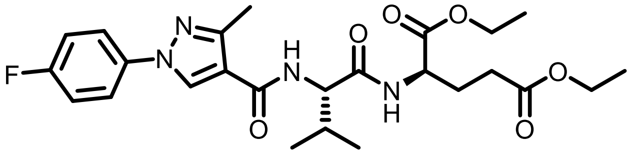

**TT011** was synthesised according to the general procedure for HATU-mediated coupling. Yield: 29 mg (0.057 mmol, 96%).

$^{1}$H NMR ($CDCl_3$, 600 MHz) δ 8.23 (s, 1H), 7.63 (dd, 2H, $J_{H,H}$ = 8.9 Hz, $^{4}J_{H,F}$ = 4.6 Hz), 7.15 (dd, 2H, $J_{H,H}$ = 8.5 Hz, $^{3}J_{H,F}$ = 8.5 Hz), 6.82 (d, 1H, $J$ = 6.7 Hz), 6.42 (d, 1H, $J$ = 7.8 Hz), 4.56 (m, 2H), 4.19 (q, 2H, $J$ = 7.1 Hz), 4.11 (m, 2H), 2.58 (s, 3H), 2.40 (m, 2H), 2.22 (m, 2H), 2.06 (m, 1H), 1,25 (m, 6H), 1.04 (d, 3H, $J$ = 6.8 Hz), 1.01 (d, 3H, $J$ = 6.8 Hz).

$^{13}$C NMR ($CDCl_3$, 150 MHz) δ 172.8 (s), 171.5 (s), 171.4 (s), 163.2 (s), 162.2 (d, $^{1}J_{C,F}$ = 245 Hz), 149.8 (s), 135.7 (s), 129.1 (s), 121.1 (d, $^{3}J_{C,F}$ = 8 Hz), 117.5 (s), 116.4 (d, $^{2}J_{C,F}$ = 23 Hz), 61.7 (s), 60.8 (s), 58.1 (s), 52.0 (s), 31.3 (s), 30.3 (s), 27.0 (s), 19.4 (s), 18.1 (s), 14.14 (s), 14.09 (s), 13.8 (s).

HRMS m/z: $[M+H]^+$ Calcd for $C_{25}H_{33}FN_4O_6$ 505.2457; Found 505.2457.

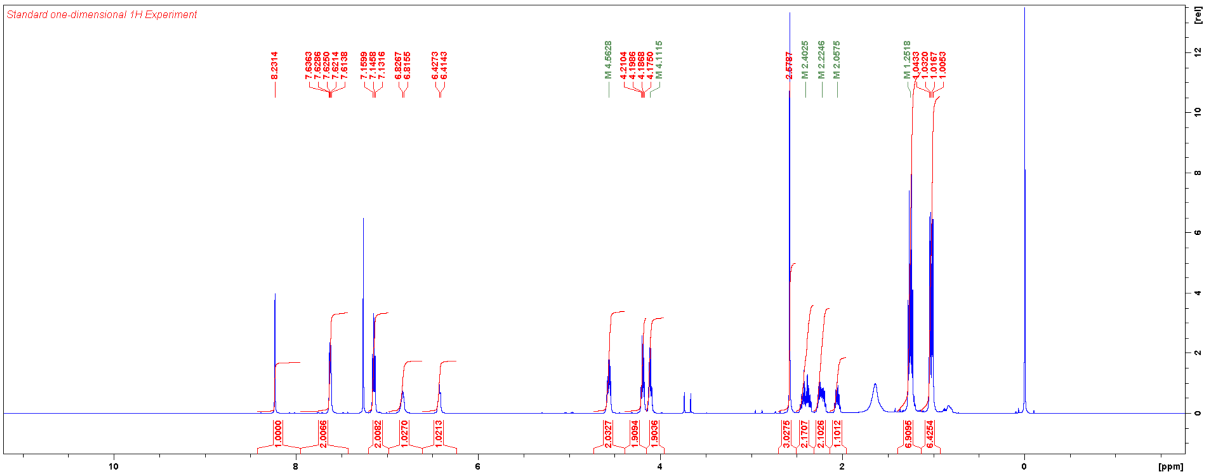


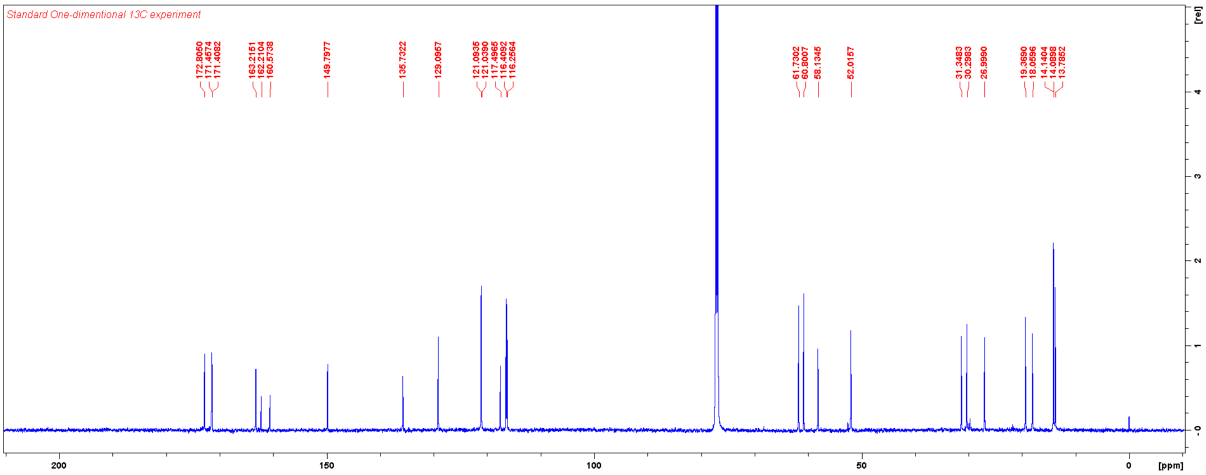

Diethyl (1-(4-fluorophenyl)-5-isopropyl-1*H*-pyrazole-4-carbonyl)-*L*-valyl-*D*-glutamate (**TT012**)

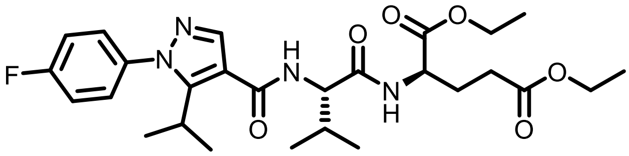

**TT012** was synthesised according to the general procedure for HATU-mediated coupling. Yield: 20 mg (0.038 mmol, 63%).

$^1$H NMR ($CDCl_3$, 600 MHz) δ 7.82 (s, 1H), 7.34 (dd, 2H, $J_{H,H}$ = 8.8 Hz, $^4J_{H,F}$ = 4.7 Hz), 7.19 (dd, 2H, $J_{H,H}$ = 8.8 Hz, $^3J_{H,F}$ = 8.5 Hz), 6.78 (d, 1H, $J$ = 6.4 Hz), 6.37 (d, 1H, $J$ = 8.2 Hz), 4.59 (m, 1H), 4.51 (dd, 1H, $J$ = 8.4, 6.0 Hz), 4.20 (m, 2H), 4.13 (q, 2H, $J$ = 7.1 Hz), 3.22 (m, 1H), 2.41 (m, 2H), 2.27 (m, 1H), 2.21 (m, 1H), 2.05 (m, 1H), 1.33 (d, 6H, $J$ = 7.1 Hz), 1.27 (m, 6H), 1.04 (d, 3H, $J$ = 6.7 Hz), 1.02 (d, 3H, $J$ = 6.8 Hz).

$^{13}$C NMR ($CDCl_3$, 150 MHz) δ 172.8 (s), 171.5 (s), 171.3 (s), 163.7(s), 162.7 (d, $^1J_{C,F}$ = 248 Hz), 151.5 (s), 139.2 (s), 135.7 (s), 128.6 (d, $^3J_{C,F}$ = 9 Hz), 116.2 (d, $^2J_{C,F}$ = 23 Hz), 114.8 (s), 61.7 (s), 60.8 (s), 58.3 (s), 51.9 (s), 31.1 (s), 30.3 (s), 27.1 (s), 26.3 (s), 20.7 (s), 20.7 (s), 19.4 (s), 18.0 (s), 14.2 (s), 14.1 (s).

HRMS m/z: $[M+H]^+$ Calcd for $C_{27}H_{37}FN_4O_6$ 533.2770; Found 533.2768.

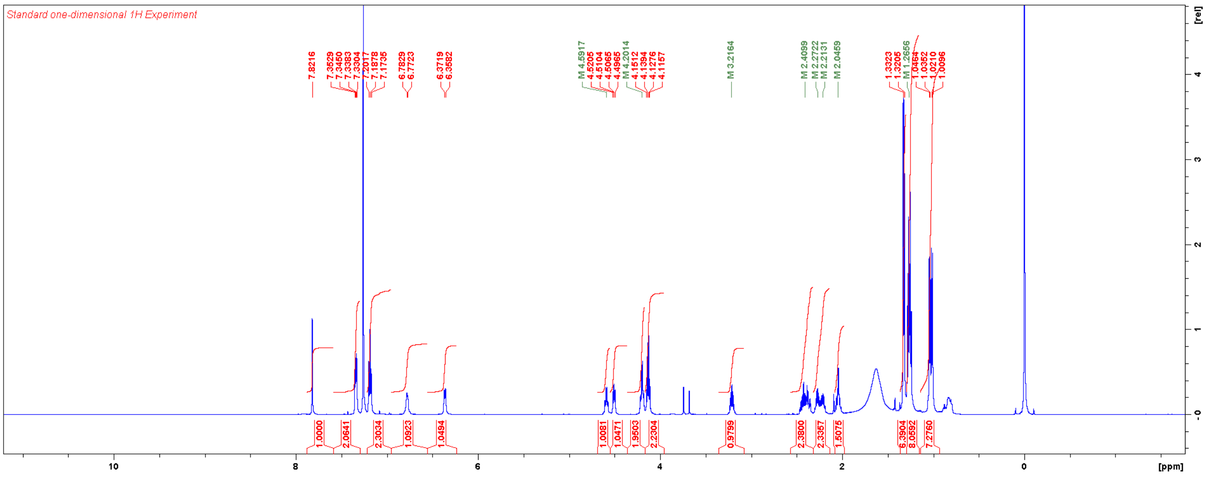


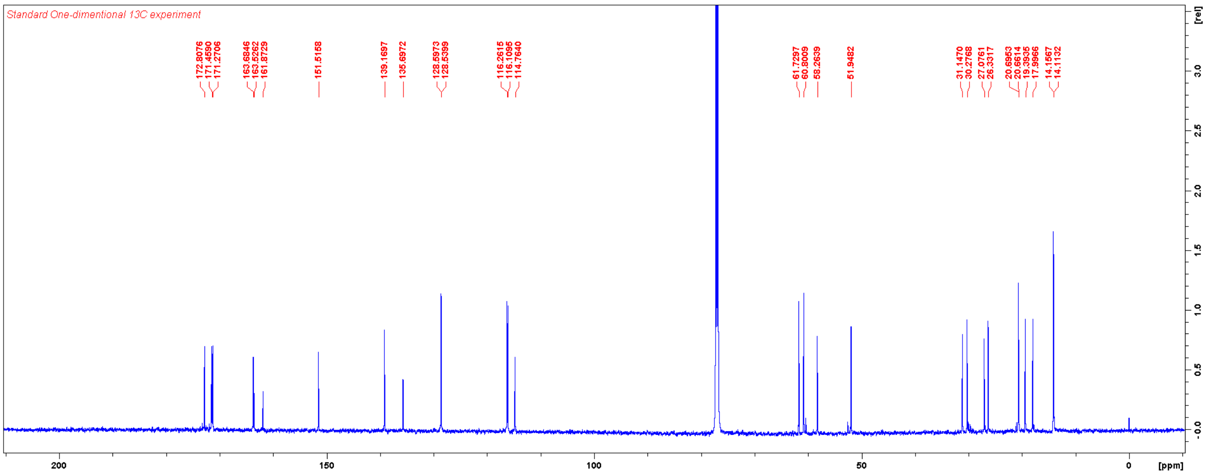

Diethyl (3-methyl-1-(p-tolyl)-1*H*-pyrazole-4-carbonyl)-*L*-valyl-*D*-glutamate (**TT013**)

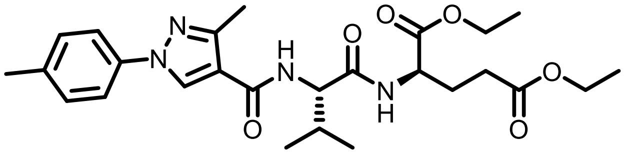

**TT013** was synthesised according to the general procedure for HATU-mediated coupling. Yield: 22 mg (0.044 mmol, 73%).

$^1$H NMR ($CDCl_3$, 600 MHz) δ 7.87 (s, 1H), 7.28 (m, 4H), 6.77 (d, 1H, $J$ = 7.2 Hz), 6.32 (d, 1H, $J$ = 8.3 Hz), 4.58 (dd, 1H, $J$ = 12.7, 7.4 Hz), 4.52 (dd, 1H, $J$ = 7.8, 6.3 Hz), 4.20 (q, 2H, $J$ = 7.0 Hz), 4.12 (q, 2H, $J$ = 7.0 Hz), 2.55 (s, 3H), 2.42 (m, 5H), 2.24 (m, 2H), 2.06 (m, 1H), 1.26 (m, 6H), 1.04 (d, 3H, $J$ = 6.7 Hz), 1.02 (d, 3H, $J$ = 6.8 Hz).

$^{13}$C NMR ($CDCl_3$, 150 MHz) δ 172.8, 171.4, 171.3, 163.7, 142.2, 138.7, 138.3, 136.4, 129.8, 125.3, 115.2, 61.7, 60.8, 58.0, 52.0, 31.2, 30.3, 27.1, 21.2, 19.4, 18.0, 14.2, 14.1, 11.9.

HRMS m/z: $[M+H]^+$ Calcd for $C_{26}H_{36}N_4O_6$ 501.2708; Found 501.2707.

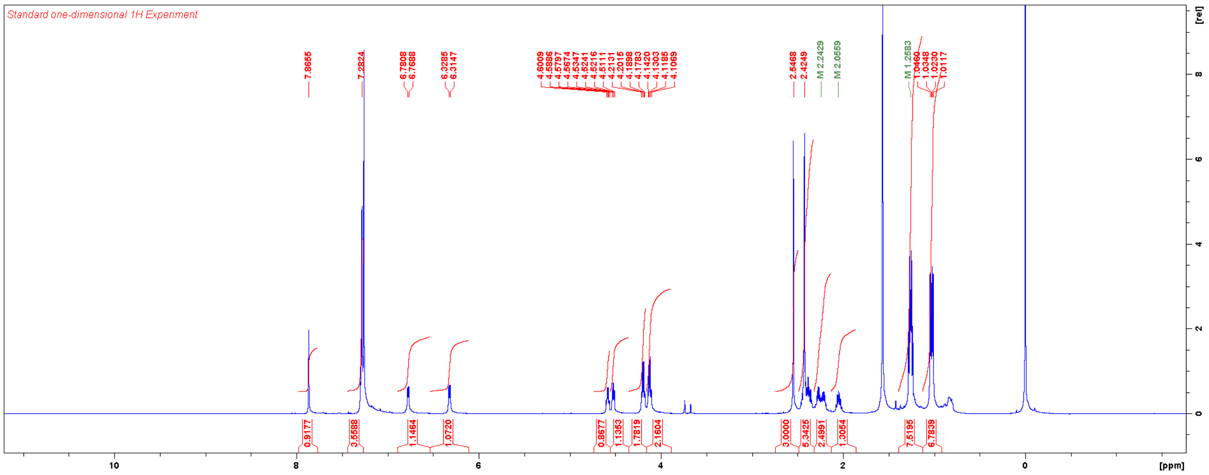


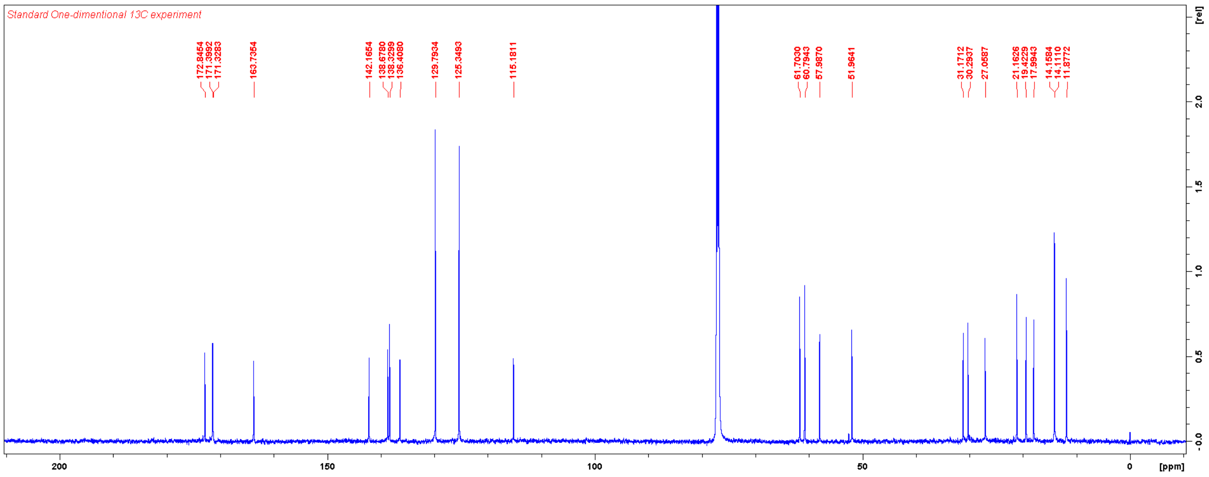

Diethyl (1-(2-hydroxyphenyl)-5-methyl-1*H*-pyrazole-4-carbonyl)-*L*-valyl-*D*-glutamate (**TT014**)

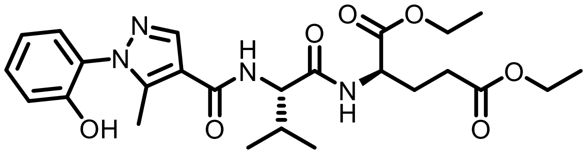

To a solution of **TT015** (20 mg, 0.039 mmol) in DCM was added 1M $BBr_3$ / DCM (78 µL, 0.078 mmol) on ice. The reaction mixture was stirred on ice for 2 h and then at RT for 1 h. The reaction mixture was directly loaded on column and purified with flash chromatography using a gradient of MeOH in DCM. Yield: 19 mg (0.038 mmol, 98%).

$^1$H NMR ($CDCl_3$, 600 MHz) δ 8.61 (br, 1H), 7.96 (s, 1H), 7.29 (m, 1H), 7.20 (dd, 1H, *J* = 7.9, 0.8 Hz), 7.10 (d, 1H, *J* = 7.9 Hz), 7.02 (d, 1H, *J* = 7.4 Hz), 6.97 (m, 1H), 6.59 (d, 1H, *J* = 8.5 Hz), 4.57 (m, 1H), 4.53 (dd, 1H, *J* = 8.3, 6.4 Hz), 4.18 (q, 2H, *J* = 7.2 Hz), 4.11 (q, 2H, *J* = 6.9 Hz), 2.60 (s, 3H), 2.40 (m, 2H), 2.22 (m, 2H), 2.04 (m, 1H), 1.25 (m, 6H), 1.02 (d, 3H, *J* = 6.7 Hz), 1.00 (d, 3H, *J* = 6.8 Hz).

$^{13}$C NMR ($CDCl_3$, 150 MHz) δ 173.0, 171.5, 171.4, 163.3, 151.2, 143.4, 139.0, 130.0, 125.0, 120.0, 118.6, 115.7, 61.8, 60.9, 58.1, 52.0, 31.3, 30.3, 26.9, 19.4, 18.1, 14.19, 14.16, 12.2.

HRMS m/z: $[M+H]^+$ Calcd for $C_{25}H_{34}N_4O_7$ 503.2500; Found 503.2499.

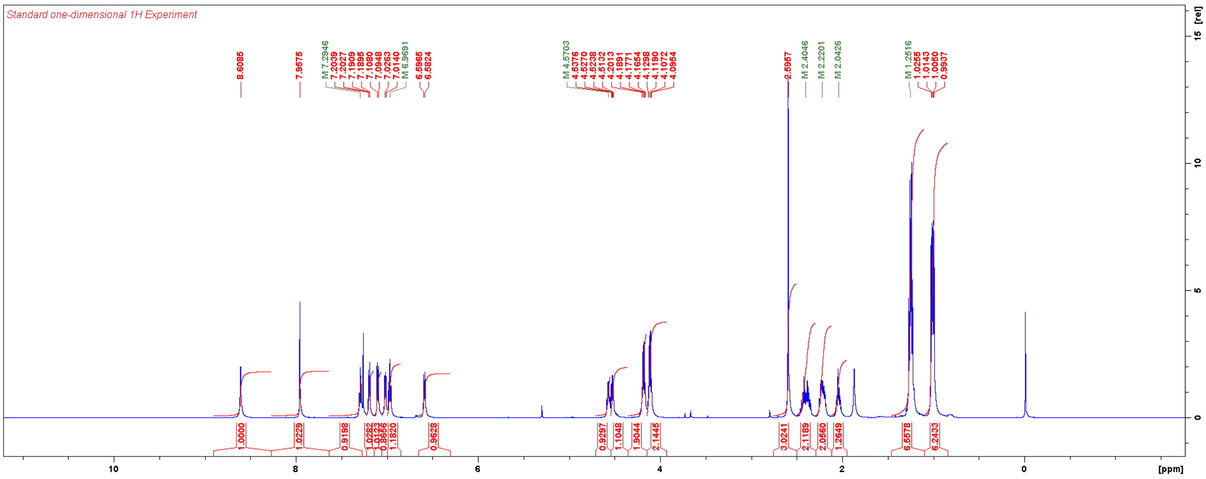


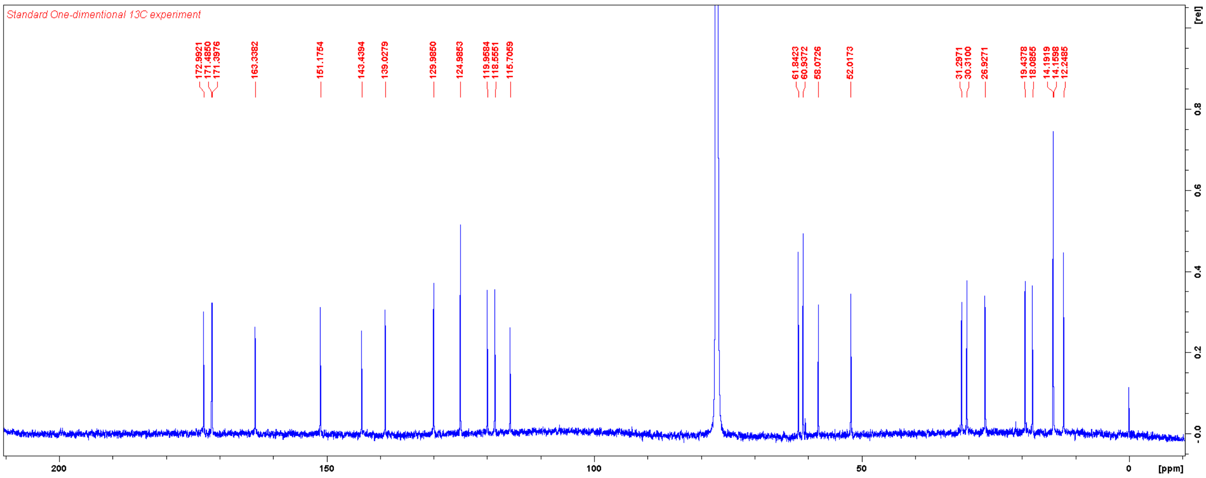

Diethyl (1-(2-methoxyphenyl)-5-methyl-1*H*-pyrazole-4-carbonyl)-*L*-valyl-*D*-glutamate (**TT015**)

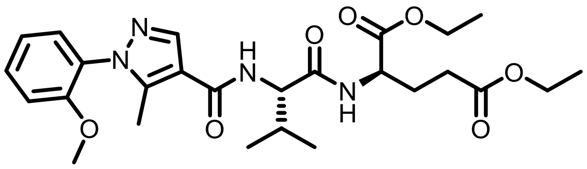

**TT015** was synthesised according to the general procedure for HATU-mediated coupling. Yield: 66 mg (0.13 mmol, 71%).

$^{1}$H NMR ($CDCl_3$, 600 MHz) δ 7.90 (s, 1H), 7.45 (m, 1H), 7.30 (m, 1H), 7.06 (m, 2H), 7.78 (d, 1H, *J* = 6.8 Hz), 6.32 (d, 1H, *J* = 8.2 Hz), 4.59 (m, 1H), 4.53 (m, 1H), 4.20 (m, 2H), 4.13 (m, 2H), 3.80 (s, 3H), 2.42 (m, 2H), 2.38 (s, 3H), 2.28 (m, 1H), 2.22 (m, 1H), 2.06 (m, 1H), 1.26 (m, 6H), 1.04 (d, 3H, *J* = 6.8 Hz), 1.02 (d, 3H, *J* = 6.8 Hz).

$^{13}$C NMR ($CDCl_3$, 150 MHz) δ 172.9, 171.5, 163.9, 154.6, 144.2, 138.7, 130.9, 129.0, 127.6, 121.0, 114.5, 112.1, 61.7, 60.8, 58.1, 55.8, 52.0, 31.3, 30.4, 27.1, 19.5, 18.1, 14.24, 14.18, 11.2.

HRMS m/z: $[M+H]^+$ Calcd for $C_{26}H_{36}N_4O_7$ 517.2657; Found 517.2654.

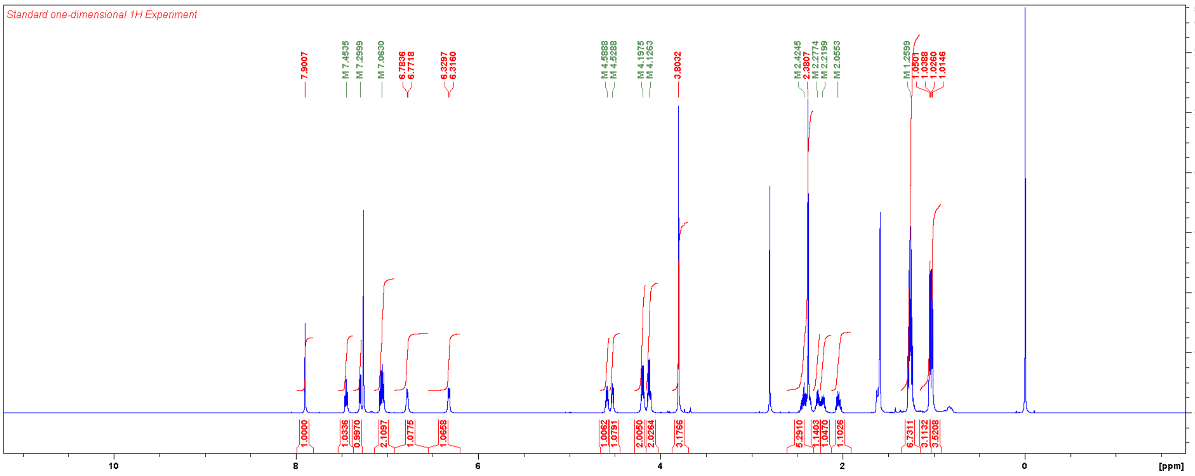


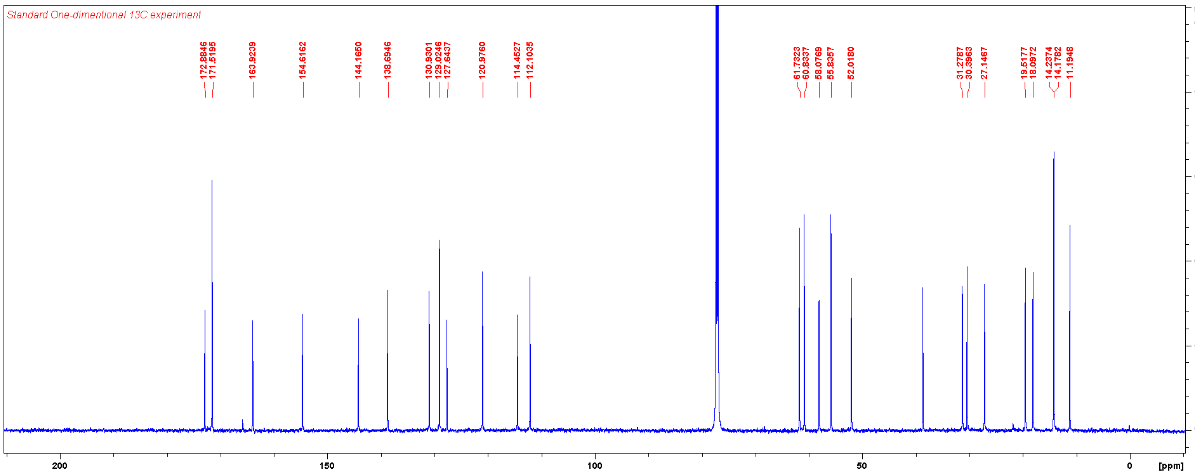

Diethyl (5-methyl-1-(pyridin-4-yl)-1*H*-pyrazole-4-carbonyl)-*L*-valyl-*D*-glutamate (**TT016**)

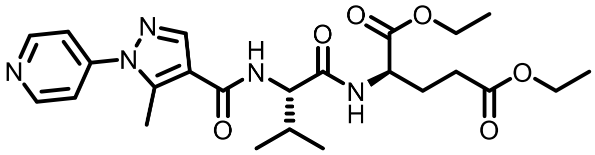

**TT016** was synthesised according to the general procedure for HATU-mediated coupling. Yield: 12 mg (0.025 mmol, 41%).

$^{1}$H NMR ($CDCl_3$, 600 MHz) δ 8.76 (d, 2H, *J* = 5.3 Hz), 7.93 (s, 1H), 7.46 (d, 2H, *J* = 6.0 Hz), 6.76 (d, 1H, *J* = 5.7 Hz), 6.40 (d, 1H, *J* = 7.6 Hz), 4.58 (m, 1H), 4.53 (dd, 1H, *J* = 8.3, 6.0 Hz), 4.20 (m, 2H), 4.13 (q, 2H, *J* = 6.9 Hz), 2.41 (m, 2H), 2.24 (m, 2H), 2.06 (m, 1H), 1.27 (m, 6H), 1.04 (d, 3H, *J* = 6.7 Hz), 1.02 (d, 3H, *J* = 6.8 Hz).

$^{13}$C NMR ($CDCl_3$, 150 MHz) δ 172.9, 171.4, 171.2, 163.1, 151.1, 145.8., 142.5, 139.7, 118.6, 116.9, 61.8, 60.8, 58.0, 52.0, 31.3, 30.3, 27.0, 19.4, 18.0, 14.2, 14.1, 12.3.

HRMS m/z: [M+H]$^{+}$ Calcd for $C_{24}H_{33}N_5O_6$ 488.2504; Found 488.2500.

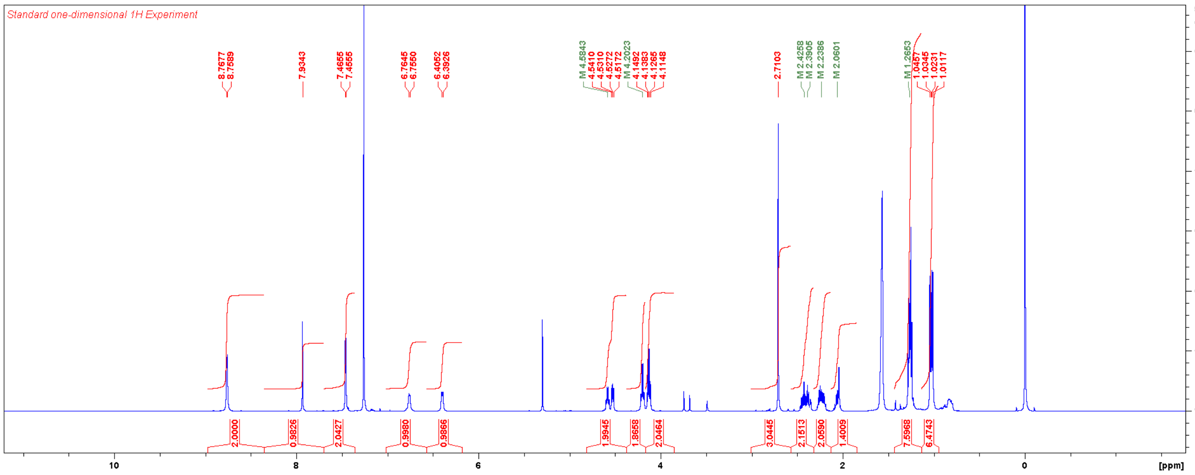


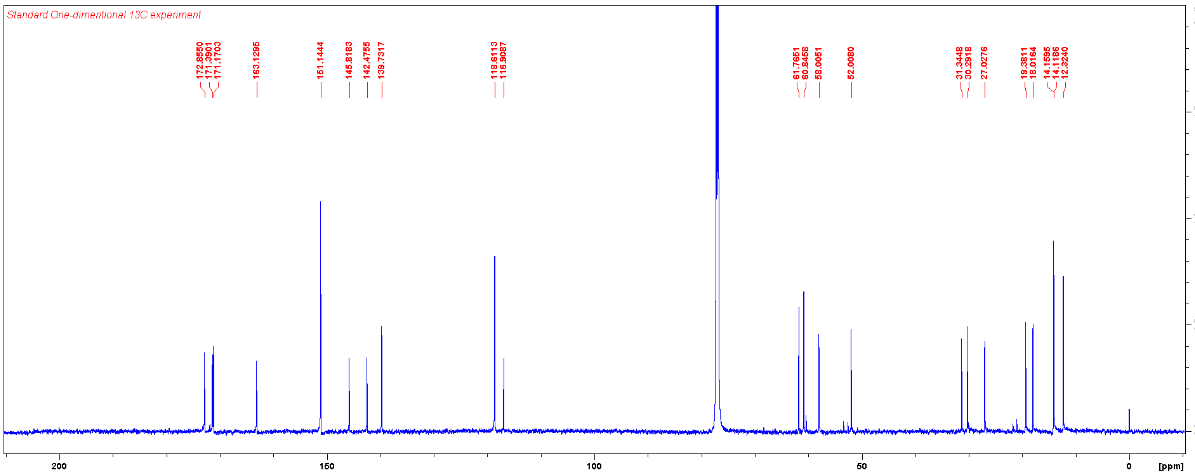

Diethyl (5-methyl-1-(pyridin-2-yl)-1*H*-pyrazole-4-carbonyl)-*L*-valyl-*D*-glutamate (**TT017**)

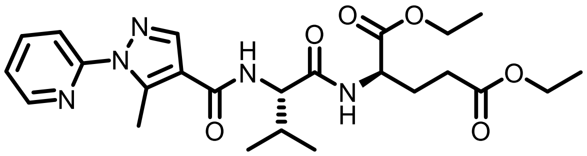

**TT017** was synthesised according to the general procedure for HATU-mediated coupling. Yield: 27 mg (0.055 mmol, 92%).

$^1$H NMR ($CDCl_3$, 600 MHz) δ 8.50 (s, 1H), 7.89 (s, 1H), 7.86 (m, 1H), 7.80 (m, 1H), 7.29 (m, 1H), 6.79 (d, 1H, *J* = 7.3 Hz), 6.34 (d, *J* = 8.2 Hz), 4.59 (m, 1H), 4.53 (m, 1H), 4.20 (q, 2H, *J* = 6.7 Hz), 4.12 (q, 2H, *J* = 7.1 Hz), 2.90 (s, 3H), 2.50–2.00 (m, 5H), 1.26 (m, 6H), 1.04 (d, 3H, *J* = 6.8 Hz), 1.02 (d, 3H, *J* = 6.8 Hz).

$^{13}$C NMR ($CDCl_3$, 150 MHz) δ 173.0, 171.6, 171.4, 163.7, 153.0, 148.0, 143.6, 139.4, 138.7, 122.6, 117.9, 116.8, 61.9, 60.9, 58.2, 52.1, 31.3, 30.4, 27.1, 19.5, 18.2, 14.3, 14.2, 13.1.

HRMS m/z: [M+H]$^+$ Calcd for $C_{24}H_{33}N_5O_6$ 488.2504; Found 488.2502.

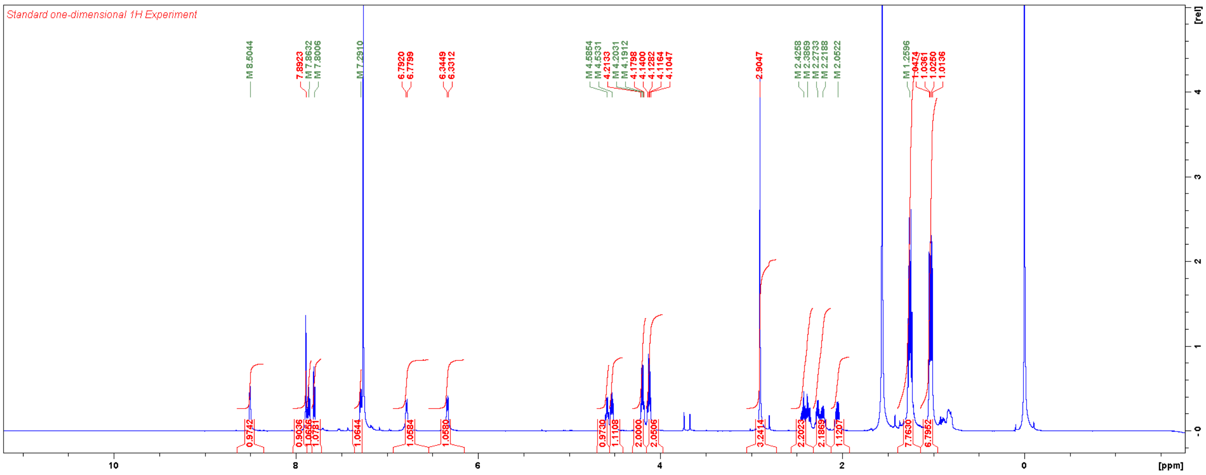


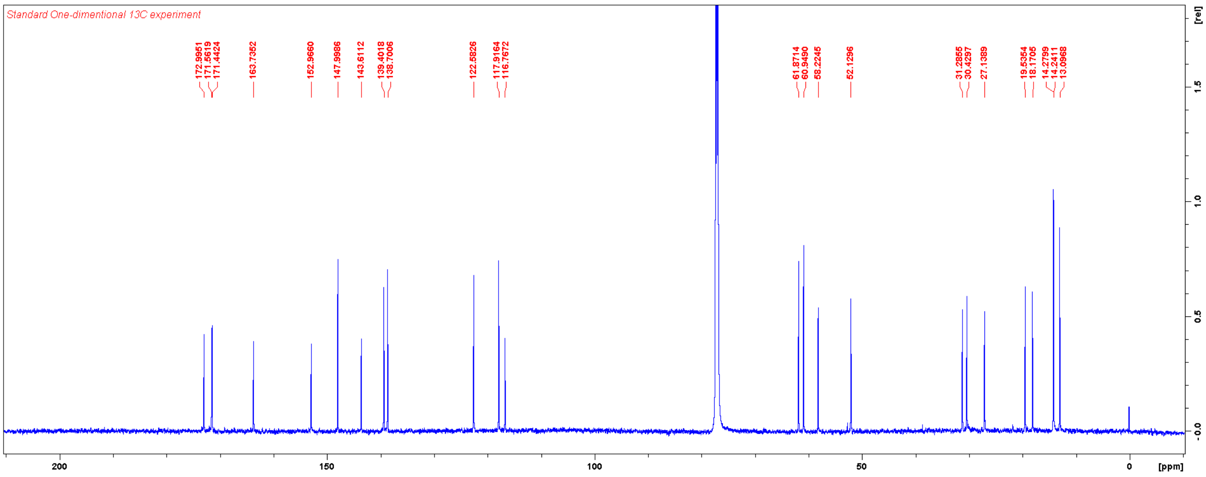

Diethyl (3,5-dimethyl-1-phenyl-1*H*-pyrazole-4-carbonyl)-*L*-valyl-*D*-glutamate (**TT018**)

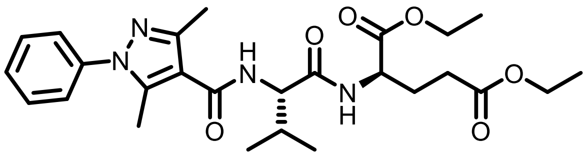

**TT018** was synthesised according to the general procedure for HATU-mediated coupling. Yield: 24 mg (0.048 mmol, 80%).

$^1$H NMR ($CDCl_3$, 600 MHz) δ 7.58–7.31 (m, 5H), 6.75 (br, 1H), 6.29 (d, 1H, *J* = 8.0 Hz), 4.58 (m, 2H), 4.20 (m, 2H), 4.12 (m, 2H), 2.54 (s, 3H), 2.50 (s, 3H), 2.41 (m, 2H), 2.25 (m, 2H), 2.05 (m, 1H), 1.26 (m, 6H), 1.06 (d, 3H, *J* = 6.8 Hz), 1.03 (d, 3H, *J* = 6.9 Hz).

$^{13}$C NMR ($CDCl_3$, 150 MHz) δ 172.9, 171.5, 171.4, 164.7, 147.5, 142.5, 139.0, 129.3, 128.5, 125.7, 114.5, 61.8, 60.9, 58.1, 52.1, 31.4, 30.4, 27.2, 19.6, 18.1, 14.4, 14.3, 14.2, 12.5.

HRMS m/z: $[M+H]^+$ Calcd for $C_{26}H_{36}N_4O_6$ 501.2708; Found 501.2703.

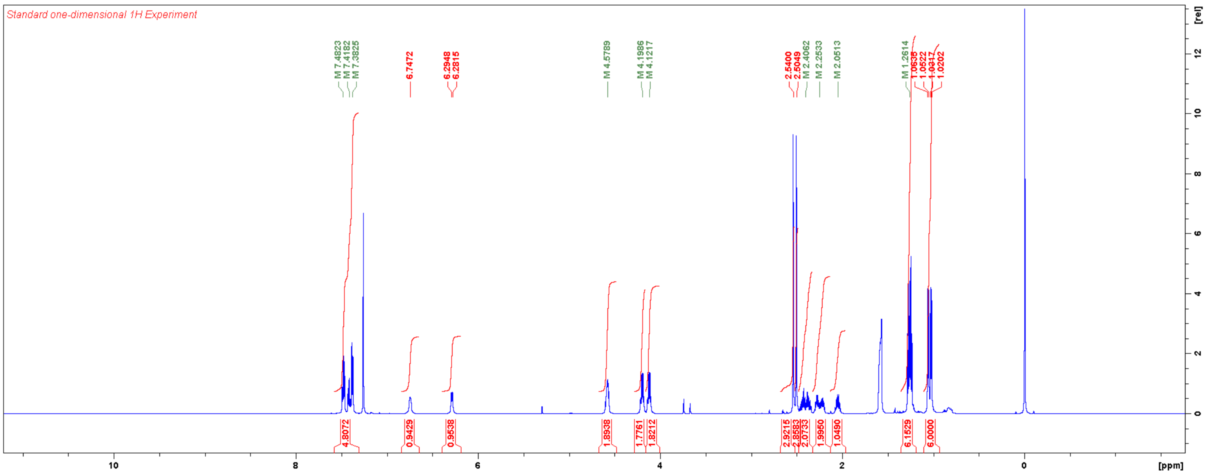


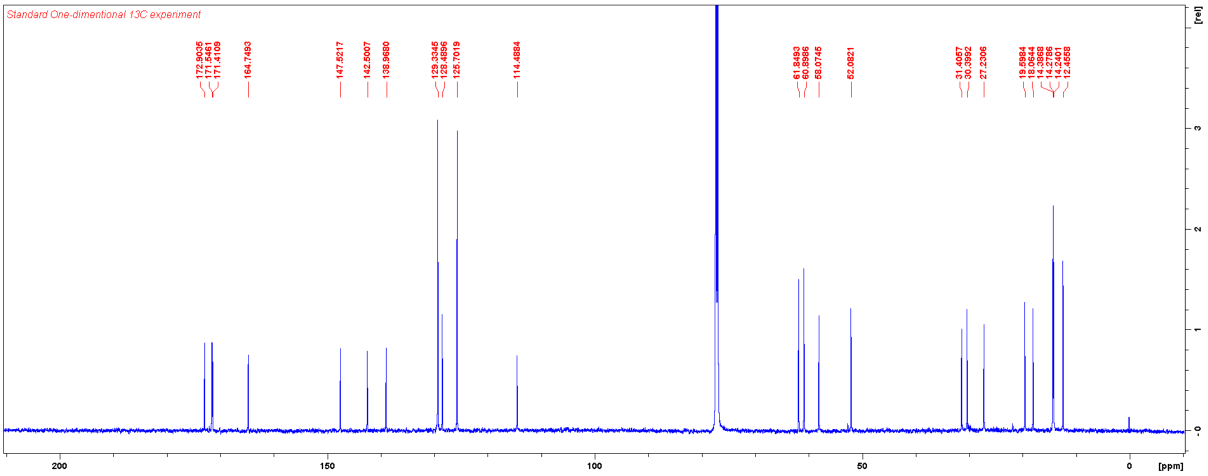

Diethyl *N*-(3,5-dimethyl-1-phenyl-1*H*-pyrazole-4-carbonyl)-*N*-methyl-*L*-valyl-*D*-glutamate (**TT019**)

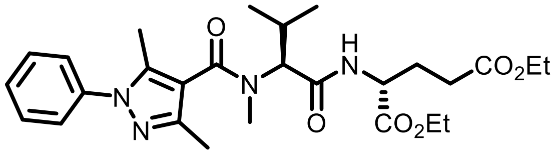

To a solution of 3,5-dimethyl-1-phenyl-1*H*-pyrazole-4-carboxylic acid (22 mg, 0.10 mmol) and DMF (2 drops) in DCM (1.0 mL) was added dropwise oxalyl chloride (43 µL, 0.50 mmol) at 0 °C. The reaction mixture was stirred for 1 h and concentrated under vacuum to remove excess oxalyl chloride. This material was used without further purification. To a solution of the dipeptide (0.080 mmol) and DIPEA (70 µL, 0.40 mmol) in DCM (0.5 mL) was added dropwise the acyl chloride in DCM (0.5 mL) at 0 °C. The reaction mixture was allowed to warm to RT over 2 days. Upon completion of the reaction, the reaction mixture was directly loaded on column and purified with flash chromatography using a gradient of EtOAc in hexanes. Yield: 16 mg (0.040 mmol, 40% over two steps).

$^1$H NMR ($CDCl_3$, 600 MHz) δ 7.44 (m, 5H), 7.24 (br, 1H), 4.67 (d, 1H, *J* = 11.3 hz), 4.52 (m, 1H), 4.14 (m, 4H), 2.93 (s, 3H), 2.42 (m, 3H), 2.33 (m, 6H), 2.22 (m, 1H), 1.98 (m, 1H), 1.25 (m, 6H), 1.04 (d, 3H, *J* = 6.4 Hz), 0.98 (d, 3H, *J* = 6.7 Hz).

$^{13}$C NMR ($CDCl_3$, 150 MHz) δ 172.4, 171.6, 170.2, 169.1, 139.0, 129.2, 128.1, 125.2, 115.7, 62.7, 61.4, 60.7, 51.6, 32.5, 30.6, 26.9, 24.9, 20.0, 18.7, 14.19, 14.16, 12.9, 11.6.

HRMS m/z: $[M+H]^+$ Calcd for $C_{27}H_{38}N_4O_6$ 515.2864; Found 515.2859.

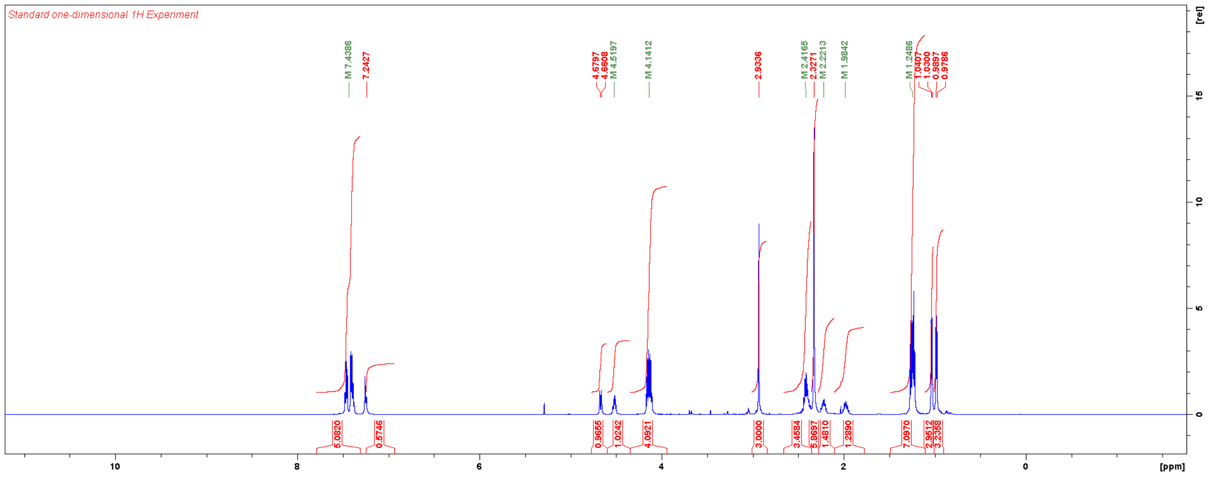

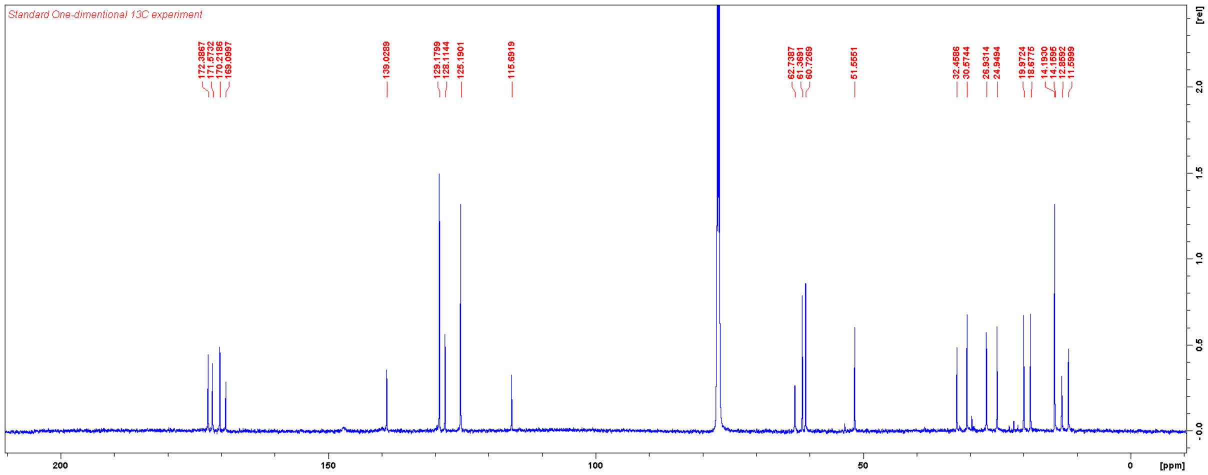
Standard One-dimentional 13C experiment
172.3867
171.5732
170.2186
169.0997
139.0289
129.1799
128.1144
125.1901
115.6919
62.7387
61.3691
60.7269
51.5551
32.4586
30.5744
26.9314
24.9494
19.9724
18.6775
14.1930
14.1595
12.8592
11.5999
[rel]
[ppm]

Diethyl *N*-(1-(4-fluorophenyl)-5-methyl-1*H*-pyrazole-4-carbonyl)-*N*-methyl-*L*-valyl-*D*-glutamate (**TT020**)

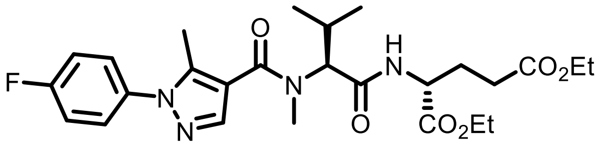

**TT020** was synthesised according to the general procedure for HATU-mediated coupling. Yield: 16 mg (0.031 mmol, 31%).

$^1$H NMR ($CDCl_3$, 600 MHz) δ 7.71 (s, 1H), 7.41 (m, 2H), 7.19 (m, 3H), 4.63 (d, 1H, $J$ = 11.2 Hz), 4.54 (m, 1H), 4.14 (m, 4H), 3.07 (s, 3H), 2.44 (s, 3H), 2.40 (m, 3H), 2.22 (m, 1H), 1.97 (m, 1H), 1.25 (m, 6H), 1.03 (d, 3H, $J$ = 6.3 Hz) 0.97 (d, 3H, $J$ = 6.6 Hz).

$^{13}$C NMR ($CDCl_3$, 150 MHz) δ 172.5 (s), 171.6 (s), 170.3 (s), 167.7 (s), 162.3 (d, $^1J_{C,F}$ = 247 Hz), 142.1 (s), 139.3 (s), 135.1 (s), 127.3 (d, $^3J_{C,F}$ = 8.7 Hz), 116.2 (d, $^2J_{C,F}$ = 23 Hz), 115.6 (s), 62.9 (s), 61.4 (s), 60.7 (s), 51.4 (s), 33.3 (s), 30.5 (s), 27.0 (s), 25.2 (s), 19.9 (s), 18.7 (s), 14.2 (s), 14.1 (s), 11.7 (s).

HRMS m/z: $[M+H]^+$ Calcd for $C_{26}H_{35}FN_4O_6$ 519.2613; Found 519.2613.

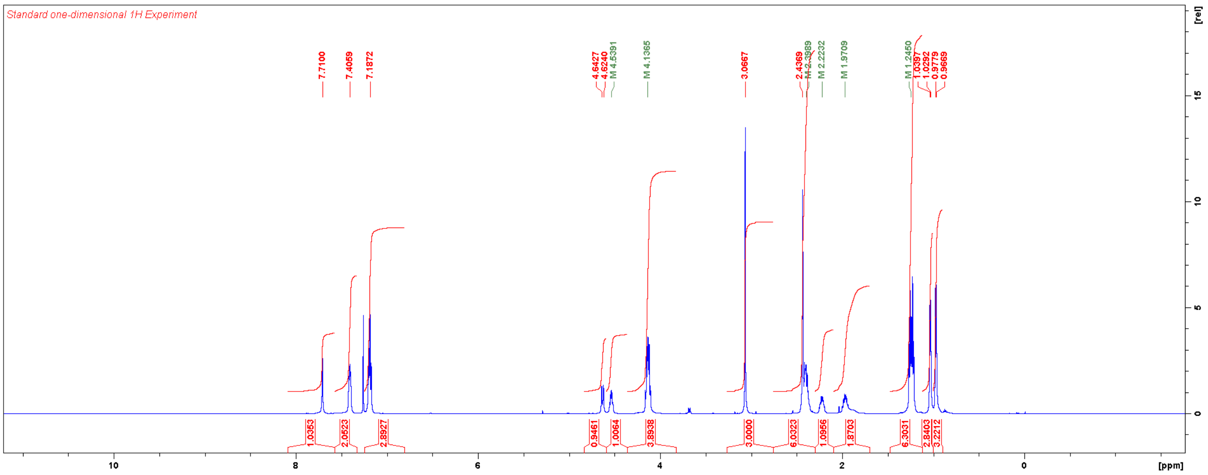


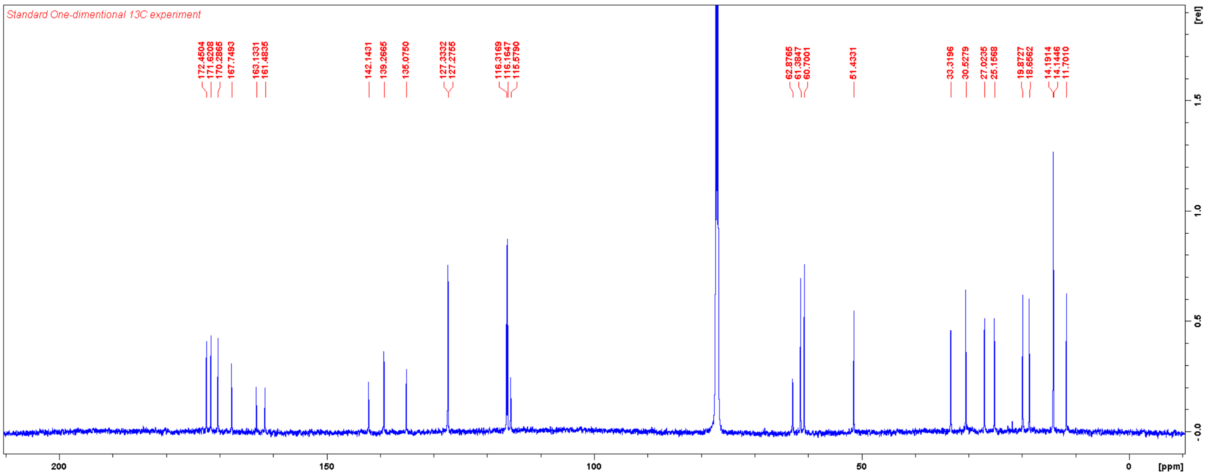

Diethyl *N*-(tert-butoxycarbonyl)-*N*-methyl-*L*-valyl-*D*-glutamate

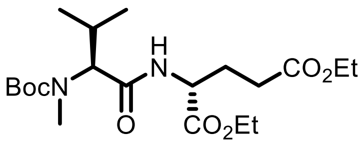


To a solution of Boc-*N*-methyl-*L*-valine-OH (386 mg, 1.67 mmol), diethyl *D*-glutamate hydrochloride (600 mg, 2.50 mmol), EDC-HCl (320 mg, 1.67 mmol) and HOBt-x$H_2O$ (248 mg, 1.84 mmol) in DMF (10 mL) was added TEA (0.70 mL, 5.01 mmol) at 0 °C. The reaction mixture was allowed to warm to RT overnight. The reaction mixture was diluted with EtOAc, washed with aqueous $NaHCO_3$, aqueous $NH_4Cl$, and brine, dried over $MgSO_4$, filtered, and concentrated. The residue was purified by flash chromatography using a gradient of EtOAc in hexanes. Yield: 388 mg (0.93 mmol, 56%), colourless oil.

$^1$H NMR ($CDCl_3$, 600 MHz) δ 6.77 (br, 1H), 4.51 (m, 1H), 4.13 (m, 4H), 2.76 (s, 3H), 2.36 (m, 2H), 2.24 (m, 2H), 1.96 (m, 1H), 1.48 (s, 9H), 1.25 (m, 6H), 0.91 (d, 3H, $J$ = 6.4 Hz), 0.87 (d, 3H, $J$ = 6.4 Hz).

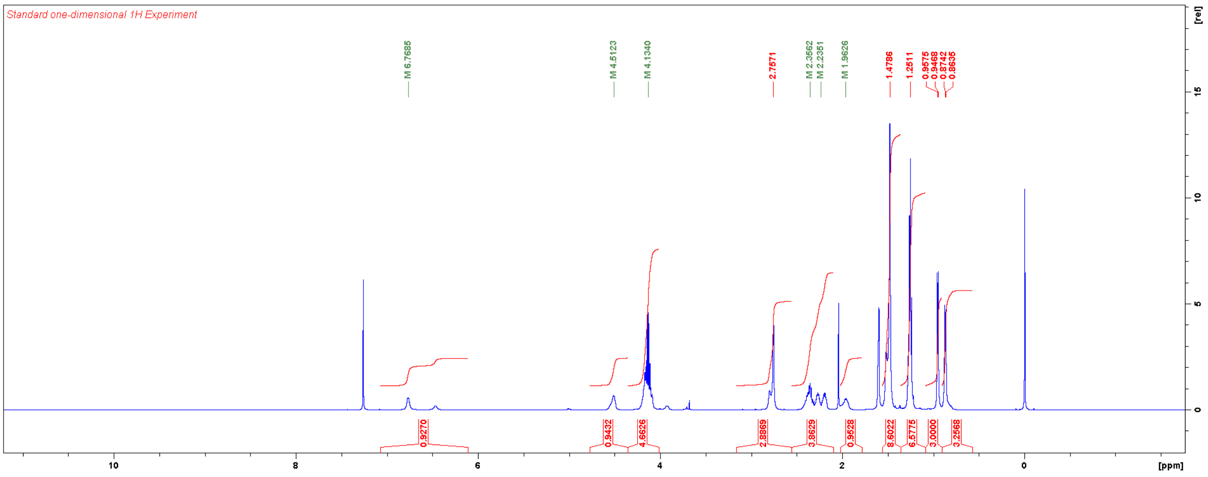

Diethyl *N*-(1-(4-fluorophenyl)-3-methyl-1*H*-pyrazole-4-carbonyl)-*N*-methyl-*L*-valyl-*D*-glutamate (**TT021**)

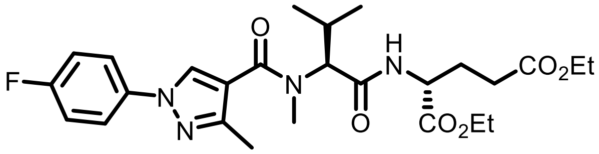


**TT021** was synthesised according to the general procedure for HATU-mediated coupling. Yield: 19 mg (0.037 mmol, 37%).

$^1$H NMR ($CDCl_3$, 600 MHz) δ 7.98 (s, 1H), 7.61 (m, 2H), 7.14 (m, 3H), 4.66 (d, 1H, $J$ = 11.1 Hz), 4.53 (m, 1H), 4.14 (m, 4H), 2.99 (s, 3H), 2.43 (s, 3H), 2.38 (m, 1H), 2.23 (m, 1H), 1.99 (m, 1H), 1.25 (m, 6H), 1.03 (d, 3H, $J$ = 6.2 Hz), 0.97 (d, 3H, $J$ = 6.5 Hz).

$^{13}$C NMR ($CDCl_3$, 150 MHz) δ 172.4 (s), 171.6 (s), 170.2 (s), 167.6 (s), 161.4 (d, $^1J_{C,F}$ = 245 Hz), 150.0 (s), 135.9 (s), 128.1 (s), 121.1 (d, $^3J_{C,F}$ = 8 Hz), 117.3 (s), 116.3 (d, $^2J_{C,F}$ = 23 Hz), 62.9 (s), 61.4 (s), 60.7 (s), 51.5 (s), 33.1 (s), 30.5 (s), 26.9 (s), 25.1 (s), 19.9 (s), 18.6 (s), 14.2 (s), 13.1 (s).

HRMS m/z: $[M+H]^+$ Calcd for $C_{26}H_{35}FN_4O_6$ 519.2613; Found 519.2614.

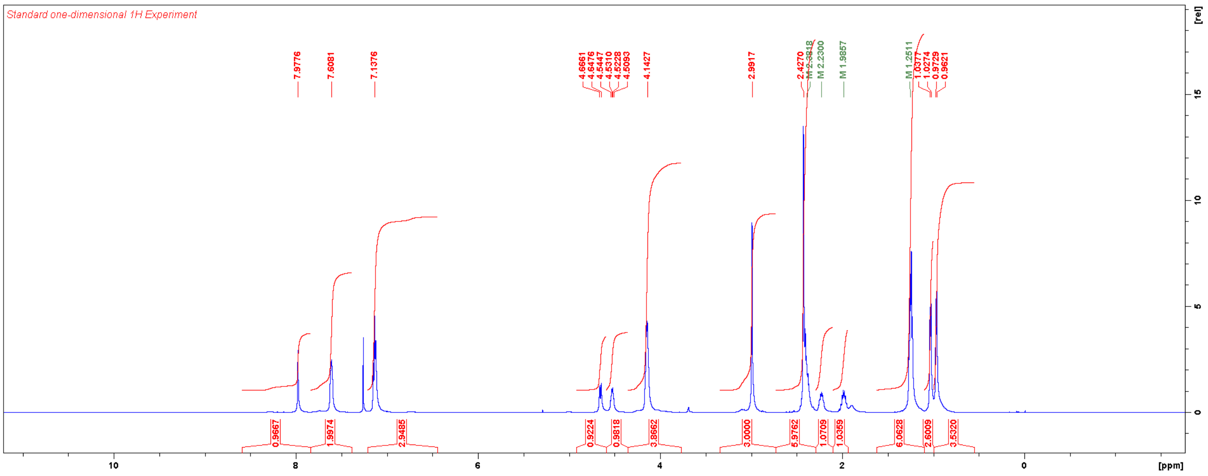


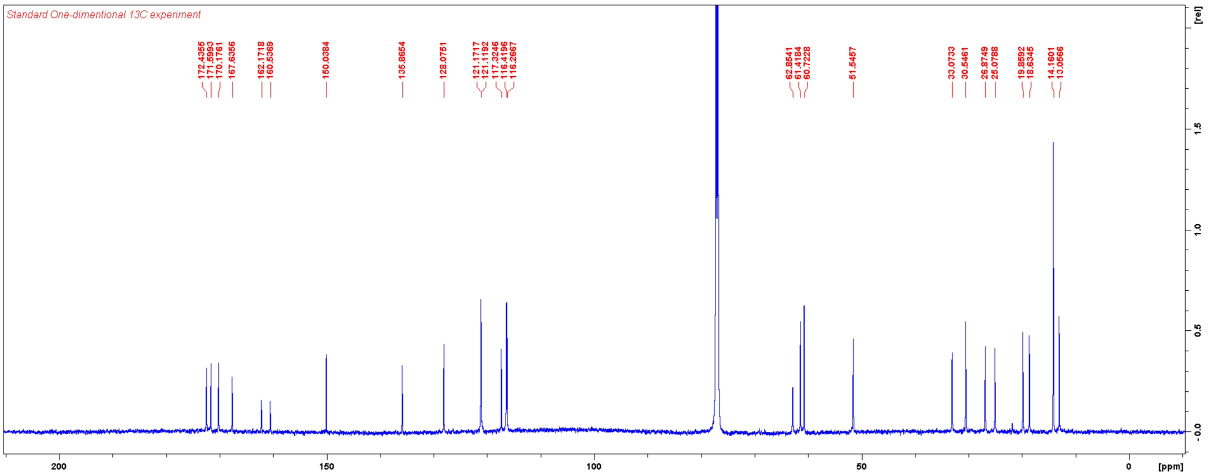

Diethyl (1-(3,4-dichlorophenyl)-3,5-dimethyl-1*H*-pyrazole-4-carbonyl)-*L*-valyl-*D*-glutamate (**TT022**)

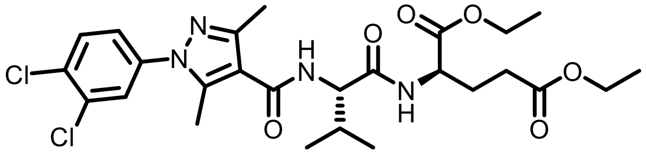

**TT022** was synthesised according to the general procedure for HATU-mediated coupling. Yield: 25 mg (0.044 mmol, 73%).

$^1$H NMR ($CDCl_3$, 600 MHz) δ 7.56 (m, 2H), 6.71 (d, 1H, *J* = 7.2 Hz), 6.30 (d, 1H, *J* = 8.1 Hz), 4.57 (m, 2H), 4.20 (m, 2H), 4.13 (m, 2H), 2.53 (s, 3H), 2.52 (s, 3H), 2.40 (m, 2H), 2.25 (m, 2H), 2.06 (m, 1H), 1.26 (m, 6H), 1.05 (d, 3H, *J* = 6.8 Hz), 1.02 (d, 3H, *J* = 6.8 Hz).

$^{13}$C NMR ($CDCl_3$, 150 MHz) δ 172.8, 171.4, 171.2, 164.2, 148.1, 142.4, 138.1, 133.3, 132.5, 130.8, 127.3, 124.4, 115.2, 61.8, 60.8, 57.9, 52.0, 31.4, 30.3, 27.1, 19.4, 17.9, 14.2, 14.1, 12.4.

HRMS m/z: $[M+H]^+$ Calcd for $C_{26}H_{34}Cl_2N_4O_6$ 569.1928; Found 569.1928.

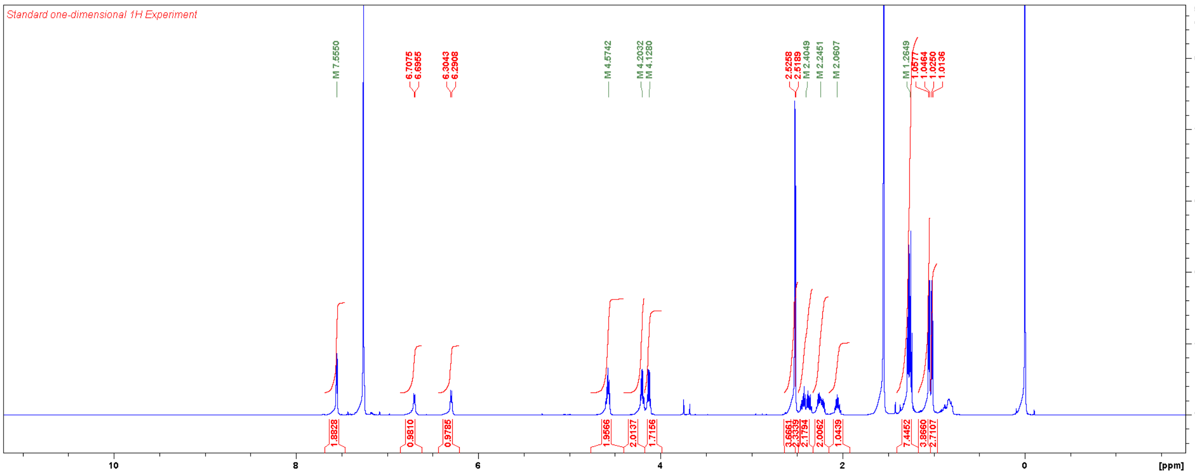


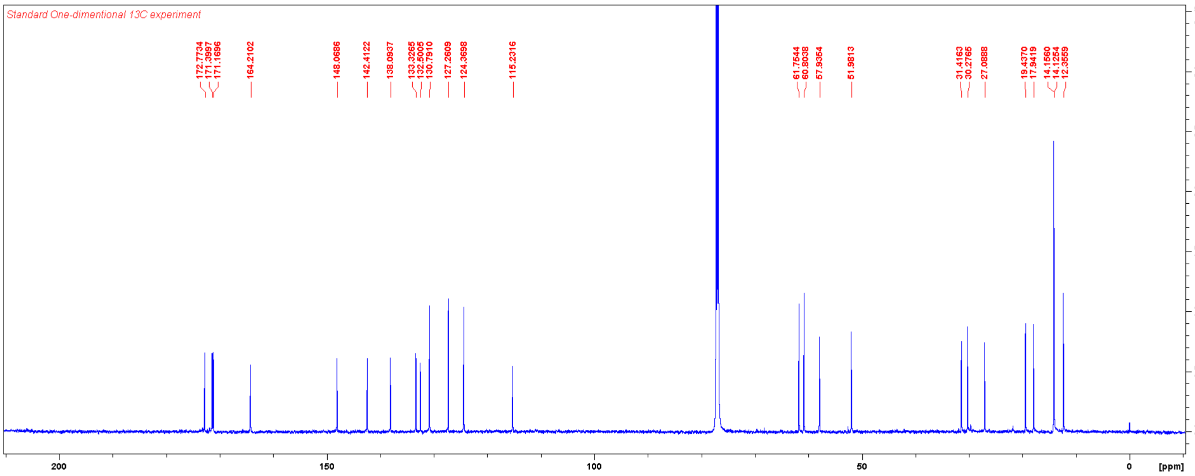

Diethyl (1-(4-methoxyphenyl)-3,5-dimethyl-1*H*-pyrazole-4-carbonyl)-*L*-valyl-*D*-glutamate (**TT023**)

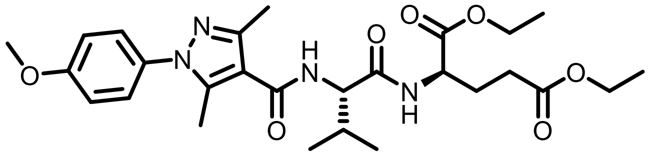

**TT023** was synthesised according to the general procedure for HATU-mediated coupling. Yield: 63 mg (0.12 mmol, 66%).

$^1$H NMR ($CDCl_3$, 600 MHz) δ 7.28 (d, 2H, *J* = 8.8 Hz), 6.97 (d, 2H, *J* = 8.8 Hz), 6.74 (d, 1H, *J* = 7.4 Hz), 6.27 (d, 1H, *J* = 8.1 Hz), 4.57 (m, 2H), 4.20 (m, 2H), 4.12 (m, 2H), 3.85 (s, 3H), 2.53 (s, 3H), 2.46 (s, 3H), 2.44–1.95 (m, 5H), 1.26 (m, 6H), 1.05 (d, 3H, *J* = 6.8 Hz), 1.02 (d, 3H, *J* = 6.8 Hz).

$^{13}$C NMR ($CDCl_3$, 150 MHz) δ 172.8, 171.5, 171.4, 164.8, 159.6, 147.2, 142.5, 131.9, 127.1, 114.4, 114.1, 61.8, 60.8, 58.0, 55.7, 52.0, 31.4, 30.4, 27.2, 19.6, 18.0, 14.3, 14.24, 14.20, 12.3.

HRMS m/z: $[M+H]^+$ Calcd for $C_{27}H_{38}N_4O_7$ 531.2813; Found 531.2811.

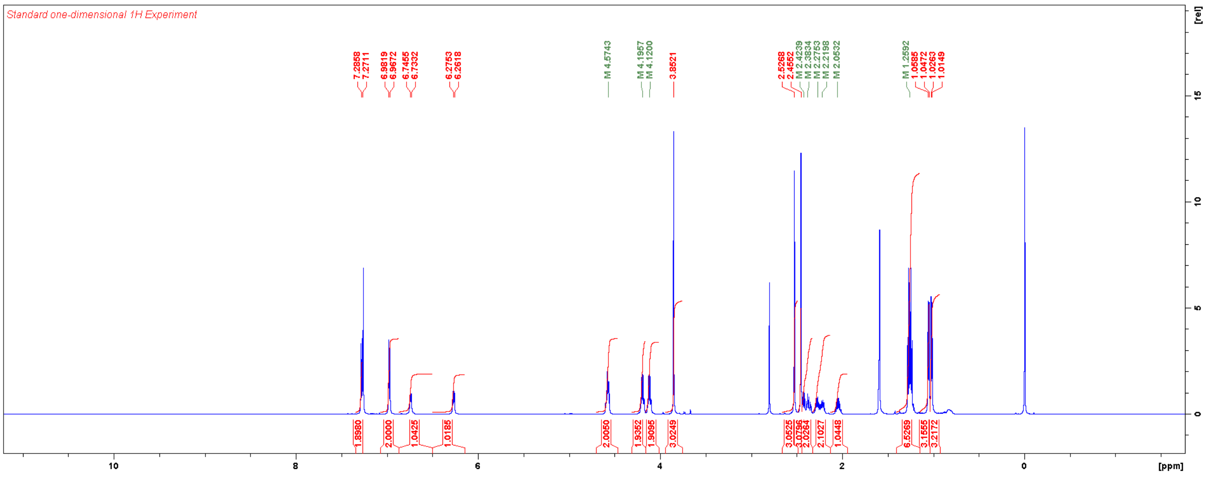


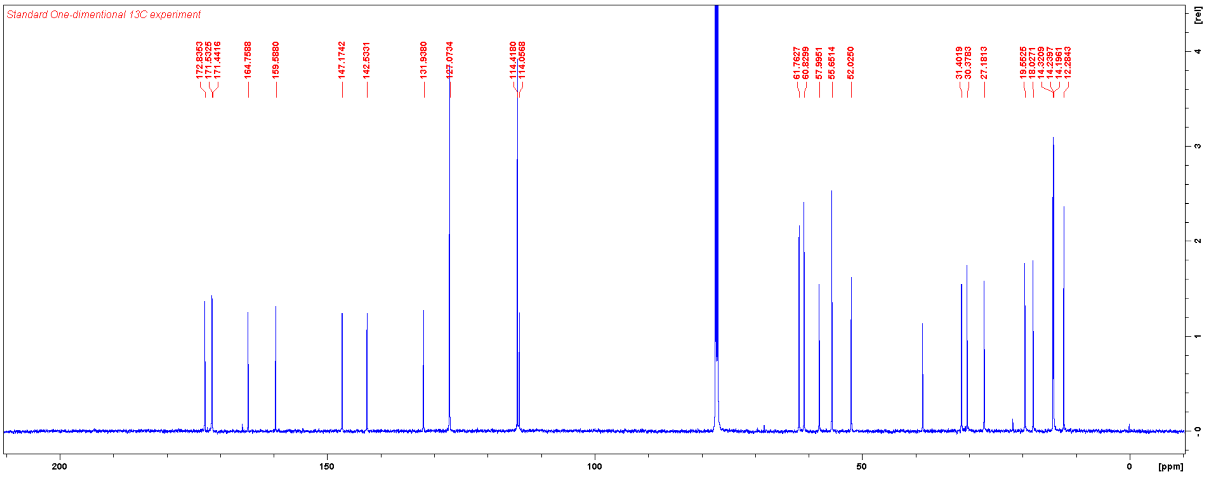

Diethyl (1-(4-chlorophenyl)-3,5-dimethyl-1*H*-pyrazole-4-carbonyl)-*L*-valyl-*D*-glutamate (**TT024**)

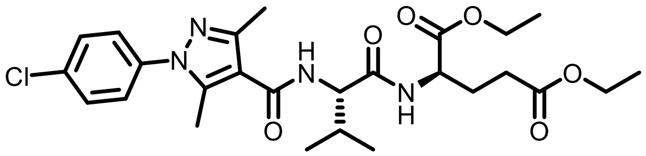

**TT024** was synthesised according to the general procedure for HATU-mediated coupling. Yield: 26 mg (0.049 mmol, 81%).

$^1$H NMR ($CDCl_3$, 600 MHz) δ 7.46 (d, 2H, *J* = 8.6 Hz), 7.34 (d, 2H, *J* = 8.6 Hz), 6.71 (d, 1H, *J* = 7.1 Hz), 6.29 (d, 1H, *J* = 8.0 Hz), 4.57 (m, 2H), 4.20 (m, 2H), 4.13 (m, 2H), 2.53 (s, 3H), 2.50 (s, 3H), 2.40 (m, 2H), 2.24 (m, 2H), 2.05 (m, 1H), 1.26 (m, 6H), 1.05 (d, 3H, *J* = 6.8 Hz), 1.02 (d, 3H, *J* = 6.8 Hz).

$^{13}$C NMR ($CDCl_3$, 150 MHz) δ 172.9, 171.5, 171.3, 164.5, 147.8, 142.5, 137.5, 134.3, 129.5, 126.8, 114.9, 61.9, 60.9, 58.1, 52.1, 31.5, 30.4, 27.2, 19.6, 18.1, 14.33, 14.29, 14.2, 12.4.

HRMS m/z: $[M+H]^+$ Calcd for $C_{26}H_{35}ClN_4O_6$ 535.2318; Found 535.2313.

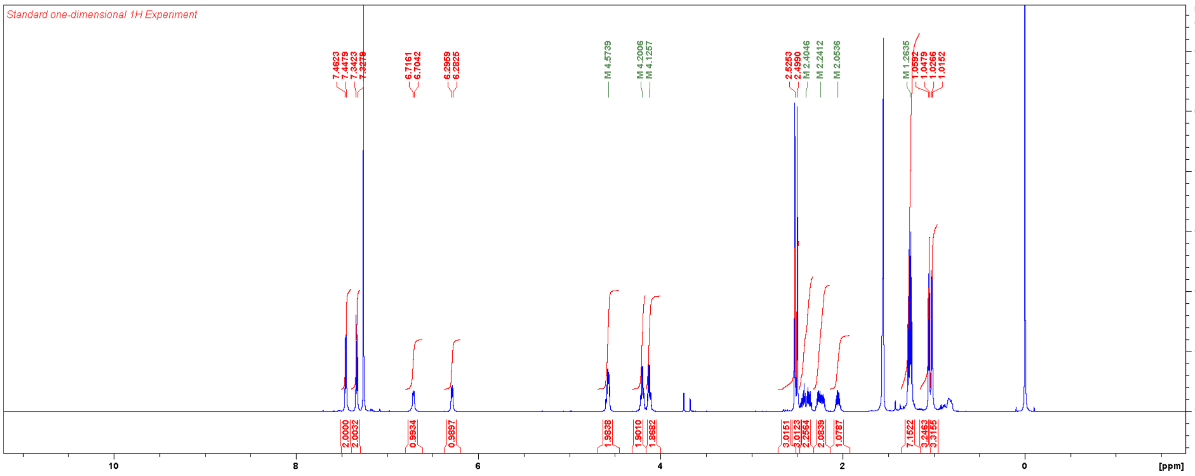


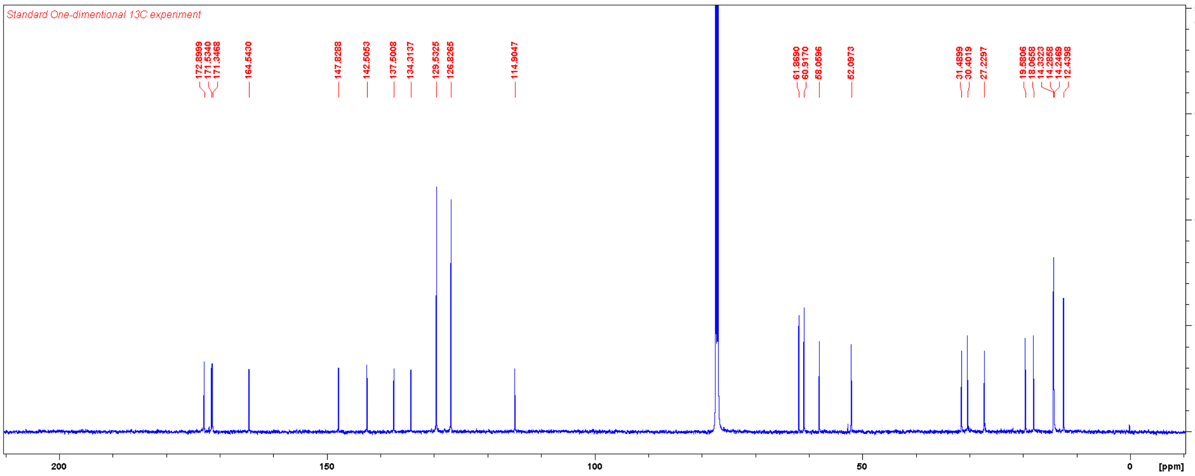

Ethyl 1-(1*H*-indol-6-yl)-3,5-dimethyl-1*H*-pyrazole-4-carboxylate

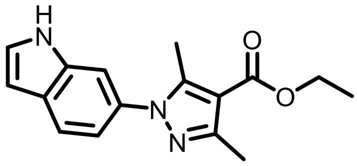

Indole-6-boronic acid (64 mg, 0.40 mmol) and ethyl 3,5-dimethyl-1H-pyrazole-4-carboxylate (67 mg, 0.40 mmol) were weighed in a scintillation vial and dissolved in DMF (2 mL) under ambient atmosphere. To this $Cu(OAc)_2$ (54 mg, 0.30 mmol) and pyridine (64 µL, 0.80 mmol) were added. The reaction mixture was stirred at RT for three days and then diluted with 50% EtOAc in hexanes and washed with brine. Flash chromatography using a gradient of EtOAc in hexanes afforded the title compound. Yield: 62 mg (0.22 mmol, 55%).

$^1$H NMR ($CDCl_3$, 600 MHz) δ 9.17 (br, 1H), 7.67 (br, 1H), 7.39 (br, 1H), 7.29 (br, 1H), 7.04 (br, 1H), 6.57 (br, 1H), 4.32 (q, 2H, *J* = 7.0 Hz), 2.49 (s, 3H), 2.48 (s, 3H), 1.37 (t, 3H, *J* = 7.1 Hz).

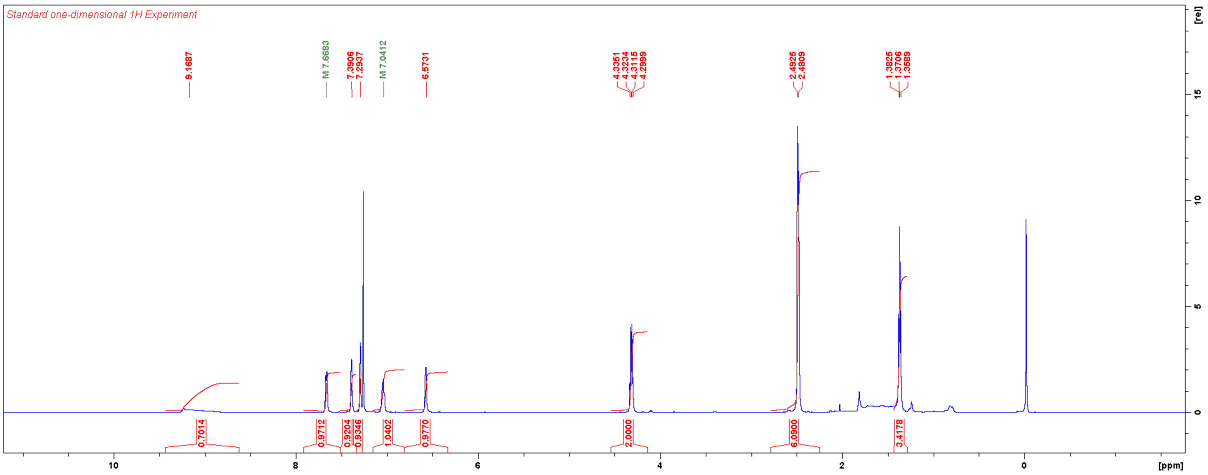

Diethyl (1-(1*H*-indol-6-yl)-3,5-dimethyl-1*H*-pyrazole-4-carbonyl)-*L*-valyl-*D*-glutamate (**TT025**)

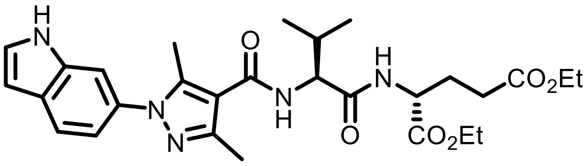

Ethyl 1-(1*H*-indol-6-yl)-3,5-dimethyl-1*H*-pyrazole-4-carboxylate (62 mg, 0.22 mmol) was hydrolysed according to the general procedure for alkaline hydrolysis [Intermediate yield: 57 mg (0.22 mmol, quant.)]. This material was used without further purification to synthesise **TT025** according to the general procedure for HATU-mediated coupling. Yield: 19 mg (0.035 mmol, 59%).

$^{1}$H NMR ($CDCl_3$, 600 MHz) δ 9.09 (br, 1H), 7.68 (m, 1H), 7.41 (br, 1H), 7.33 (br, 1H), 7.06 (dd, 1H, *J* = 8.3, 1.4 Hz), 6.84 (d, 1H, *J* = 7.3 Hz), 6.63 (br, 1H), 6.58 (s, 1H), 4.58 (m, 2H), 4.20 (q, 2H, *J* = 7.1 Hz), 4.13 (q, 2H, *J* = 6.8 Hz), 2.46 (s, 3H), 2.45 (s, 3H), 2.40 (m, 2H), 2.27 (m, 2H), 2.07 (m, 1H), 1.26 (m, 6H), 1.07 (d, 3H, *J* = 6.7 Hz), 1.05 (d, 3H, *J* = 6.8 Hz).

$^{13}$C NMR ($CDCl_3$, 150 MHz) δ 172.8, 171.4, 171.3, 164.5, 146.9, 142.9, 135.4, 132.3, 128.2, 126.5, 121.0, 117.7, 114.2, 109.0, 102.6, 61.7, 60.8, 58.2, 52.0, 31.2, 30.3, 27.1, 19.5, 18.0, 14.2, 14.1, 13.8, 12.3.

HRMS m/z: $[M+H]^+$ Calcd for $C_{28}H_{37}N_5O_6$ 540.2817; Found 540.2812.

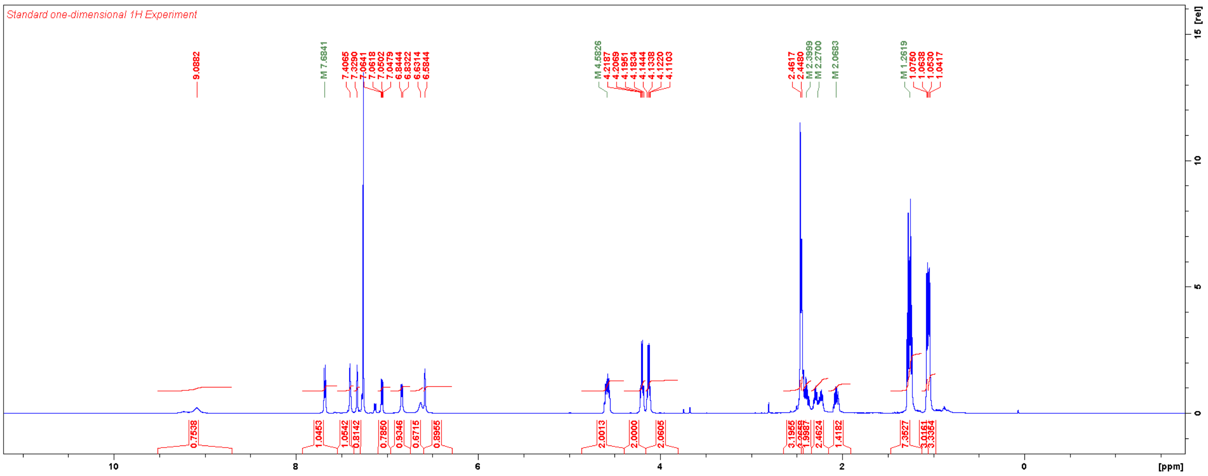


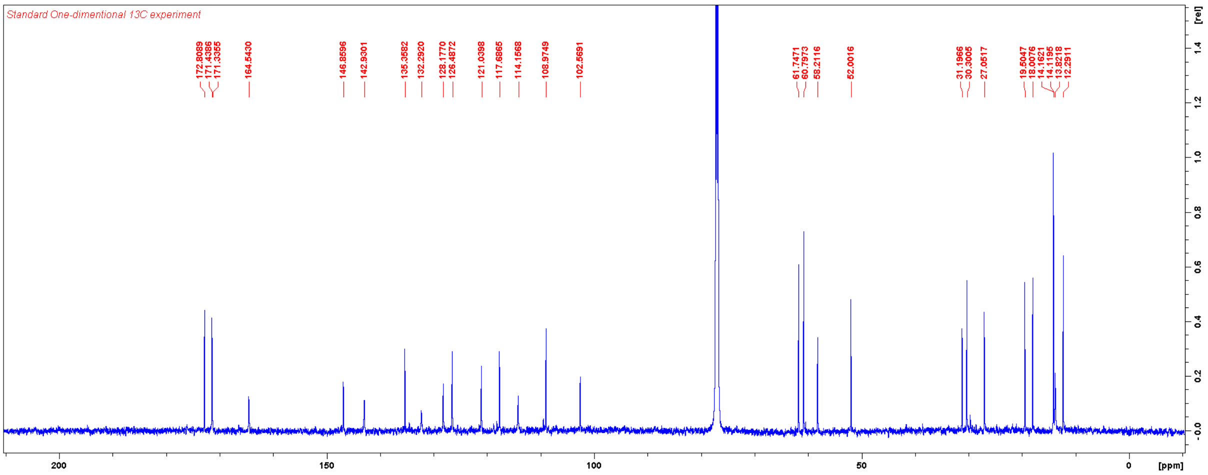

Diethyl (1-(6-aminopyridin-3-yl)-3,5-dimethyl-1*H*-pyrazole-4-carbonyl)-*L*-valyl-*D*-glutamate (**TT026**)

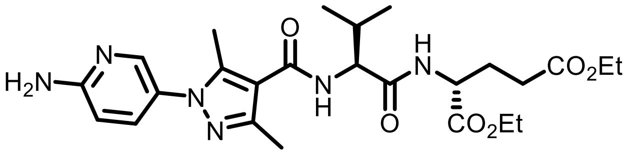

**TT027** (7 mg, 0.011 mmol) was stirred in 4 N HCl in 1,4-dioxane at RT for 4 h. Upon completion of the reaction, the reaction mixture was concentrated to dryness to afford **TT026** as HCl salt. Yield: 6 mg (0.011 mmol, quant.).

HRMS m/z: $[M+H]^+$ Calcd for $C_{25}H_{36}N_6O_6$ 517.2769; Found 517.2768.

*tert*-Butyl (5-(4,4,5,5-tetramethyl-1,3,2-dioxaborolan-2-yl)pyridin-2-yl)carbamate

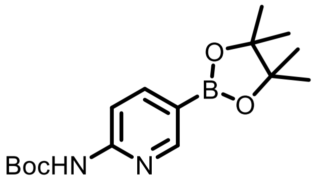


To a solution of 5-(4,4,5,5-tetramethyl-1,3,2-dioxaborolan -2-yl)pyridin-2-amine (220 mg, 1.00 mmol), triethylamine (348 µL, 2.50 mmol), and DMAP (12 mg, 0.10 mmol) in DCM (5 mL) was added di-*tert*-butyl decarbonate (276 µL, 1.20 mmol) at 0 °C. The reaction mixture was allowed to warm to RT overnight. The reaction mixture was then diluted with EtOAc and washed with brine, dried over $Na_2SO_4$, filtered, and concentrated. The residue was purified with flash chromatography using a gradient of EtOAc in hexanes to afford the title compound as white solid. Yield: 147 mg (0.44 mmol, 44%).

$^1$H NMR ($CDCl_3$, 600 MHz) δ 9.55 (br, 1H), 8.70 (s, 1H), 8.03 (m, 2H), 1.57 (s, 9H), 1.33 (s, 12H).

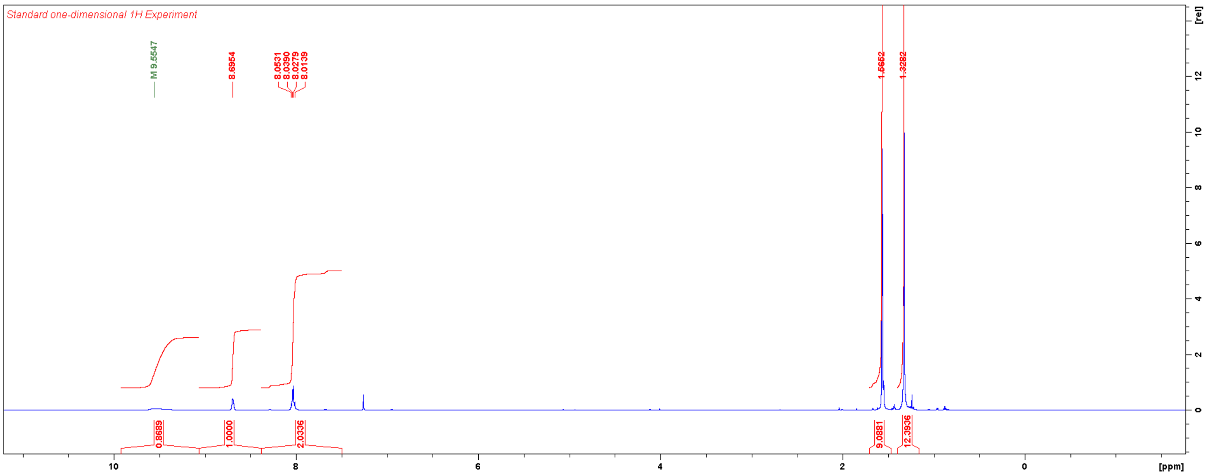

Ethyl 1-(6-((tert-butoxycarbonyl)amino)pyridin-3-yl)-3,5-dimethyl-1*H*-pyrazole-4-carboxylate

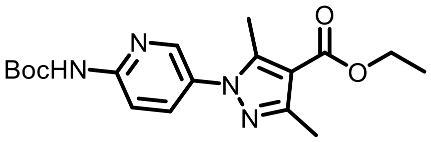


*tert*-Butyl (5-(4,4,5,5-tetramethyl-1,3,2-dioxaborolan-2-yl)pyridin-2-yl)carbamate (67 mg, 0.20 mmol) and ethyl 3,5-dimethyl-1H-pyrazole-4-carboxylate (34 mg, 0.20 mmol) were weighed in a scintillation vial and suspended in DMF (2 mL) under ambient atmosphere. To this $Cu(OAc)_2$ (27 mg, 0.15 mmol) and pyridine (32 µL, 0.40 mmol) were added. The reaction mixture was stirred at 50 °C for 24 h and then diluted with 50% EtOAc in hexanes and washed with brine. Flash chromatography using a gradient of EtOAc in hexanes afforded the title compound. Yield: 37 mg (0.10 mmol, 51%).

$^1$H NMR ($CDCl_3$, 600 MHz) δ 8.33 (s, 1H), 8.11 (d, 1H, *J* = 8.9 Hz), 7.97 (br, 1H), 7.70 (dd, 1H, *J* = 8.8, 2.3 Hz), 4.33 (q, 2H, *J* = 7.1 Hz), 2.50 (s, 3H), 2.48 (s, 3H), 1.54 (s, 9H), 1.38 (t, 3H, *J* = 7.1 Hz).

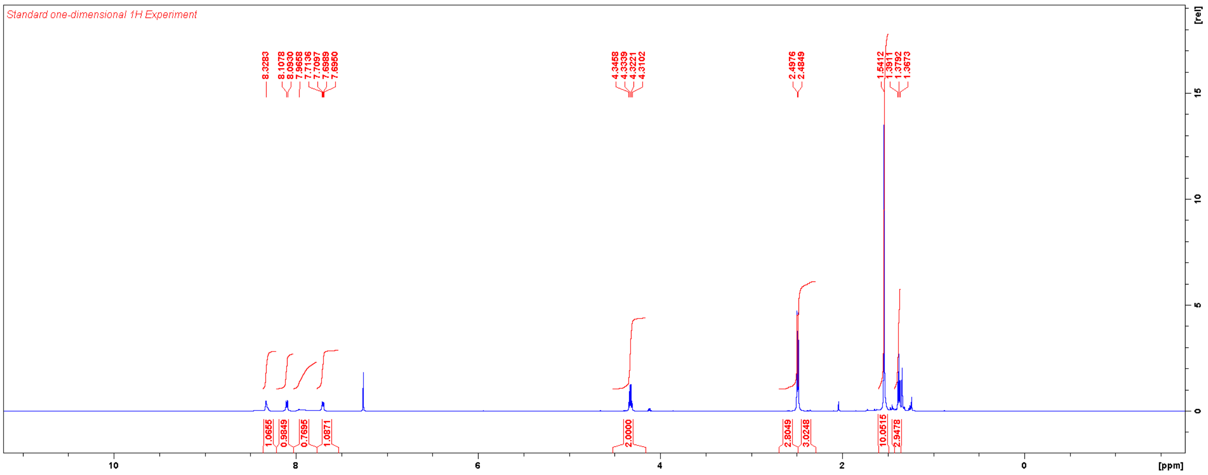

Diethyl (1-(6-((*tert*-butoxycarbonyl)amino)pyridin-3-yl)-3,5-dimethyl-1*H*-pyrazole-4-carbonyl)-*L*-valyl-*D*-glutamate (**TT027**)

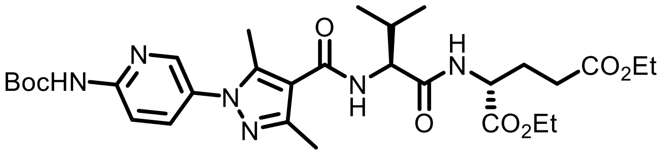


Ethyl 1-(6-((tert-butoxycarbonyl)amino)pyridin-3-yl)-3,5-dimethyl-1*H*-pyrazole-4-carboxylate (37 mg, 0.10 mmol) was hydrolysed according to the general procedure for alkaline hydrolysis [Intermediate yield: 33 mg (0.10 mmol, quant.)]. This material was used without further purification to synthesise **TT027** according to the general procedure for HATU-mediated coupling. Yield: 16 mg (0.026 mmol, 26%).

$^1$H NMR ($CDCl_3$, 600 MHz) δ 8.29 (s, 1H), 8.20 (br, 1H), 7.80 (br, 1H), 6.75 (d, 1H, *J* = 7.3 Hz), 6.31 (d, 1H, *J* = 8.1 Hz), 4.58 (m, 2H), 4.20 (m, 2H), 4.13 (q, 2H, *J* = 6.8 Hz), 2.52 (s, 3H), 2.50 (s, 3H), 2.41 (m, 2H), 2.25 (m, 2H), 2.05 (m, 1H), 1.55 (s, 9H), 1.26 (m, 6H), 1.05 (d, 3H, *J* = 6.8 Hz), 1.02 (d, 3H, *J* = 6.8 Hz).

$^{13}$C NMR ($CDCl_3$, 150 MHz) δ 172.8, 171.4, 171.2, 164.2, 151.9, 151.3, 148.2, 142.8, 136.3, 130.7, 115.0, 112.7, 82.1, 61.8, 60.8, 58.0, 52.0, 31.4, 30.3, 28.2, 27.1, 19.5, 17.9, 14.2, 14.1, 12.1.

HRMS m/z: $[M+H]^+$ Calcd for $C_{30}H_{44}N_6O_8$ 617.3293; Found 617.3290.

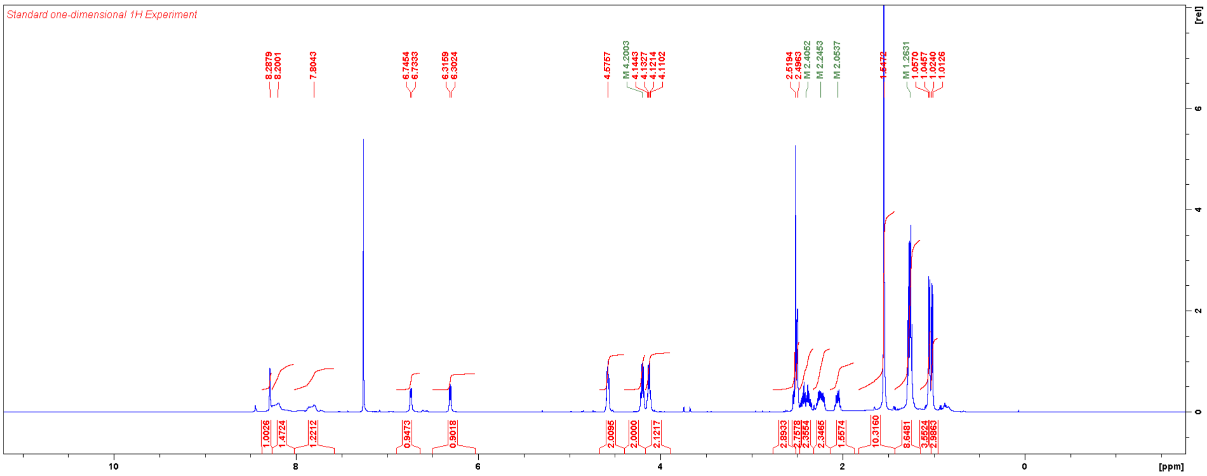


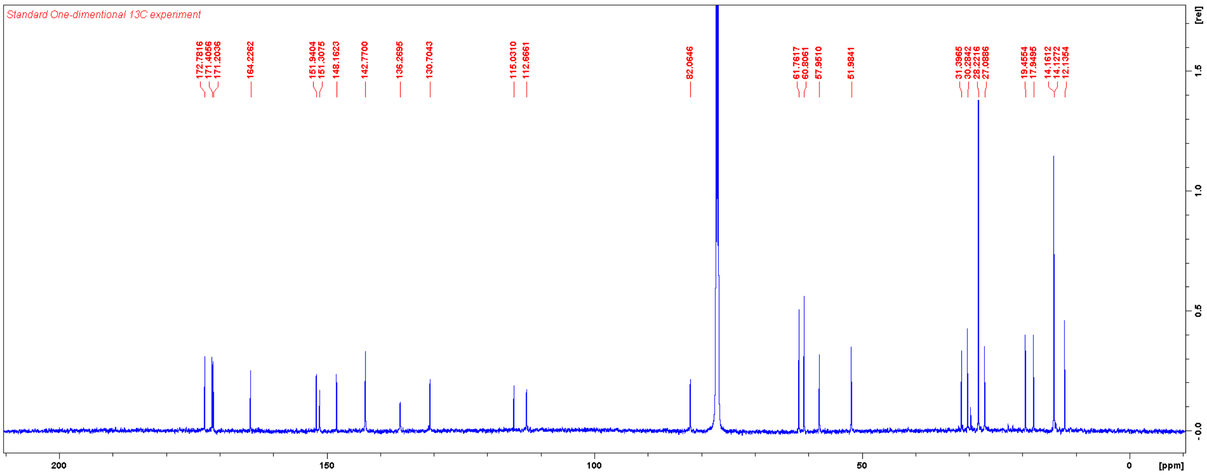

Ethyl 1-(1*H*-indol-5-yl)-3,5-dimethyl-1*H*-pyrazole-4-carboxylate

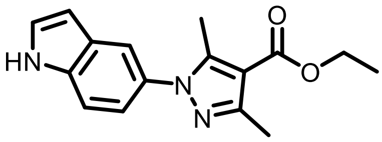

(1*H*-Indol-5-yl)boronic acid (64 mg, 0.40 mmol) and ethyl 3,5-dimethyl-1H-pyrazole-4-carboxylate (67 mg, 0.40 mmol) were weighed in a scintillation vial and dissolved in DMF (2 mL) under ambient atmosphere. To this $Cu(OAc)_2$ (54 mg, 0.30 mmol) and pyridine (64 µL, 0.80 mmol) were added. The reaction mixture was stirred at RT for three days and then diluted with 50% EtOAc in hexanes and washed with brine. Flash chromatography using a gradient of EtOAc in hexanes afforded the title compound. Yield: 76 mg (0.27 mmol, 67%).

$^1$H NMR ($CDCl_3$, 600 MHz) δ 8.39 (br, 1H), 7.61 (s, 1H), 7.44 (d, *J* = 8.5 Hz), 7.30 (br, 1H), 7.17 (dd, *J* = 8.5, 1.7 Hz), 6.61 (br, 1H), 4.34 (q, 2H, *J* = 7.1 Hz), 2.52 (s, 3H), 2.49 (s, 3H), 1.39 (t, 3H, *J* = 7.1 Hz).

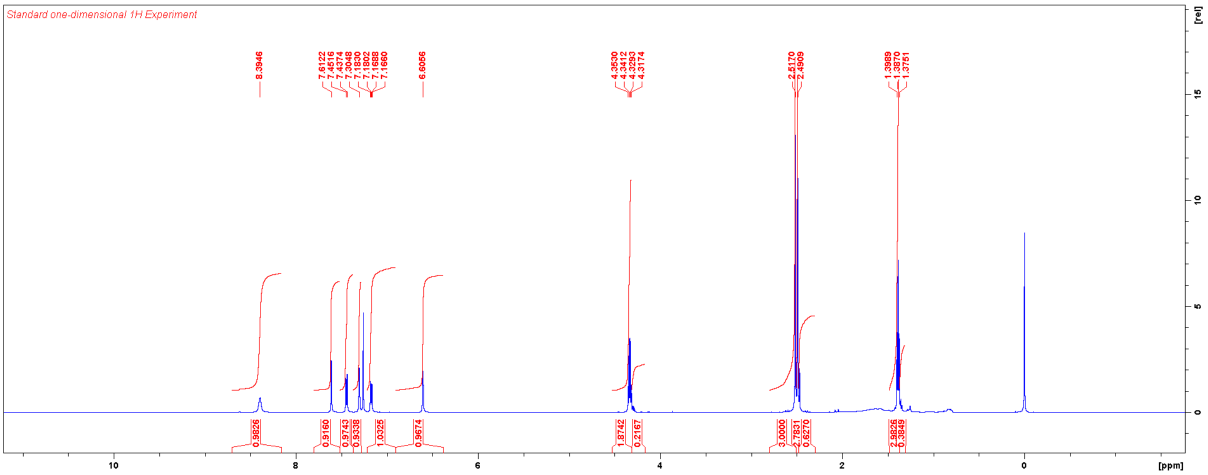

Diethyl (1-(1*H*-indol-5-yl)-3,5-dimethyl-1*H*-pyrazole-4-carbonyl)-*L*-valyl-*D*-glutamate (**TT028**)

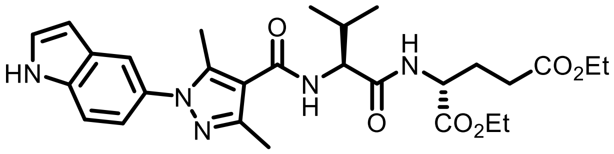


Ethyl 1-(1*H*-indol-5-yl)-3,5-dimethyl-1*H*-pyrazole-4-carboxylate (76 mg, 0.27 mmol) was hydrolysed according to the general procedure for alkaline hydrolysis [Intermediate yield: 70 mg (0.27 mmol, quant.)]. This material was used without further purification to synthesise **TT028** according to the general procedure for HATU-mediated coupling. Yield: 23 mg (0.043 mmol, 71%).

$^1$H NMR ($CDCl_3$, 600 MHz) δ 7.62 (s, 1H), 7.41 (br, 1H), 7.32 (br, 1H), 7.12 (br, 1H), 6.86 (br, 1H), 6.59 (br, 1H), 6.53 (br, 1H), 4.60 (m, 2H), 4.24 (q, 2H, $J$ = 6.7 Hz), 4.16 (q, 2H, $J$ = 6.9 Hz), 2.64 (s, 3H), 2.51 (s, 3H), 2.45 (m, 2H), 2.29 (m, 2H), 2.10 (m, 1H), 1.30 (m, 6H), 1.10 (d, 3H, $J$ = 6.7 Hz), 1.07 (d, 3H, $J$ = 6.8 Hz).

$^{13}$C NMR ($CDCl_3$, 150 MHz) δ 172.8, 171.44, 171.41, 164.7, 146.7, 143.0, 135.5, 130.9, 127.9, 126.2, 119.9, 118.3, 113.7, 111.6, 103.1, 61.7, 60.8, 58.1, 52.0, 31.2, 30.3, 27.1, 19.5, 18.0, 14.2, 14.1, 12.3.

HRMS m/z: $[M+H]^+$ Calcd for $C_{28}H_{37}N_5O_6$ 540.2817; Found 540.2813.

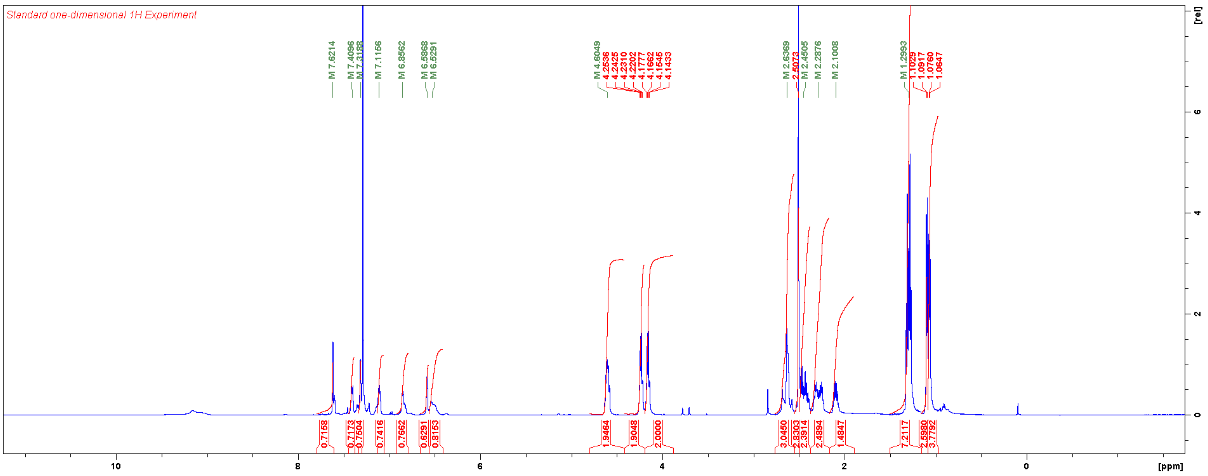


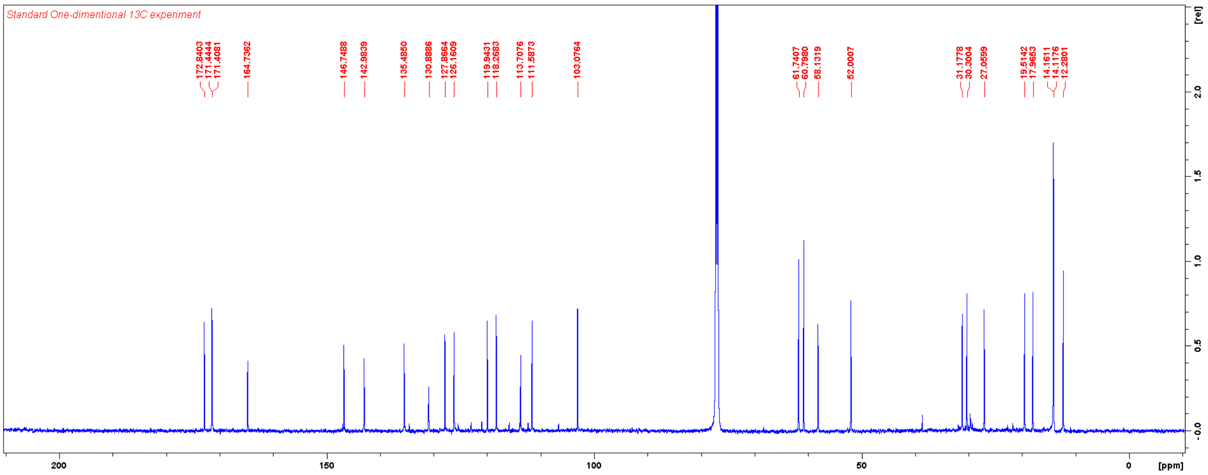

Ethyl 3,5-dimethyl-1-(1*H*-pyrrolo[2,3-b]pyridin-5-yl)-1*H*-pyrazole-4-carboxylate

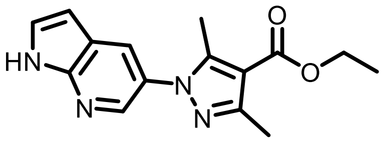

(1H-Pyrrolo[2,3-b]pyridin-5-yl)boronic acid (65 mg, 0.40 mmol) and ethyl 3,5-dimethyl-1H-pyrazole-4-carboxylate (67 mg, 0.40 mmol) were weighed in a scintillation vial and dissolved in DMF (2 mL) under ambient atmosphere. To this $Cu(OAc)_2$ (54 mg, 0.30 mmol) and pyridine (64 µL, 0.80 mmol) were added. The reaction mixture was stirred at RT for three days and then diluted with 50% EtOAc in hexanes and washed with brine. Flash chromatography using a gradient of EtOAc in hexanes afforded the title compound. Yield: 65 mg (0.23 mmol, 57%).

$^1$H NMR ($CDCl_3$, 600 MHz) δ 9.23 (br, 1H), 8.35 (s, 1H), 7.96 (s, 1H), 7.44 (d, 1H, *J* = 2.3 Hz), 6.59 (dd, 1H, *J* = 3.2 , 2.0 Hz), 4.35 (q, 2H, *J* = 7.1 Hz), 2.52 (s, 3H), 2.51 (s, 3H), 1.39 (t, 3H, *J* = 7.1 Hz).

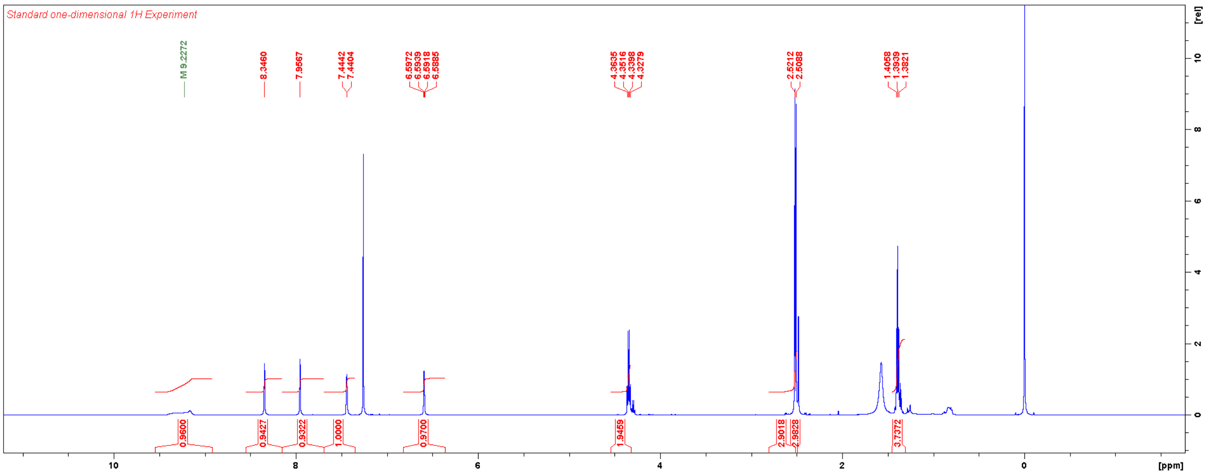

Diethyl (3,5-dimethyl-1-(1*H*-pyrrolo[2,3-b]pyridin-5-yl)-1*H*-pyrazole-4-carbonyl)-*L*-valyl-*D*-glutamate (**TT029**)

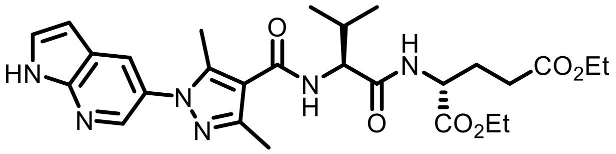


Ethyl 3,5-dimethyl-1-(1*H*-pyrrolo[2,3-b]pyridin-5-yl)-1*H*-pyrazole-4-carboxylate (65 mg, 0.23 mmol) was hydrolysed according to the general procedure for alkaline hydrolysis [Intermediate yield: 26 mg (0.10 mmol, 44%)]. This material was used without further purification to synthesise **TT029** according to the general procedure for HATU-mediated coupling. Yield: 16 mg (0.030 mmol, 30%).

$^1$H NMR ($CDCl_3$, 600 MHz) δ 8.49 (s, 1H), 8.40 (s, 1H), 7.69 (s, 1H), 7.07 (d, 1H, *J* = 7.4 Hz), 6.81 (d, 1H, *J* = 2.2 Hz), 6.59 (d, 1H, *J* = 8.3 Hz), 4.67 (m, 1H), 4,62 (m, 1H), 4.22 (q, 2H, *J* = 6.8 Hz), 4.16 (q, 2H, *J* = 7.1 Hz), 2.55 (s, 3H), 2.53 (s, 3H), 2.46 (m, 2H), 2.29 (m, 2H), 2.11 (m, 1H), 1.29 (m, 6H), 1.09 (d, 3H, *J* = 6.7 Hz), 1.07 (d, *J* = 6.8 Hz).

$^{13}$C NMR ($CDCl_3$, 150 MHz) δ 172.7, 171.8, 171.4, 164.9, 147.9, 146.6, 142.5, 138.9, 129.1, 127.9, 127.0, 120.6, 115.1, 101.6, 61.7, 60.8, 58.4, 52.1, 31.3, 30.3, 27.1, 19.5, 18.2, 14.2, 14.1, 13.9, 12.1.

HRMS m/z: $[M+H]^+$ Calcd for $C_{27}H_{36}N_6O_6$ 541.2769; Found 541.2764.

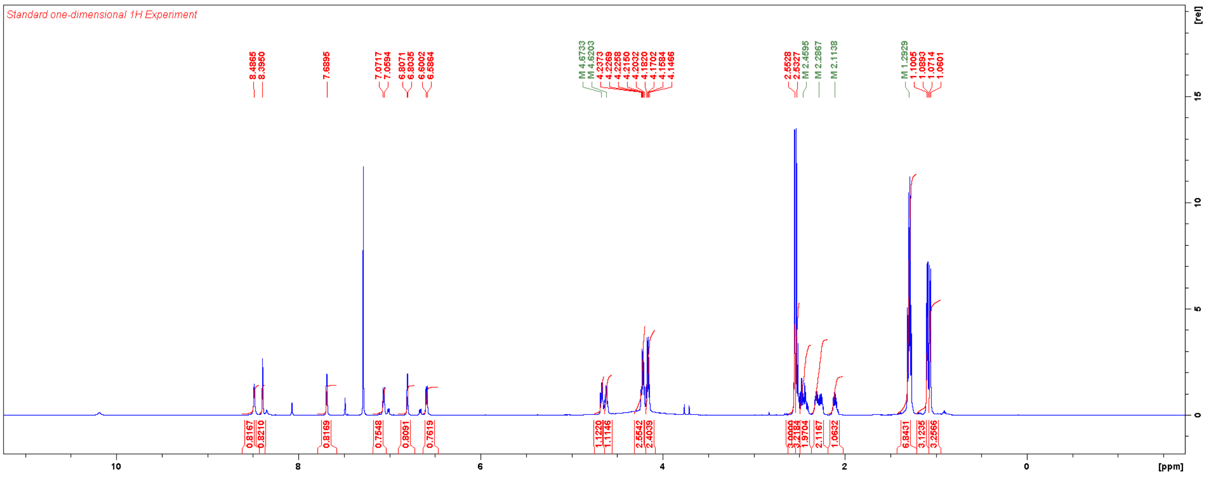


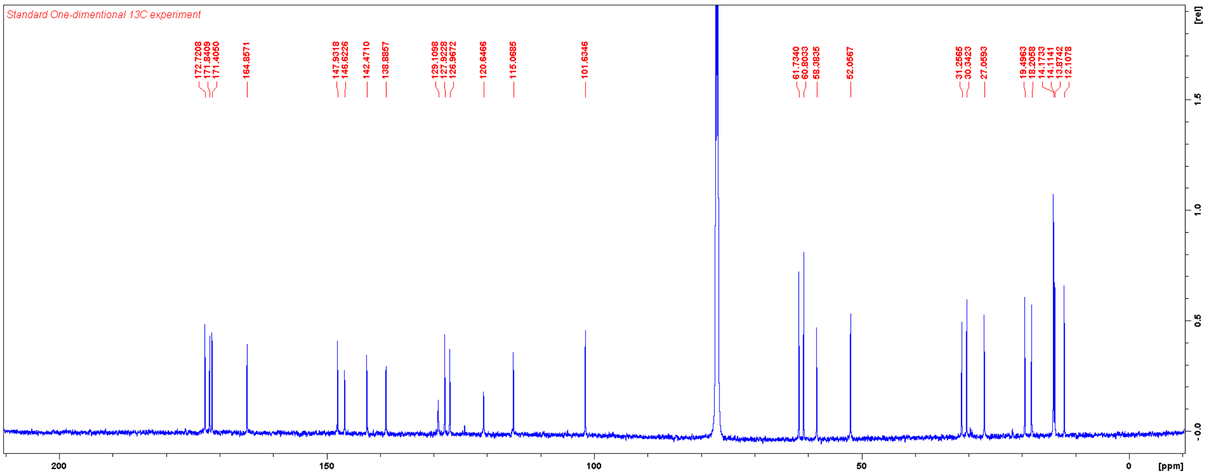

Ethyl 3,5-dimethyl-1-(2-oxo-1,2-dihydropyridin-4-yl)-1*H*-pyrazole-4-carboxylate

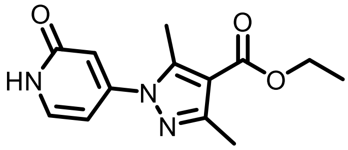

A solution of 4-hydrazinylpyridin-2(1*H*)-one (197 mg, 1.57 mmol), ethyl 2-acetyl-3-oxobutanoate (246 μL, 1.57 mmol) in acetic acid (3 mL) was stirred at RT overnight. The solvent was evaporated, and the residue was purified with column chromatography eluting with acetonitrile (two column volume) followed by 5% MeOH/DCM. Yield: 381 mg (1.45 mmol, 92%).

$^1$H NMR ($CDCl_3$, 600 MHz) δ 7.45 (d, 1H, *J* = 7.1 Hz), 6.67 (d, 1H, *J* = 7.1, 2.0 Hz), 6.57 (d, 1H, *J* = 1.8 Hz), 4.34 (q, 2H, *J* = 7.1 Hz), 2.69 (s, 3H), 2.48 (s, 3H), 1.38 (t, 3H, *J* = 7.1 Hz).

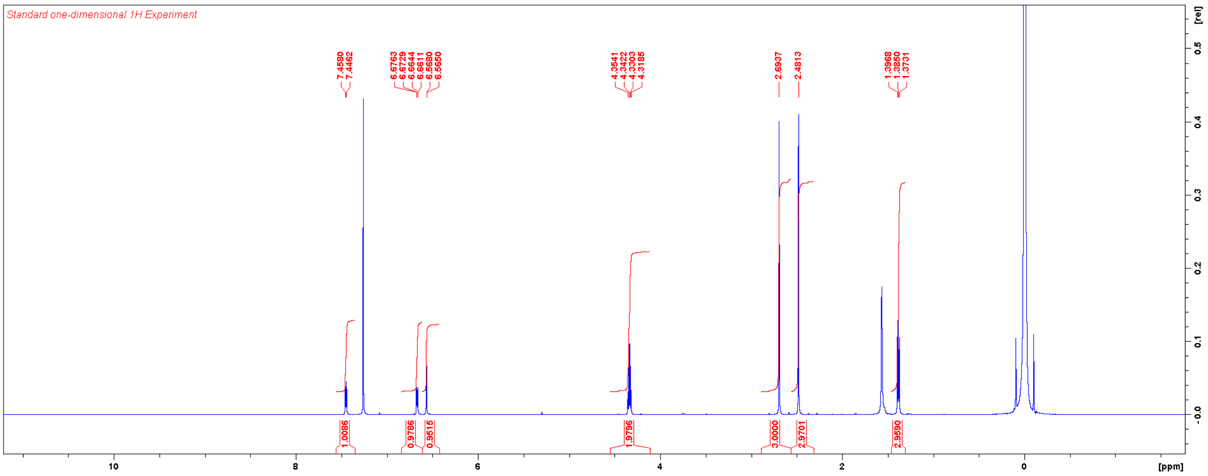

Diethyl (3,5-dimethyl-1-(2-oxo-1,2-dihydropyridin-4-yl)-1*H*-pyrazole-4-carbonyl)-*L*-valyl-*D*-glutamate (**TT030**)

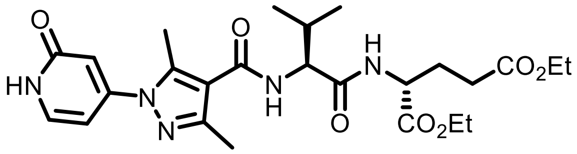

Ethyl 3,5-dimethyl-1-(2-oxo-1,2-dihydropyridin-4-yl)-1*H*-pyrazole-4-carboxylate (133 mg, 0.51 mmol) was hydrolysed according to the general procedure for alkaline hydrolysis. Upon completion of the reaction, the solvent was evaporated, and the residual aqueous phase was acidified with 1N HCl, which resulted in precipitation. The precipitates were collected by filtration and dried under vacuum at 60 °C. [Intermediate yield: 119 mg (0.51 mmol, quant.)]. This material was used without further purification to synthesise **TT030** according to the general procedure for HATU-mediated coupling. Upon completion of the reaction, reaction mixture was diluted with EtOAc, washed with aqueous $NH_4Cl$, dried over $Na_2SO_4$, filtered, and concentrated. The residue was purified with column chromatography using two different columns and eluents: first with 5% MeOH/DCM; then on a new column with 100% THF. Yield: 19 mg (0.037 mmol, 37%).

$^1$H NMR ($CDCl_3$, 600 MHz) δ 7.40 (d, 1H, *J* = 7.1 Hz), 6.91 (d, 1H, *J* = 7.1 Hz), 6.62 (m, 2H), 6.51 (d, 1H, *J* = 7.1 Hz), 4.57 (m, 2H), 4.20 (m, 2H), 4.13 (q, 2H, *J* = 7.1 Hz), 2.62 (s, 3H), 2.48 (s, 3H), 2.42 (m, 2H), 2.25 (m, 2H), 2.06 (m, 1H), 1.27 (m, 6H), 1.06 (d, 3H, *J* = 6.8 Hz), 1.04 (d, 3H, *J* = 6.8 Hz).

$^{13}$C NMR ($CDCl_3$, 150 MHz) δ 172.8, 171.8, 171.6, 164.9, 164.3, 150.1, 149.2, 142.2, 135.4, 117.4, 111.6, 103.9, 61.8, 60.8, 58.4, 52.0, 31.2, 30.4, 27.0, 19.5, 18.2, 14.2, 14.1, 13.9, 13.1.

HRMS m/z: $[M+H]^+$ Calcd for $C_{25}H_{35}N_5O_7$ 518.2609; Found 518.2606.

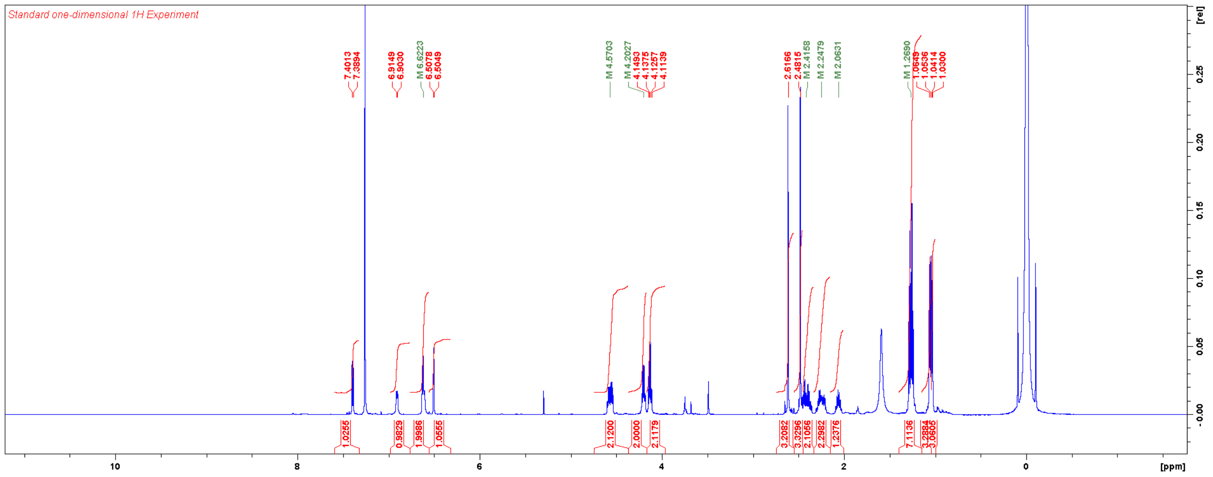

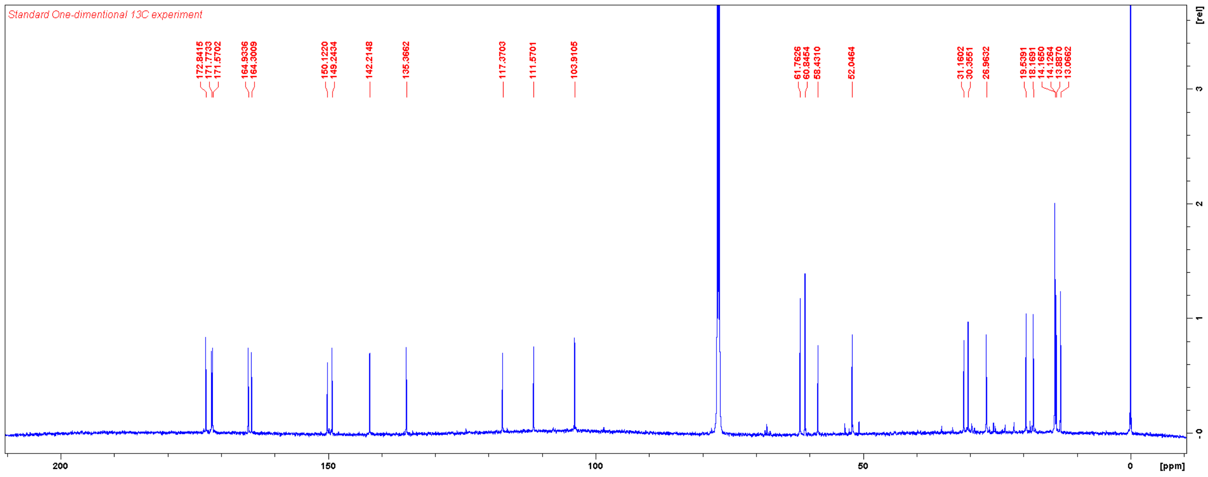
Standard One-dimentional 13C experiment
172.8415
171.7733
171.5702
164.9336
164.3009
150.1220
149.2434
142.2148
135.3662
117.3703
111.5701
103.9105
61.7626
60.8454
58.4310
52.0464
31.1602
30.3551
26.9632
19.5391
18.1691
14.1650
14.1264
13.8870
13.0662
200
150
100
50
0
[ppm]
[rel]
3
2
1
0

Ethyl 3,5-dimethyl-1-(pyridin-4-yl)-1*H*-pyrazole-4-carboxylate

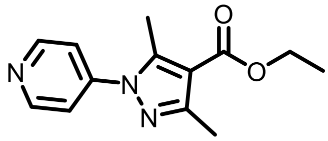

A solution of 4-hydrazinylpyridine hydrochloride (73 mg, 0.5 mmol), ethyl 2-acetyl-3-oxobutanoate (78 μL, 0.5 mmol) in acetic acid (1 mL) was stirred at RT for three days. The solvent was evaporated, and the residue was purified with flash chromatography using a gradient of EtOAc in hexanes. Yield: 78 mg (0.28 mmol, 57%).

$^{1}$H NMR ($CDCl_3$, 600 MHz) δ 8.73 (d, 2H, *J* = 5.9 Hz), 7.44 (d, 2H, *J* = 6.1 Hz), 4.34 (q, 2H, *J* = 7.1 Hz), 2.66 (s, 3H), 2.50 (s, 3H), 1.39 (t, 3H, *J* = 7.1 Hz).

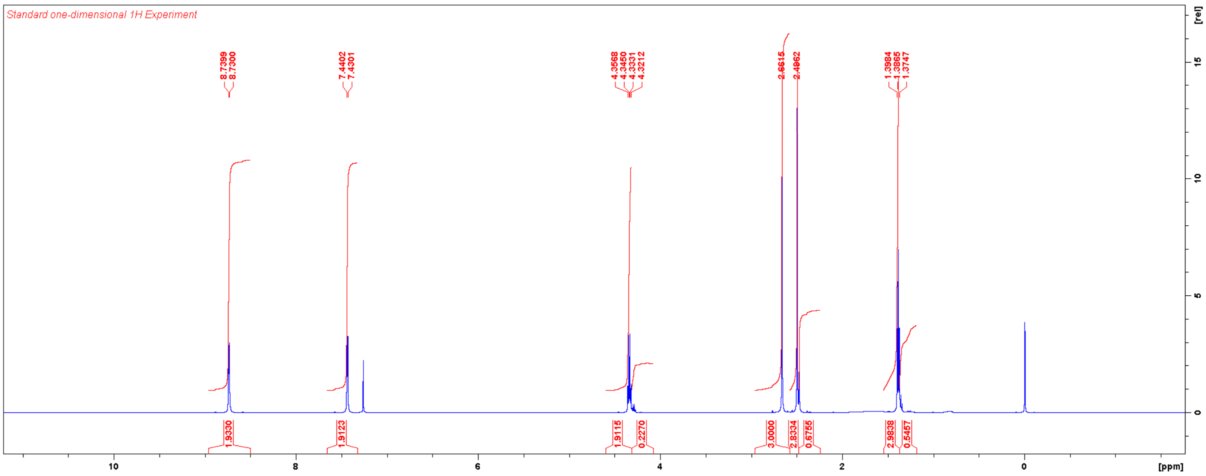

Diethyl (3,5-dimethyl-1-(pyridin-4-yl)-1*H*-pyrazole-4-carbonyl)-*L*-valyl-*D*-glutamate (**TT031**)

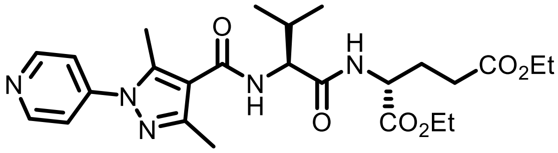

Ethyl 3,5-dimethyl-1-(pyridin-4-yl)-1*H*-pyrazole-4-carboxylate (78 mg, 0.28 mmol) was hydrolysed according to the general procedure for alkaline hydrolysis. Upon completion of the reaction, the aqueous phase was neutralised (pH ≈ 6) with 1 N HCl resulting in precipitation. The precipitates were collected by filtration to afford the carboxylic acid derivative [Intermediate yield: 54 mg (0.25 mmol, 89%)]. This material was used without further purification to synthesise **TT031** according to the general procedure for HATU-mediated coupling. Yield: 17 mg (0.034 mmol, 56%).

$^1$H NMR (CDCl$_3$, 600 MHz) δ 8.78 (d, 2H, *J* = 5.6 Hz), 8.14 (d, 2H, *J* = 5.8 Hz), 6.88 (br, 1H), 6.51 (d, 1H, *J* = 5.3 Hz), 4.57 (m, 2H), 4.21 (q, 2H, *J* = 7.0 Hz), 4.14 (q, 2H, *J* = 7.0 Hz), 2.83 (s, 3H), 2.52 (s, 3H), 2.42 (m, 2H), 2.23 (m, 2H), 2.08 (m, 1H), 1.28 (m, 6H), 1.05 (d, 3H, *J* = 6.8 Hz), 1.02 (d, 3H, *J* = 6.8 Hz).

$^{13}$C NMR (CDCl$_3$, 150 MHz) δ 172.8, 171.4, 171.1, 163.9, 149.7, 149.3, 146.7, 142.6, 118.3, 117.0, 61.8, 60.8, 58.0, 52.0, 31.5, 30.3, 27.0, 19.4, 18.0, 14.2, 14.1, 12.9.

HRMS m/z: [M+H]$^+$ Calcd for $C_{25}H_{35}N_5O_6$ 502.2660; Found 502.2657.

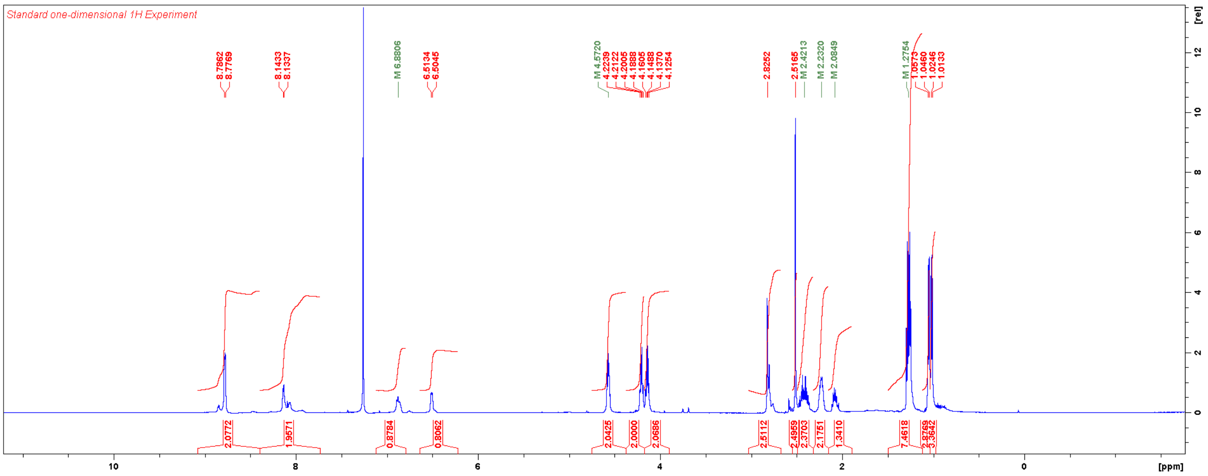


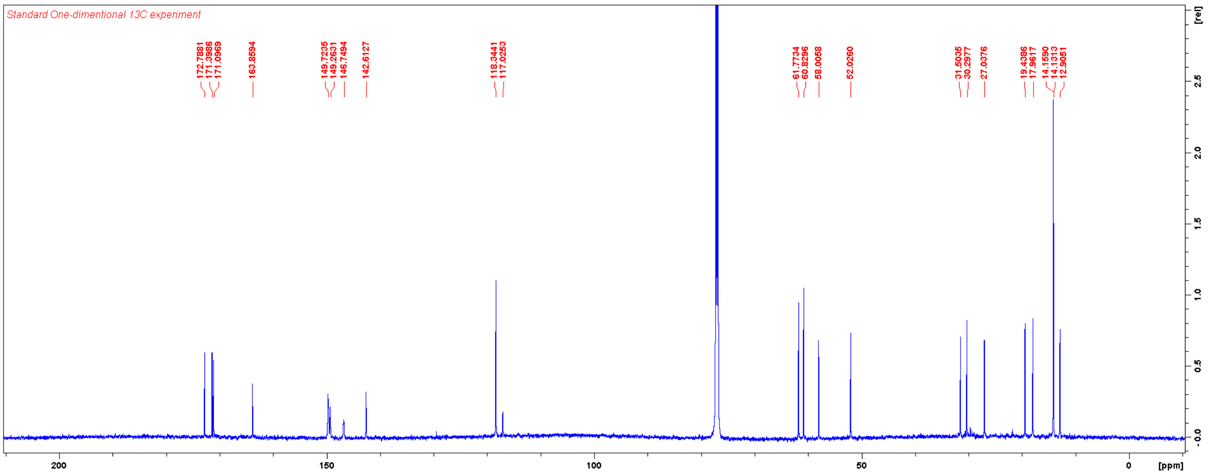

Ethyl 3,5-dimethyl-1-(pyridin-2-yl)-1*H*-pyrazole-4-carboxylate

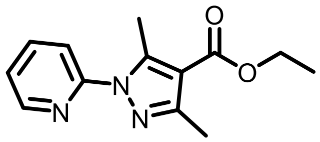

A solution of 2-hydrazinylpyridine (55 mg, 0.5 mmol) and ethyl 2-acetyl-3-oxobutanoate (78 μL, 0.5 mmol) in acetic acid (0.5 mL) was stirred at RT overnight. The solvent was evaporated, and the residue was purified with flash chromatography using a gradient of EtOAc in hexanes. Yield: 109 mg (0.40 mmol, 79%).

$^1$H NMR ($CDCl_3$, 600 MHz) δ 8.49 (d, 1H, *J* = 4.1 Hz), 7.83 (dd, 1H, *J* = 7.1, 7.1 Hz), 7.76 (d, 1H, *J* = 8.1 Hz), 7.26 (dd, 1H, *J* = 6.1 Hz), 4.33 (q, 2H, *J* = 7.1 Hz), 2.85 (s, 3H), 2.50 (s, 3H), 1.38 (t, 3H, *J* = 7.1 Hz).

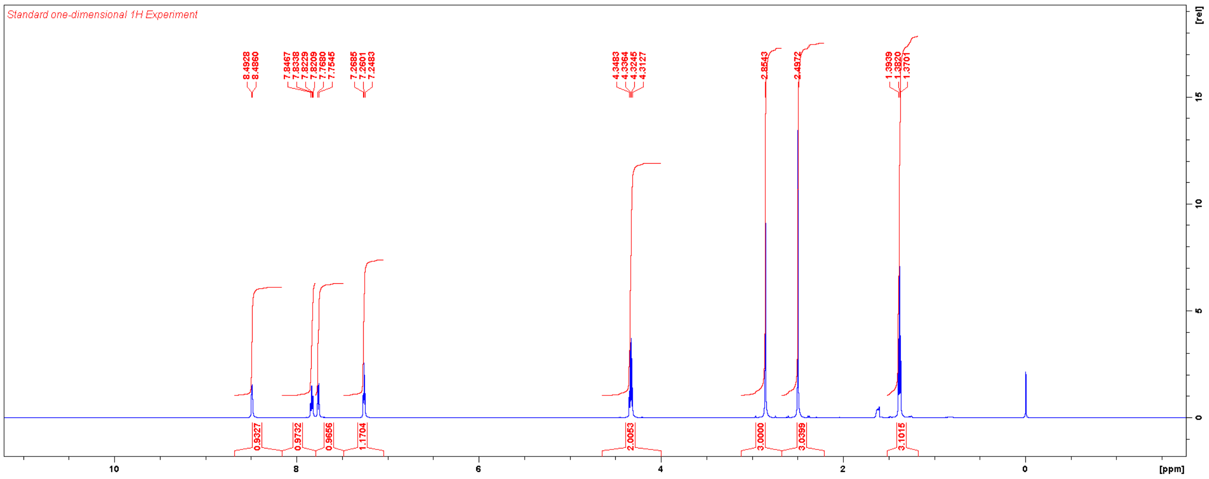

Diethyl (3,5-dimethyl-1-(pyridin-2-yl)-1*H*-pyrazole-4-carbonyl)-*L*-valyl-*D*-glutamate (**TT032**)

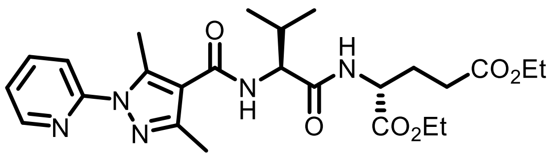


Ethyl 3,5-dimethyl-1-(pyridin-2-yl)-1*H*-pyrazole-4-carboxylate (109 mg, 0.40 mmol) was hydrolysed according to the general procedure for alkaline hydrolysis [Intermediate yield: 76 mg (0.37 mmol, 94%)]. This material was used without further purification to synthesise **TT032** according to the general procedure for HATU-mediated coupling. Yield: 13 mg (0.026 mmol, 43%).

$^{1}$H NMR ($CDCl_3$, 600 MHz) δ 8.51 (m, 1H), 7.86 (m, 2H), 7.32 (m, 1H), 6.74 (br, 1H), 6.32 (br, 1H), 4.58 (m, 2H), 4.20 (m, 2H), 412 (q, 2H, *J* = 7.1 Hz), 2.69–1.95 (m, 11H), 1.27 (m, 6H), 1.06 (d, 3H, *J* = 6.9 Hz), 1.02 (d, 3H, *J* = 6.8 Hz).

$^{13}$C NMR ($CDCl_3$, 150 MHz) δ 172.8, 171.4, 171.2, 164.5, 152.6, 148.4, 147.8, 143.1, 138.5, 122.1, 117.7, 116.2, 61.7, 60.8, 58.1, 52.0, 31.2, 30.3, 27.1, 19.5, 18.0, 14.2, 14.13, 14.09, 13.3.

HRMS m/z: $[M+H]^+$ Calcd for $C_{25}H_{35}N_5O_6$ 502.2660; Found 502.2658.

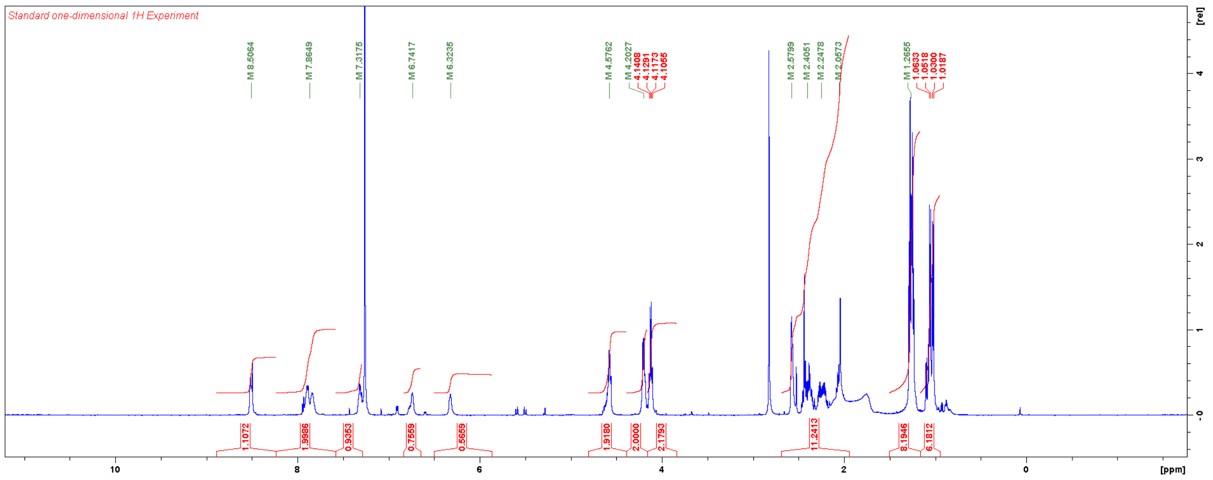


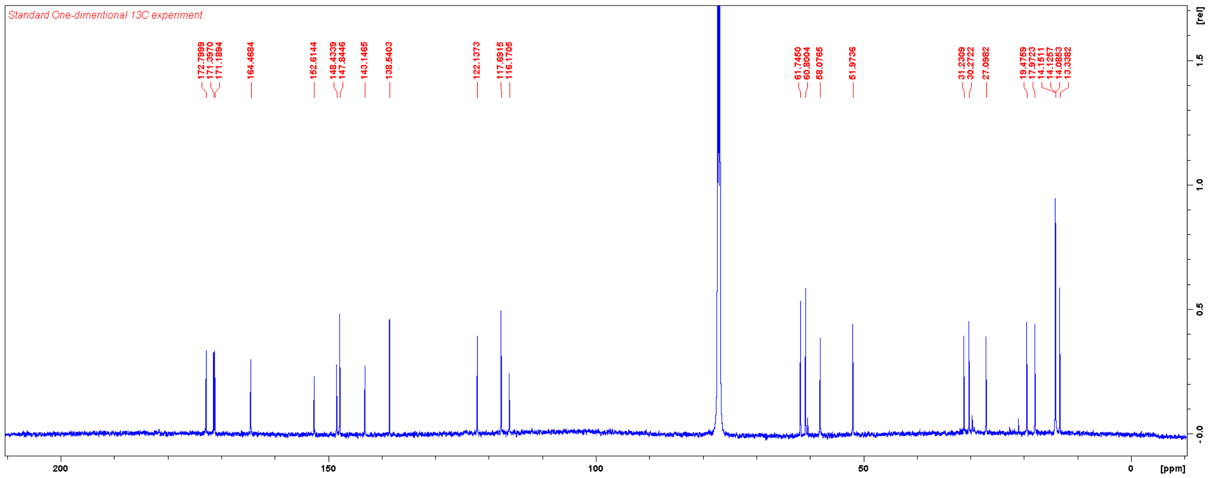

Ethyl 3,5-dimethyl-1-(pyridin-3-yl)-1*H*-pyrazole-4-carboxylate

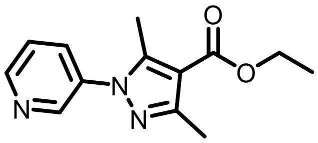

A solution of 3-hydrazinylpyridine dihydrochloride (91 mg, 0.5 mmol), ethyl 2-acetyl-3-oxobutanoate (78 µL, 0.5 mmol), and triethylamine (140 µL, 1.0 mmol) in ethanol (1 mL) was stirred at RT for three days. The solvent was evaporated, and the residue was purified with flash chromatography using a gradient of EtOAc in hexanes. Yield: 25 mg (0.091 mmol, 18%).

$^1$H NMR ($CDCl_3$, 600 MHz) δ 8.70 (m, 2H), 7.80 (d, 1H), 7.46 (m, 1H), 4.34 (q, 2H, *J* = 7.1 Hz), 2.56 (s, 3H), 2.50 (s, 3H), 1.39 (t, 3H, *J* = 7.1 Hz).

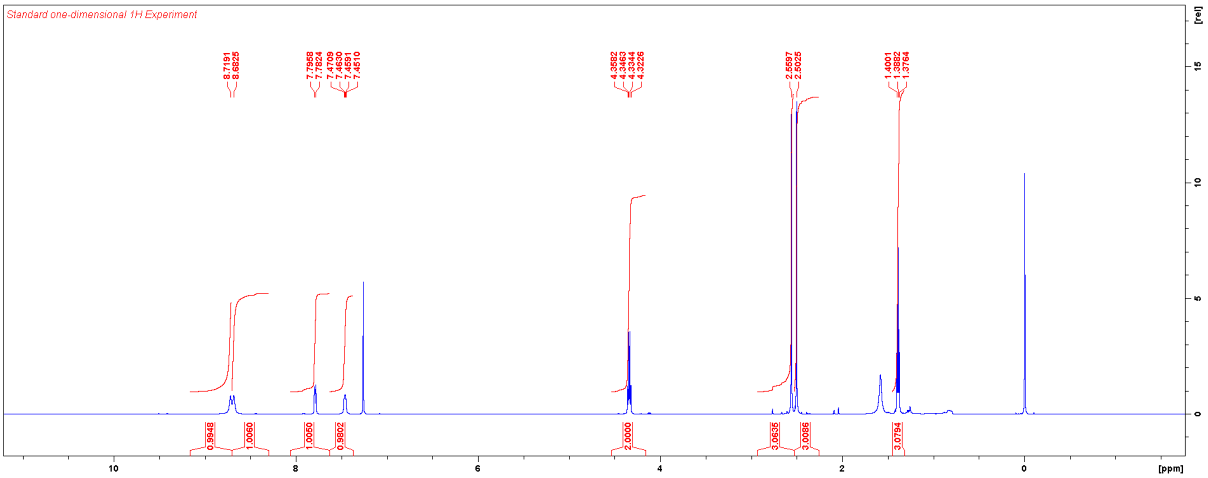

Diethyl (3,5-dimethyl-1-(pyridin-3-yl)-1*H*-pyrazole-4-carbonyl)-*L*-valyl-*D*-glutamate (**TT033**)

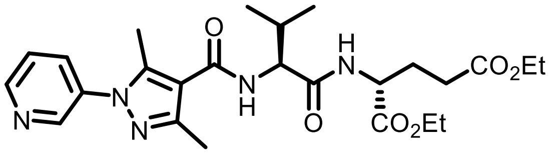


Ethyl 3,5-dimethyl-1-(pyridin-3-yl)-1*H*-pyrazole-4-carboxylate (25 mg, 0.091 mmol) was hydrolysed according to the general procedure for alkaline hydrolysis. Upon completion of the reaction, the aqueous phase was acidified with 1 N HCl to pH ≈ 5 and extracted five times with EtOAc. The combined organic layers were dried over $MgSO_4$, filtered, and concentrated to afford the carboxylic acid derivative [Intermediate yield: 13 mg (0.06 mmol, 66%)]. This material was used without further purification to synthesise **TT033** according to the general procedure for HATU-mediated coupling. Yield: 16 mg (0.032 mmol, 53%).

$^1$H NMR ($CDCl_3$, 600 MHz) δ 8.88 (br, 1H), 8.71 (br, 1H), 8.16 (br, 1H), 7.73 (br, 1H), 6.82 (d, 1H, *J* = 7.3 Hz), 6.37 (d, 1H, *J* = 8.2 Hz), 4.59 (m, 2H), 4.20 (m, 2H), 4.13 (q, 2H, 6.8 Hz), 2,61 (s, 3H), 2.53 (s, 3H), 2.42 (m, 2H), 2.24 (m, 3H), 2.06 (m, 1H), 1.27 (m, 6H), 1.06 (d, 3H, *J* = 6.8 Hz), 1.02 (d, 3H, *J* = 6.8 Hz).

$^{13}$C NMR ($CDCl_3$, 150 MHz) δ 172.8, 171.4, 171.2, 164.1, 148.7, 148.0, 145.0, 142.7, 133.6, 129.5, 124.3, 115.6, 61.8, 60.8, 58.0, 52.0, 31.4, 30.3, 27.1, 19.5, 18.0, 14.2, 14.1, 14.0, 12.3.

HRMS m/z: $[M+H]^+$ Calcd for $C_{25}H_{35}N_5O_6$ 502.2660; Found 502.2657.

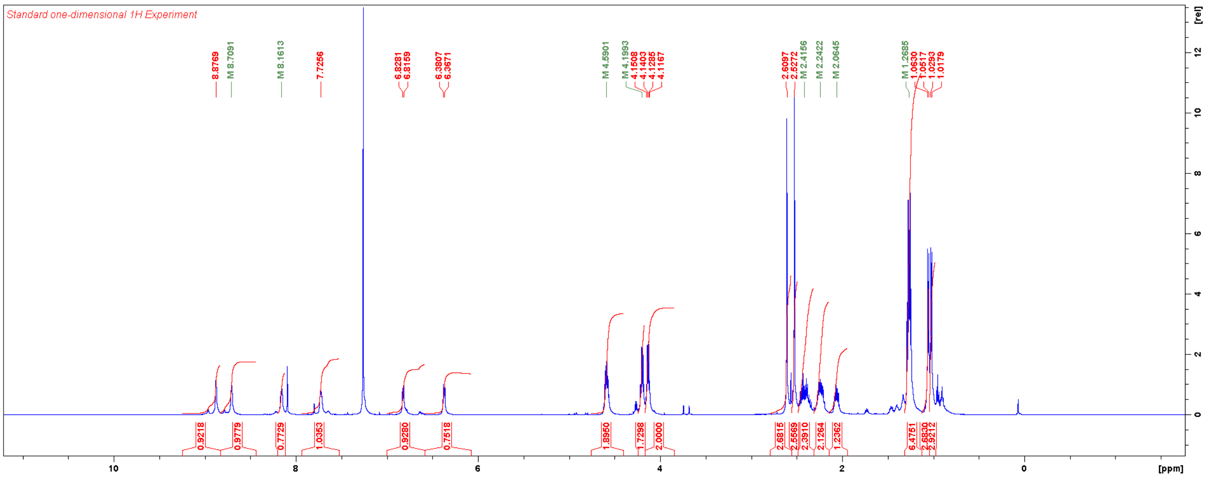


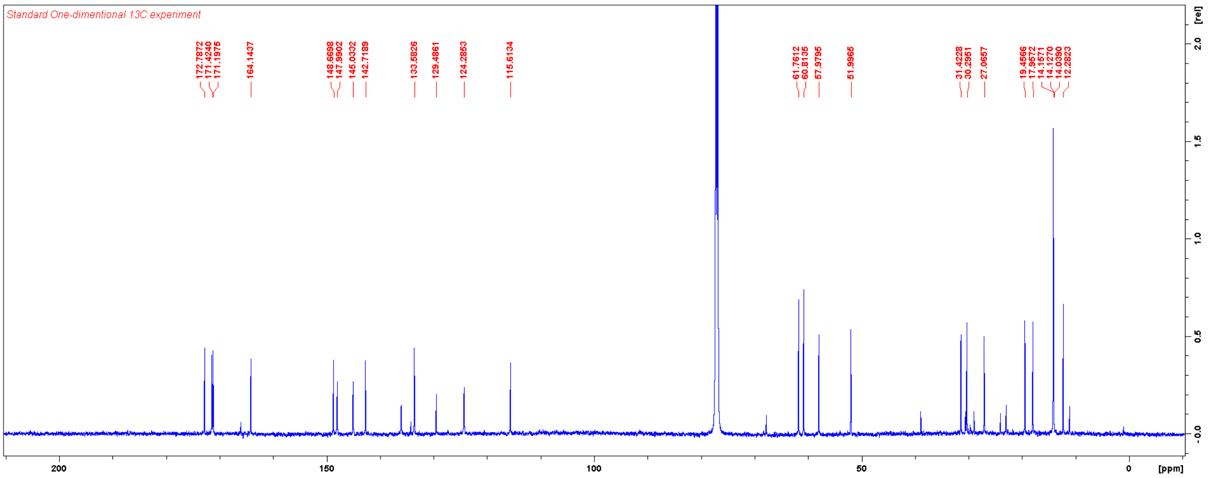

Ethyl 3,5-dimethyl-1-(pyrimidin-5-yl)-1*H*-pyrazole-4-carboxylate

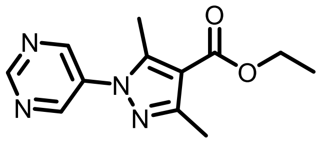


Pyrimidin-5-ylboronic acid (62 mg, 0.50 mmol) and ethyl 3,5-dimethyl-1H-pyrazole-4-carboxylate (84 mg, 0.50 mmol) were weighed in a scintillation vial and dissolved in DMF (2 mL) under ambient atmosphere. To this $Cu(OAc)_2$ (68 mg, 0.38 mmol) and pyridine (81 µL, 1.0 mmol) were added. The reaction mixture was stirred at RT for three days and then diluted with 75% EtoAc in hexanes and washed with brine. Flash chromatography using a gradient of EtOAc in hexanes afforded the title compound. Yield: 24 mg (0.097 mmol, 19%).

$^1$H NMR ($CDCl_3$, 600 MHz) δ 9.22 (s, 1H), 8.88 (m, 2H), 4.32 (q, 2H, *J* = 7.1 Hz), 2.59 (s, 3H), 2.48 (s, 3H), 1.37 (t, 3H, *J* = 7.1 Hz).

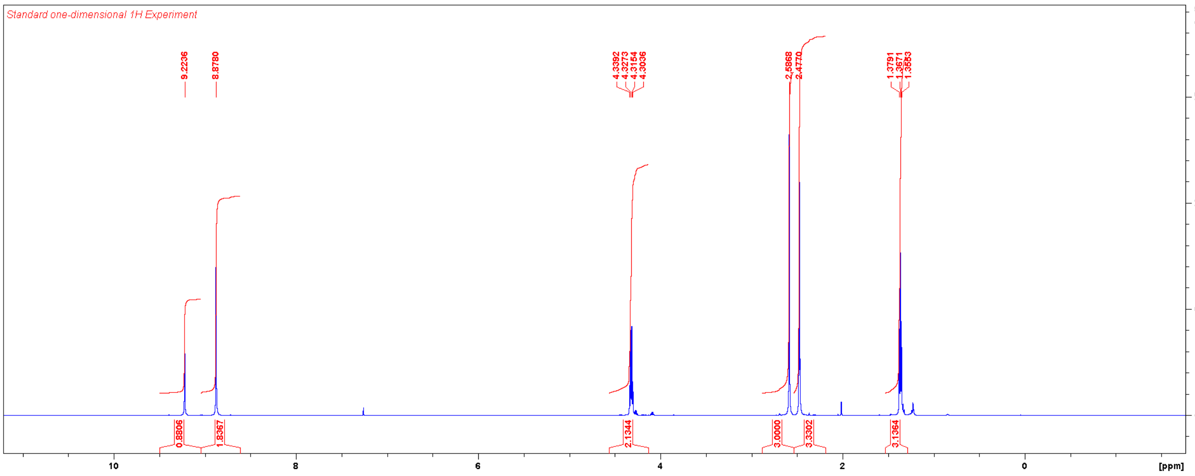

Diethyl (3,5-dimethyl-1-(pyrimidin-5-yl)-1*H*-pyrazole-4-carbonyl)-*L*-valyl-*D*-glutamate (**TT034**)

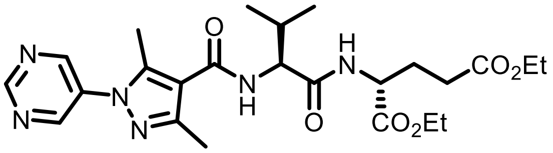

Ethyl 3,5-dimethyl-1-(pyrimidin-5-yl)-1*H*-pyrazole-4-carboxylate (24 mg, 0.097 mmol) was hydrolysed according to the general procedure for alkaline hydrolysis [Intermediate yield: 21 mg (0.10 mmol, quant.)]. This material was used without further purification to synthesise **TT034** according to the general procedure for HATU-mediated coupling. Yield: 19 mg (0.038 mmol, 38%).

$^{1}$H NMR ($CDCl_3$, 600 MHz) δ 9.26 (s, 1H), 8.93 (m, 2H), 6.82 (br, 1H), 6.43 (br, 1H), 4.58 (m, 2H), 4.21 (m, 2H), 4.14 (m, 2H), 2.59 (s, 3H), 2.54 (s, 3H), 2.41 (m, 2H), 2.24 (m, 2H), 2.07 (m, 1H), 1.27 (m, 6H), 1.06 (d, 3H, *J* = 6.7 Hz), 1.03 (d, 3H, *J* = 6.8 Hz).

$^{13}$C NMR ($CDCl_3$, 150 MHz) δ 172.8, 171.4, 171.2, 163.9, 157.1, 152.6, 149.4, 143.0, 134.3, 116.2, 61.8, 60.9, 58.1, 52.1, 31.5, 30.3, 27.0, 19.4, 18.0, 14.2, 14.1, 14.0, 12.2.

HRMS m/z: $[M+H]^+$ Calcd for $C_{24}H_{34}N_6O_6$ 503.2613; Found 503.2611.

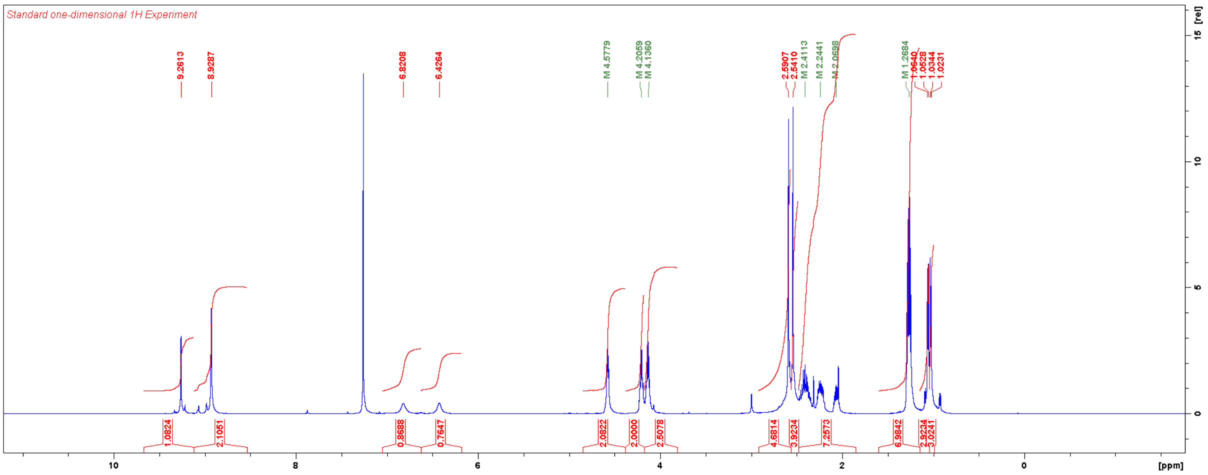


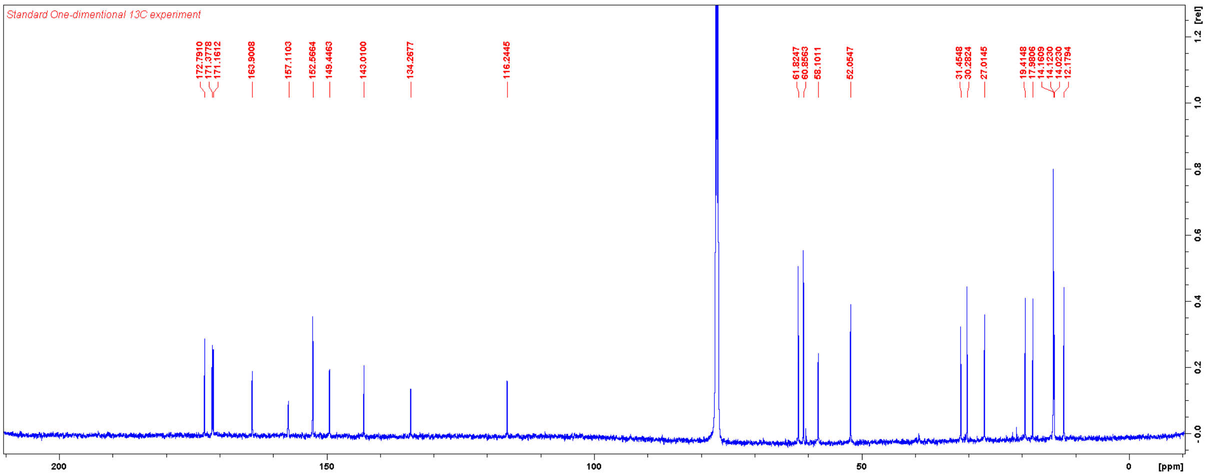

Ethyl 1-(2-methoxyphenyl)-3,5-dimethyl-1*H*-pyrazole-4-carboxylate

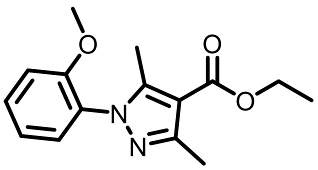

A solution of 2-methoxyphenylhydrazine hydrochloride (349 mg, 2.00 mmol) and ethyl 2-acetyl-3-oxobutanoate (312 µL, 2.00 mmol) in ethanol (5 mL) was refluxed for 3 h. The solvent was evaporated, and the residue was purified with column chromatography eluting with 15% EtOAc/hexanes. Yield: 152 mg (0.55 mmol, 28%).

$^1$H NMR ($CDCl_3$, 600 MHz) δ 7.43 (m, 1H), 7.31 (dd, 1H, *J* = 7.7, 1.4 Hz), 7.04 (m, 2H), 4.32 (q, 2H, *J* = 7.1 Hz), 3.80 (s, 3H), 2.50 (s, 3H), 2.33 (s, 3H), 1.38 (t, 3H, *J* = 7.1 Hz).

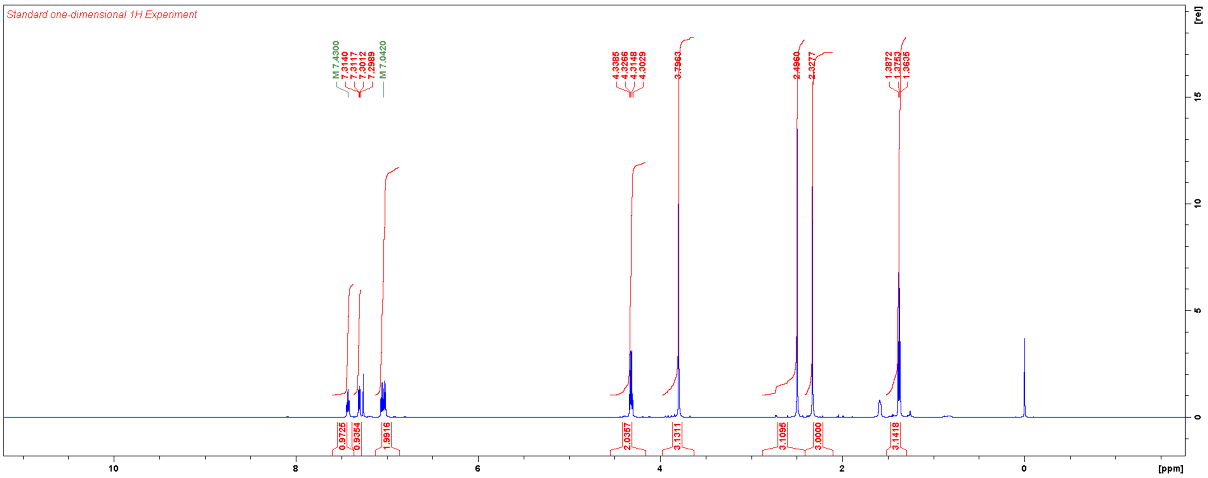

Diethyl (1-(2-methoxyphenyl)-3,5-dimethyl-1*H*-pyrazole-4-carbonyl)-*L*-valyl-*D*-glutamate (**TT035**)

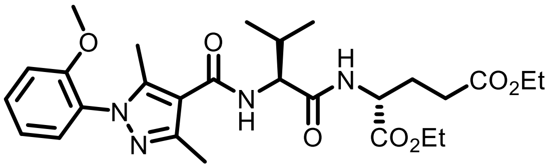

Ethyl 1-(2-methoxyphenyl)-3,5-dimethyl-1*H*-pyrazole-4-carboxylate (58 mg, 0.21 mmol) was hydrolysed according to the general procedure for alkaline hydrolysis [Intermediate yield: 45 mg (0.18 mmol, 87%)]. This material was used without further purification to synthesise **TT035** according to the general procedure for HATU-mediated coupling. Yield: 29 mg (0.054 mmol, 30%).

$^{1}$H NMR ($CDCl_3$, 600 MHz) δ 7.48 (m, 1H), 7.34 (m, 1H), 7.07 (m, 2H), 6.73 (br, 1H), 6.35 (br, 1H), 4,58 (m, 2H), 4.20 (m, 2H), 4.13 (m, 2H), 3.83 (s, 3H), 2.81–1.89 (m, 11H), 1.27 (m, 6H), 1.06 (d, 3H, *J* = 6.8 Hz), 1.03 (d, 3H, *J* = 6.8 Hz).

$^{13}$C NMR ($CDCl_3$, 150 MHz) δ 172.8, 171.43, 171.35, 164.7, 154.6, 147.5, 144.4, 130.8, 129.1, 127.3, 120.9, 113.3, 111.9, 61.7, 60.8, 58.0, 55.8, 52.0, 31.3, 30.3, 27.1, 19.5, 18.0, 14.3, 14.2, 14.1, 11.6.

HRMS m/z: $[M+H]^+$ Calcd for $C_{27}H_{38}N_4O_7$ 531.2813; Found 531.2810.

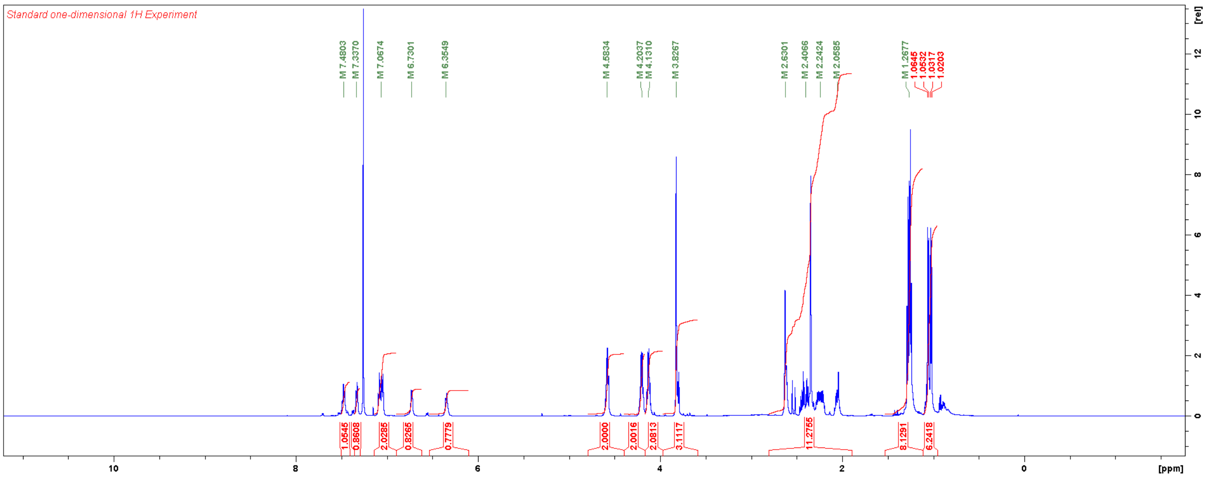


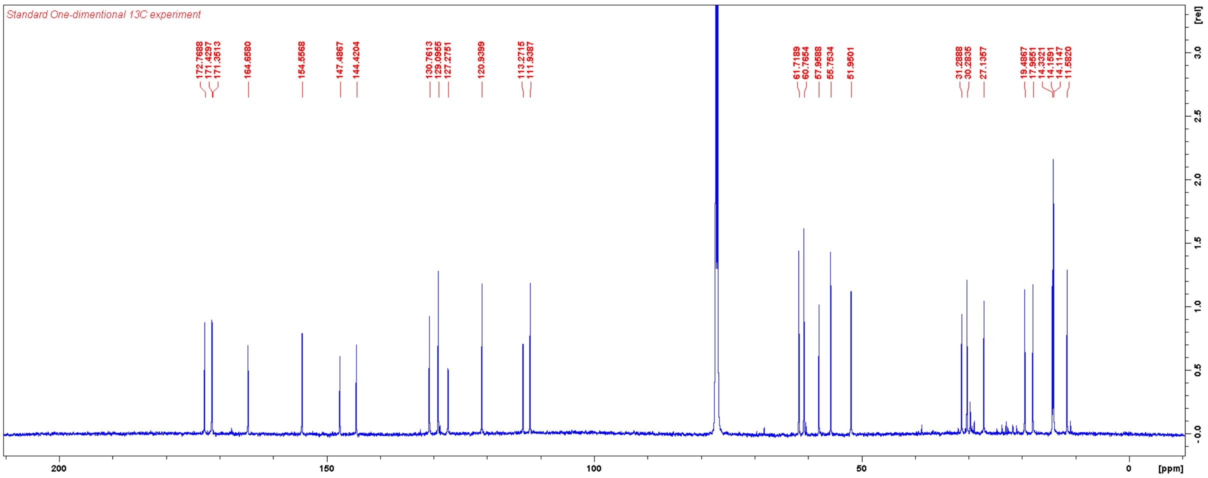

Ethyl 1-(3-methoxyphenyl)-3,5-dimethyl-1*H*-pyrazole-4-carboxylate

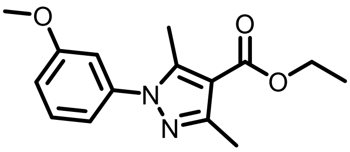

A solution of 3-methoxyphenylhydrazine hydrochloride (87 mg, 0.5 mmol) and ethyl 2-acetyl-3-oxobutanoate (78 µL, 0.5 mmol) in ethanol (1 mL) was stirred at RT overnight. The solvent was evaporated, and the residue was purified with flash chromatography using a gradient of EtOAc in hexanes. Yield: 26 mg (0.094 mmol, 19%).

$^1$H NMR ($CDCl_3$, 600 MHz) δ 7.37 (m, 1H), 6.95 (m, 3H), 4.33 (q, 2H, *J* = 7.1 Hz), 3.84 (s, 3H), 2.52 (s, 3H), 2.49 (s, 3H), 1.38 (t, 3H, *J* = 7.1 Hz).

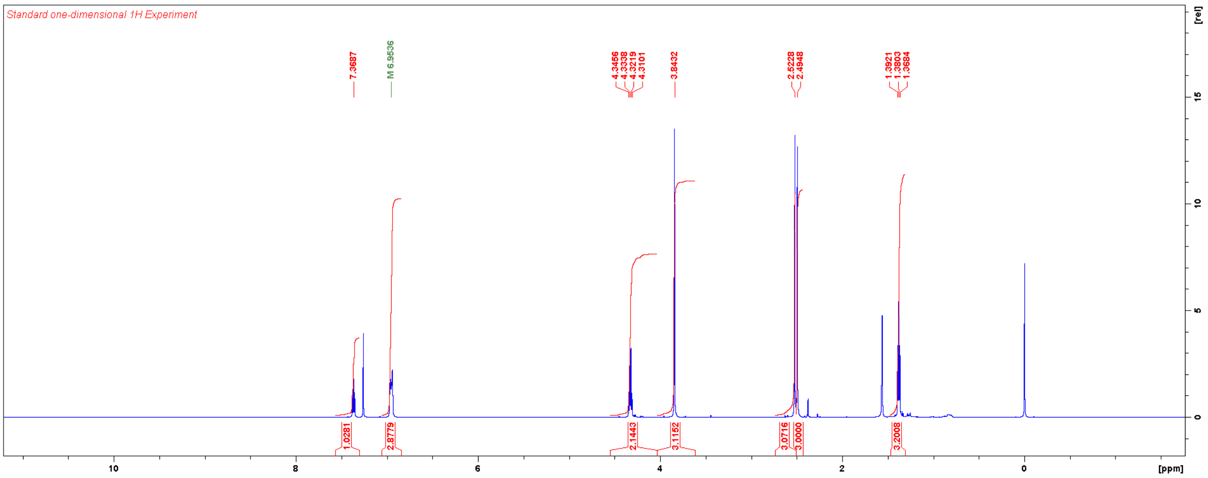

Diethyl (1-(3-methoxyphenyl)-3,5-dimethyl-1*H*-pyrazole-4-carbonyl)-*L*-valyl-*D*-glutamate (**TT036**)

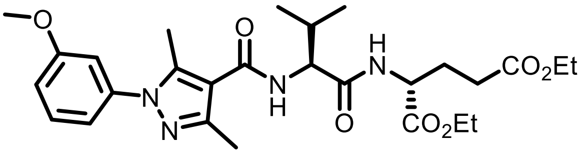

Ethyl 1-(3-methoxyphenyl)-3,5-dimethyl-1*H*-pyrazole-4-carboxylate (50 mg, 0.18 mmol) was hydrolysed according to the general procedure for alkaline hydrolysis [Intermediate yield: 33 mg (0.13 mmol, 71%)]. This material was used without further purification to synthesise **TT036** according to the general procedure for HATU-mediated coupling. Yield: 22 mg (0.041 mmol, 32%).

$^{1}$H NMR ($CDCl_3$, 600 MHz) δ 7.39 (m, 1H), 6.98 (m, 3H), 6.74 (d, 1H, *J* = 6.9 Hz), 6.34 (d, 1H, *J* = 7.7 Hz), 4.58 (m, 2H), 4.20 (m, 2H), 4.13 (m, 2H), 3.86 (s, 3H), 2.60 (s, 3H), 2.52 (s, 3H), 2.41 (m, 2H), 2.25 (m, 2H), 2.06 (m, 1H), 1.27 (m, 6H), 1.06 (d, 3H, *J* = 6.8 Hz), 1.02 (d, 3H, *J* = 6.8 Hz).

$^{13}$C NMR ($CDCl_3$, 150 MHz) δ 172.8, 171.4, 171.3, 164.5, 160.2, 147.3, 142.5, 139.7, 129.9, 117.8, 114.5, 114.4, 111.3, 61.7, 60.8, 58.0, 55.5, 52.0, 31.3, 30.3, 27.1, 19.5, 17.9, 14.2, 14.1, 12.4.

HRMS m/z: $[M+H]^+$ Calcd for $C_{27}H_{38}N_4O_7$ 531.2813; Found 531.2810.

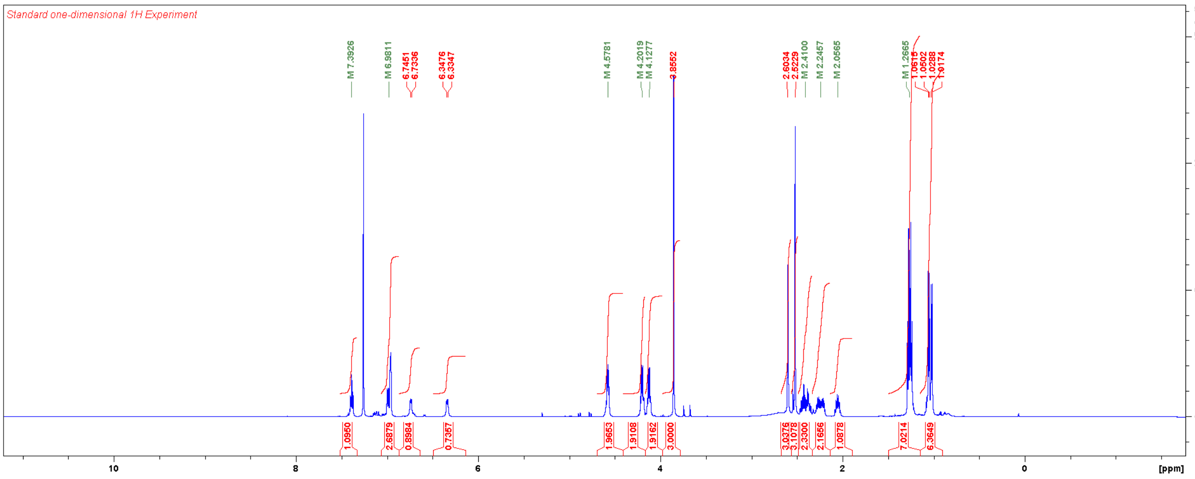


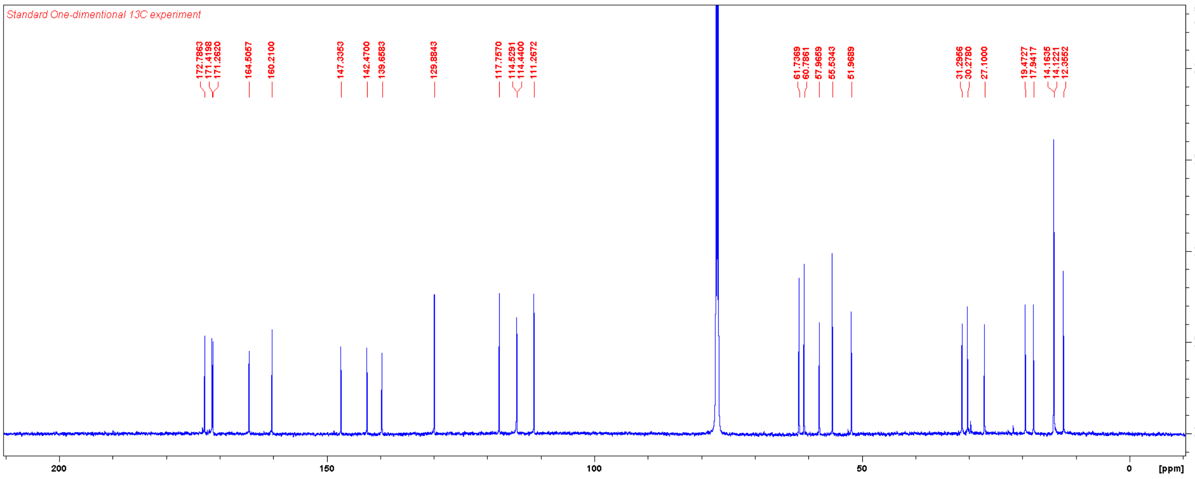

Diethyl (tert-butoxycarbonyl)-*D*-valyl-*L*-glutamate

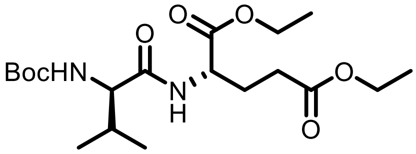

To a solution of Boc-*D*-valine-OH (227 mg, 1.04 mmol), diethyl *L*-glutamate hydrochloride (250 mg, 1.04 mmol), EDC-HCl (200 mg, 1.04 mmol) and HOBt-x$H_2O$ (155 mg, 1.15 mmol) in DMF (5 mL) was added TEA (291 µL, 2.09 mmol) at 0 °C. The reaction mixture was allowed to warm to RT overnight. The reaction mixture was quenched with aqueous $NH_4Cl$ and extracted three times with 50% EtOAc in hexanes. Combined organic layers was washed with brine, dried over $Na_2SO_4$, filtered, concentrated, dried under vacuum for three days. This material was used in proceeding reactions without further purification. Yield: 416 mg (1.03 mmol, 99%), white powder.

$^1$H NMR ($CDCl_3$, 600 MHz) δ 6.69 (d, 1H, *J* = 7.0 Hz), 4.97 (m, 1H), 4.58 (dd, 1H, *J* = 12.8, 7.5 Hz), 4.20 (q, 2H, *J* = 7.1 Hz), 4.13 (q, 2H, *J* = 7.1 Hz), 4.00 (m 1H), 2.38 (m, 2H), 2.21 (m, 2H), 2.01 (m, 1H), 1.45 (s, 9H), 1.28 (t, 3H, *J* = 7.1 Hz), 1.25 (t, 3H, *J* = 7.1 Hz), 0.98 (d, 3H, *J* = 6.7 Hz), 0.91 (d, 3H, *J* = 6.8 Hz).

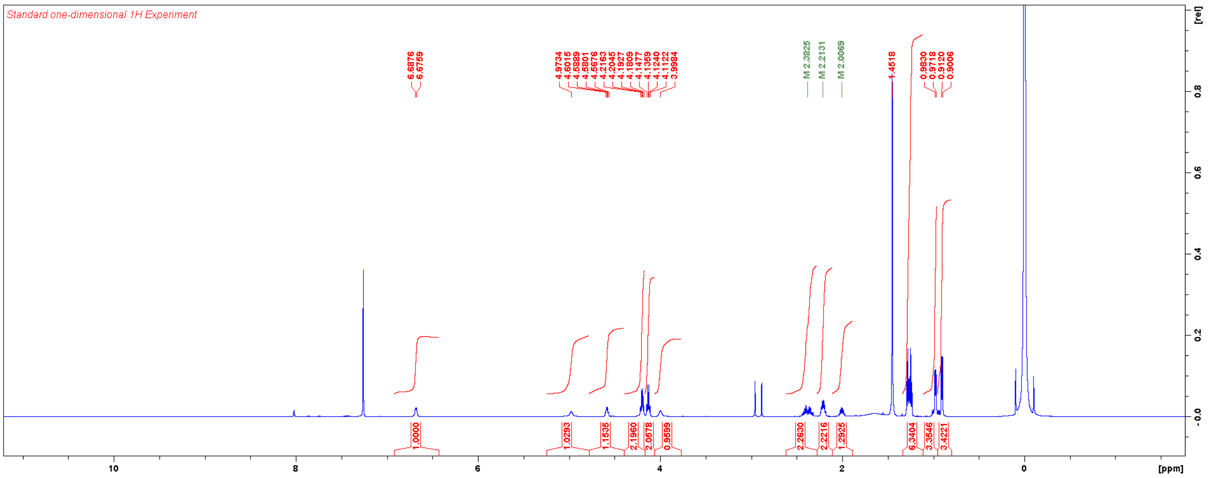

Diethyl (3,5-dimethyl-1-(2-oxo-1,2-dihydropyridin-4-yl)-1*H*-pyrazole-4-carbonyl)-*D*-valyl-*L*-glutamate (**TT030, DL**)

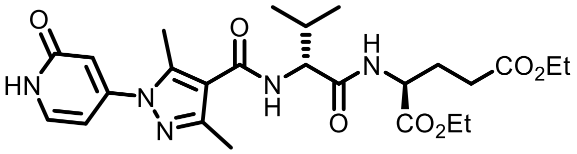

**TT030, DL** was synthesised according to the general procedure for HATU-mediated coupling. Upon completion of the reaction, reaction mixture was diluted with EtOAc, washed with aqueous $NH_4Cl$, dried over $Na_2SO_4$, filtered, and concentrated. The residue was purified with column chromatography using two different columns and eluents: first with 5% MeOH/DCM; then on a new column with 100% THF. Yield: 9 mg (0.017 mmol, 16%).

$^1$H NMR ($CDCl_3$, 600 MHz) δ 7.41 (d, 1H, *J* = 7.1 Hz), 6.94 (br, 1H), 6.63 (m, 2H), 6.51 (d, 1H, *J* = 7.1 Hz), 4.57 (m, 2H), 4.20 (m, 2H), 4.13 (q, 2H, *J* = 7.1 Hz), 2.62 (s, 3H), 2.48 (s, 3H), 2.41 (m, 2H), 2.25 (m, 2H), 2.07 (m, 1H), 1.27 (m, 6H), 1.06 (d, 3H, *J* = 6.8 Hz), 1.03 (d, 3H, *J* = 6.8 Hz).

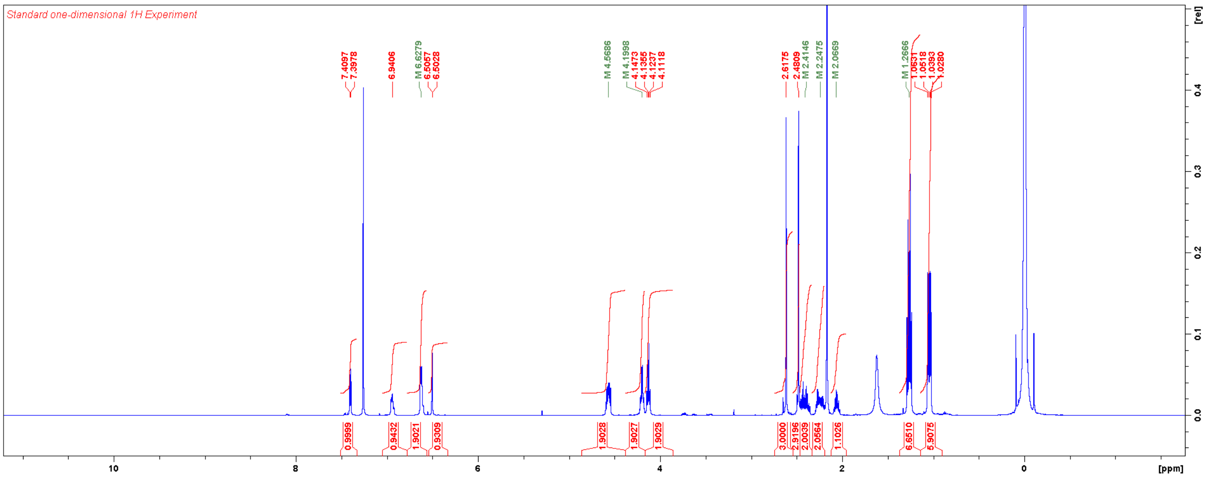
